# A population-scale atlas of blood and tissue in lupus nephritis

**DOI:** 10.1101/2025.08.11.669754

**Authors:** Nicholas W. Sugiarto, Siddarth Gurajala, Michelle Curtis, Thomas M. Eisenhaure, Arnon Arazi, Andrea Fava, Qian Xiao, Joseph Mears, Brad Rovin, Celine C. Berthier, Yu Zhao, Peter M. Izmirly, Jennifer L. Barnas, Paul J. Hoover, Michael Peters, Raktima Raychowdhury, Alice Horisberger, Saori Sakaue, Richard A. Furie, H. Michael Belmont, David A. Hildeman, E. Steve Woodle, Maria Dall’Era, Chaim Putterman, Diane L. Kamen, Maureen A. McMahon, Jennifer Grossman, Kenneth C. Kalunian, Jeffrey B. Hodgin, Fernanda Payan-Schober, William Apruzzese, Harris Perlman, Carla M. Cuda, David Wofsy, Joel M. Guthridge, Jennifer H. Anolik, Judith A. James, Accelerating Medicines Partnerships Rheumatoid Arthritis/Systemic Lupus Erythematosus, Deepak A. Rao, Anne Davidson, Michelle A. Petri, Jill P. Buyon, Nir Hacohen, Betty Diamond, Soumya Raychaudhuri

**Affiliations:** Center for Data Sciences, Brigham and Women’s Hospital and Harvard Medical School, Boston, MA 02115, USA; Division of Rheumatology, Inflammation, and Immunity, Department of Medicine, Brigham and Women’s Hospital and Harvard Medical School, Boston, MA 02115, USA; Broad Institute of MIT and Harvard, Cambridge, MA 02142, USA; Division of Genetics, Department of Medicine, Brigham and Women’s Hospital and Harvard Medical School, Boston, MA 02115, USA; Center for Autoimmune, Musculoskeletal and Hematopoietic Diseases, The Feinstein Institute for Medical Research, Northwell Health, Manhasset, NY, USA; Division of Rheumatology, Johns Hopkins University, Baltimore, MD, USA; Ohio State University Wexner Medical Center, Columbus, OH, 43210, USA; Internal Medicine, Department of Nephrology, University of Michigan, Ann Arbor, MI, USA; Department of Medicine, Division of Rheumatology, New York University School of Medicine, New York, NY, USA; Department of Medicine, Division of Allergy, Immunology, and Rheumatology, University of Rochester Medical Center, Rochester, NY, USA; Division of Rheumatology, Northwell Health, Great Neck, NY, USA; Department of Pediatrics, University of Cincinnati, Cincinnati, OH, USA; Division of Immunobiology, Cincinnati Children’s Hospital Medical Center, Cincinnati, Ohio, USA; Division of Transplantation, Department of Surgery, University of Cincinnati College of Medicine, Cincinnati, OH, USA; Rheumatology Division, University of California San Francisco, San Francisco, CA, USA; Division of Rheumatology and Department of Microbiology and Immunology, Albert Einstein College of Medicine and Montefiore Medical Center, Bronx, NY, USA; Division of Rheumatology and Immunology, Medical University of South Carolina, Charleston, SC, USA; Department of Medicine, University of California Los Angeles, Los Angeles, CA, USA; School of Medicine, University of California San Diego, La Jolla, California, USA; Department of Pathology, University of Michigan, Ann Arbor, MI, USA; Department of Medicine, Paul L. Foster School of Medicine, Texas Tech University Health Sciences Center, El Paso, TX, USA; Pfizer Inc., New York, NY, USA; Division of Rheumatology, Department of Medicine, Northwestern University Feinberg School of Medicine, Chicago, IL, USA; Arthritis and Clinical Immunology, Oklahoma Medical Research Foundation, Oklahoma City, OK, USA; Department of Biomedical Informatics, Harvard Medical School, Boston, MA 02115, USA; Division of Rheumatology, The University of Colorado Anschutz Medical Campus, Aurora, CO, USA; Kao Autoimmunity Institute, Cedars-Sinai Medical Center, Los Angeles, California; Laboratory for RNA Molecular Biology, The Rockefeller University, 1230 York Ave, Box 186, New York, NY 10065, USA; Stanford University School of Medicine, Stanford, California

**Author notes:** These authors contributed equally (Nicholas W. Sugiarto, Siddarth Gurajala, Michelle Curtis, Thomas M. Eisenhaure). Co-senior, co-corresponding authorship (Betty Diamond, Soumya Raychaudhuri).

## Abstract

**One Sentence Summary:** A single-cell atlas of paired blood and tissue samples from Lupus Nephritis patients and healthy controls identified stromal and immune populations within renal tissue, including the scar-associated macrophage populations, which correlate with and may drive renal disease activity.

Lupus nephritis (LN), a severe manifestation of Systemic Lupus Erythematosus (SLE), is a heterogeneous disease driven by diverse immune and tissue cell types. We obtained 538K single-cell and 140K single-nuclear profiles from kidney biopsies of 155 LN patients and 30 pre-implantation transplant biopsy controls, along with 325K single-cell blood profiles overlapping many of these patients. We identified key tissue cell types and cell states, and immune cell states; we were able to determine cell states that were tissue specific, and those that were present in the blood. We observed that LN pathological features are significantly associated with cell states using differential gene expression and Covarying Neighborhood Analysis (CNA). These analyses revealed broad changes in cell states associated with irreversible chronic tissue damage. After controlling for the effects of ongoing tissue damage, we observed that expansion of key glomerular and Scar Associated Macrophages (SAMs) populations tracked with increasing inflammatory disease activity. SAMs appear to drive LN fibrosis and, in active disease, infiltrate the glomeruli more than other myeloid cells. These observations strongly support that therapeutic targeting of myeloid populations may offer an as-of-yet unproven strategy to prevent renal inflammation and ongoing kidney damage in LN.

## Introduction

Systemic lupus erythematosus (SLE) is a chronic autoimmune disease characterized by autoantibodies against nuclear components and heterogenous clinical manifestations across multiple organ systems(*1*). Affecting 0.1% of the population, SLE predominantly affects female and non-white individuals(*2*). Lupus nephritis (LN) presents in 30-50% of patients and results from inflammation-induced kidney damage(*3*, *4*). Immunosuppressive therapies including prednisone, mycophenolate mofetil, cyclophosphamide, rituximab, belimumab, and voclosporin, are inconsistently effective and can increase infection risk. Fewer than 50% of individuals achieve complete renal response(*5*), and relapse is common(*6*). Approximately 15% progress to end-stage renal disease (ESRD) despite therapy(*7*) resulting in morbidity and mortality(*8*, *9*).

Kidney biopsy remains essential for LN diagnosis and treatment despite efforts to identify serum biomarkers. The International Society of Nephrology (ISN) classifies LN into six classes. Class I-IV glomerular pathology denotes increasing proliferative disease. Class V designates membranous disease, alone or in conjunction with proliferative disease. The rare Class VI reflects severe scarring with globally sclerotic glomeruli. Clinicians treat class III, IV, and V aggressively, although emerging data support treating less advanced disease forms as well(*10*, *11*). Renal biopsies are also scored using the NIH activity and chronicity indices(*12*, *13*). The activity index captures active and potentially reversible inflammation, but high sustained scores predict poor survival(*14*). It reflects the accumulation of immune cells in the glomerular capillaries (glomerular endocapillary hypercellularity), neutrophilic infiltration, fibrinoid necrosis, large subendothelial immune complexes (hyaline deposits), cellular and fibrocellular crescents, and cortical tubulointerstitial inflammation(*13*). The chronicity index captures irreversible tissue damage, which increases with each LN flare(*15*) and correlates with reduced kidney survival. It includes glomerulosclerosis, fibrous crescents, tubular atrophy, and interstitial fibrosis. While imperfect predictors, biopsy characteristics outperform other clinical metrics. Indicators such as low serum albumin and persistent proteinuria (>800mg/g creatinine) suggest poor long-term prognosis(*16–18*), but may not be concordant with the underlying histology(*19*). Thus, clinicians variably place weight on ISN class, activity, and chronicity in therapeutic decisions.

There has been great interest in using high-dimensional genomics to understand LN. The largest studies have been in bulk PBMCs and have confirmed previously identified Interferon Stimulated Gene (ISG) signatures upregulated in LN cases(*20*). Single-cell data have allowed us to localize these interferon signals to particular cell types by identifying the myeloid compartment as the largest expressor of ISGs(*21*). Recent efforts to integrate multiple public PBMC LN datasets have also pinpointed *GZMK*^+^*GZMH*^+^*HLA-DR*^+^ effector memory CD8+ T cells as a major expressor of pro-inflammatory cytokines(*22*). However, renal tissue investigations of LN may be particularly valuable. Thus far, these primary tissue studies have been limited in sample size(*23*) or confined to histopathology(*24*). Spatial datasets of LN tissue are now emerging(*25*, *26*). While these studies have led to valuable insights, they are currently limited by the number of samples profiled. Furthermore, these atlases are limited by the number of genes they can incorporate into their panels and therefore often require external single-cell datasets to accurately call cell types. In many instances, they lack the necessary number of transcripts in single cell profiles to resolve specific cell states and their associations to LN subphenotypes.

The goal of the Accelerating Medicines Partnership in SLE is to profile single cells from human patient tissues, such as the nephritic kidney, to advance understanding of LN pathology at cellular resolution(*16*, *27*). We applied single-cell (scRNAseq) and single-nuclear (snRNAseq) RNA-sequencing to profile both immune and non-immune cell populations. These methods recovered different proportions of cell types, as non-immune cells were more sensitive to the enzymatic disruption of cryopreserved tissue used to generate scRNAseq data. With these data, we defined key tissue-specific cell states and identified those linked to case/control status, chronicity, activity, and other SLE parameters. We observed that the chronicity and activity better explain effects on cell states than ISN class.

## Results

### Cell State Definition and Clustering

We obtained kidney biopsies and peripheral blood mononuclear cell (PBMC) samples from patients with LN and healthy controls. Our cohort was 87.0% female in case samples and 76.7% female in control samples **(Table S1, Table S2C)**. 32.9% of case samples and 20.0% of control samples had a prior history of biopsy prior to enrollment. Each donor was only sampled once in our cohort. After quality control, the final dataset included tissue and clinical data from 155 LN patients and 30 healthy controls, and matched PBMC samples from 118 of the LN patients and 19 healthy controls (**Fig. 1A, Table S2A**). Following quality control to remove poor-quality cells based on low UMI counts, low gene features, and high mitochondrial reads **(Table S3, Fig. S1A-C)**, the kidney tissue data included 538,194 scRNA-seq profiles and 142,881 snRNA-seq profiles from a subset of 50 patients, the latter preserving disaggregation-sensitive cell types. Per-biopsy cell yields were variable across donors with 95.7% of scRNAseq donors and 95.9% of snRNAseq donors contributing >500 cells **(Fig. S2B).** We applied weighted principal components analysis(*28*) followed by Harmony(*29*) to ensure both modalities contributed equally. Clustering and annotating cells together ensured that shared transcriptional signals represented biology and not modality-specific technical artifacts. (**Fig. 1B**). The PBMC data included 327,326 scRNA-seq profiles, which we integrated with Harmony. We also integrated PBMC and kidney tissue data to identify tissue-specific cell states.

**Fig. 1.**
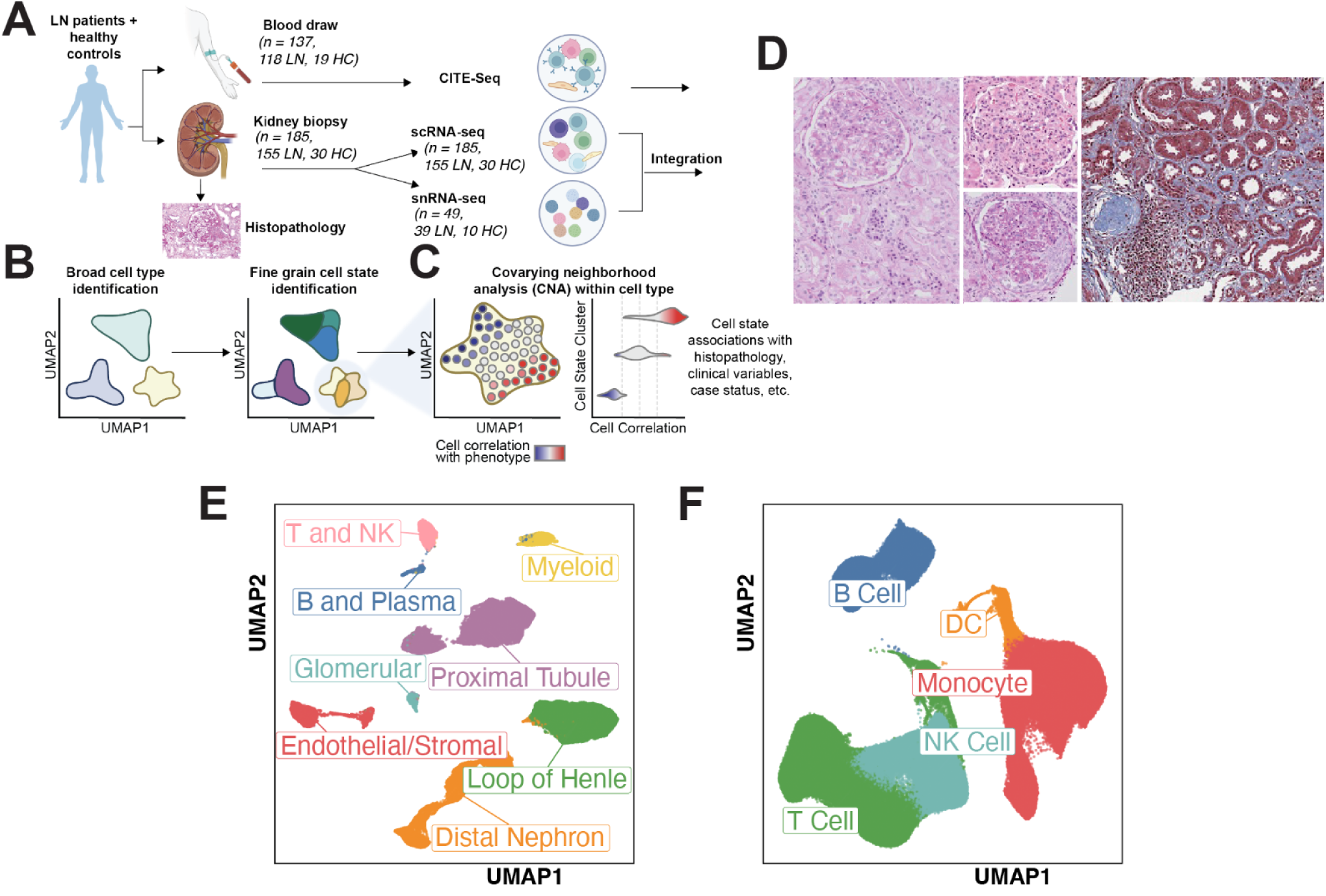
Dataset overview. **A-C,** Schematic overview of the data collection and analysis pipeline. (**A**) Blood and kidney samples were collected from LN patients and healthy controls (HC). Histopathology and clinical scoring was performed on renal biopsies. CITE-seq was performed on blood samples, and scRNA-seq and snRNA-seq were performed on tissue samples. Both tissue modalities were integrated with Harmony^24^. (**B**) Broad cell types were annotated separately in tissue and blood. Fine grain clustering for cell state annotation was performed on each broad cell type. (**C**) For each tissue, we performed CNA within each cell type to define cell state associations with various clinical variables. **(D)** Left: Normal glomerulus and tubulointerstitium (PAS stain). An H&E image of the same region is preferred and will be obtained. Middle: Two glomeruli showing active lesions. Top: H&E stain demonstrating global endocapillary hypercellularity. Bottom: PAS stain showing a cellular crescent. Right: Low-power trichrome stain highlighting chronic/fibrotic changes in blue, including diffuse interstitial fibrosis, tubular atrophy, and a globally sclerotic glomerulus. **(E-F),** UMAPs of all single cell or single nuclear annotations in tissue (**E**) and blood (**F**), colored by broad cell type annotation.

We associated cell states with case/control status and histopathological metrics using Covarying Neighborhood Analysis (CNA)(*30*) (**Fig. 1C, D**). Rather than relying on discrete clusters, CNA overcomes multiple hypothesis testing power limitations by examining groups of similar individual cells, known as neighborhoods, and identifying those significantly associated with a sample-level variable of interest. We interpret the associated neighborhoods within the context of annotated clusters.

Clustering transcriptionally similar cells in renal tissue revealed 3 immune (T/NK, B/Plasma, Myeloid) and 5 renal tissue (glomeruli, proximal tubule, loop of Henle, distal nephron, endothelial/stromal) populations (**Fig. 1E, F**), with consistent marker genes across integrated scRNAseq and snRNAseq data **(Fig. S3-10)**. The immune compartment included 71,859 single-cells and 4,652 single-nuclei. The tissue compartment included 466,335 single-cells and 138,229 single-nuclei. Prior studies show related immune cell enrichment in scRNAseq data, likely due to destruction of tissue cells during tissue dissociation(*31*, *32*). Consistent with this, 93.9% of immune cell profiles were from scRNAseq rather than snRNAseq data **(Table S2B**).

Subclustering each broad cell type revealed 55 fine-grained renal immune cell states including 22 T/NK (**Fig. 2A**), 21 Myeloid (**Fig. 2B**), and 9 B/Plasma clusters (**Fig. 2C**). Within the renal parenchymal tissue cell types, we obtained 36 states, including 8 proximal tubule (PT, **Fig. 2D**), 5 loop of Henle (LOH, **Fig. 2E**), 8 distal nephron (DN, **Fig. 2F**), 10 endothelial/stromal (EC, **Fig. 2G**), and 5 glomerular clusters (GLOM, **Fig. 2H**). Marker genes and cell state frequencies were concordant across scRNAseq and snRNAseq data **(Fig. S3-10)**. Read distributions across parenchymal cell types were largely consistent between each other and between modalities **(Fig. S2A)**. Immune cell read distributions were consistent with each other but captured less transcripts in snRNAseq than scRNAseq **(Fig. S2A)**

**Fig. 2.**
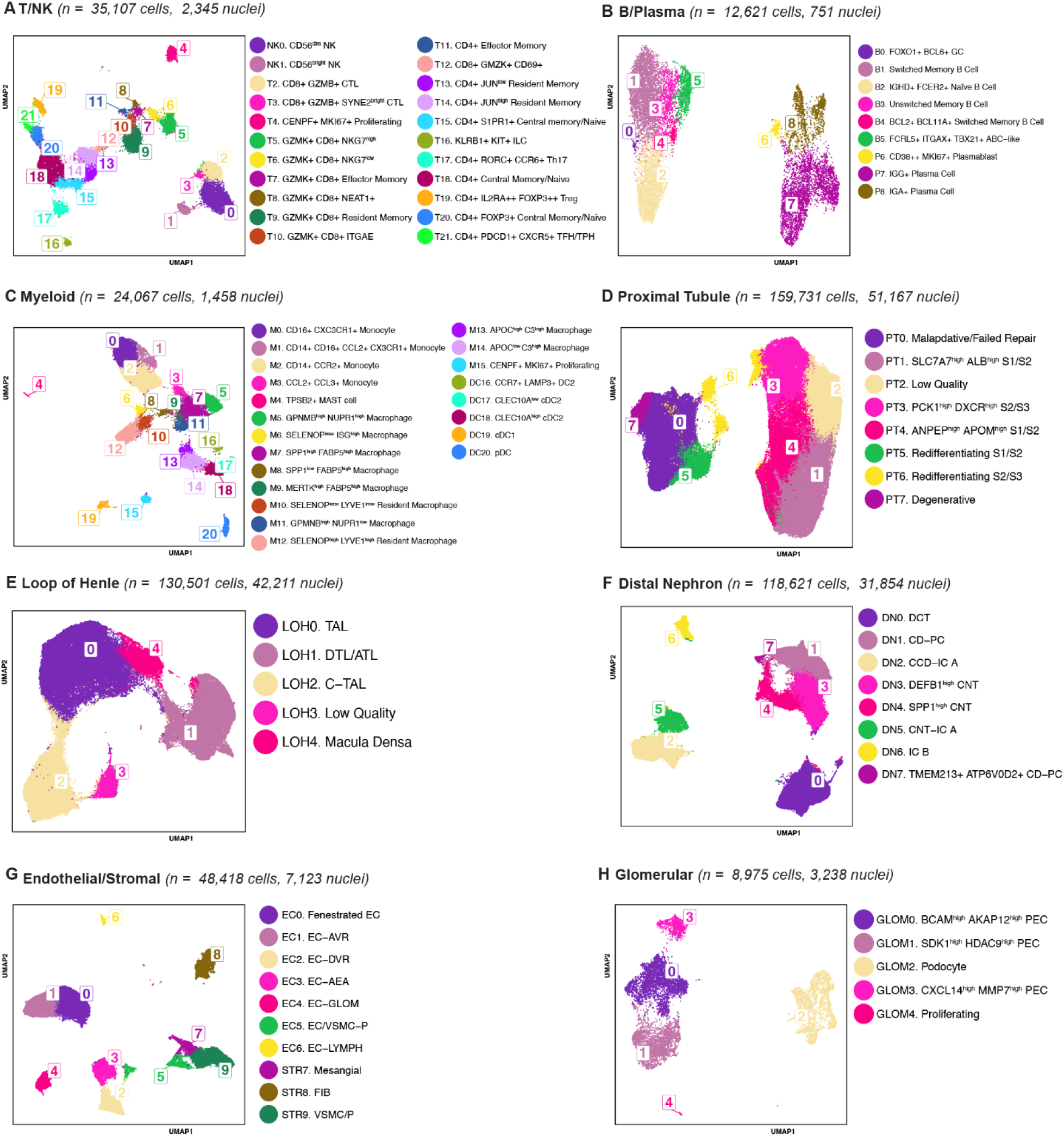
Cell state annotations in tissue. **(A)** UMAP of T/NK cell states in tissue, colored by fine grain cell state annotations. Annotation boxes are located at the centroid (mean) of each cluster. **(B-H),** Same as **(A)** for B/plasma, myeloid, proximal tubule, loop of Henle, distal nephron, endothelial/stromal, and glomerular cells, respectively.

### Renal Immune Cell States

Of the 22 T/NK clusters, two were NK clusters with high (NK0) or low (NK1) *NCAM1* (*CD56*) expression (**Table S4**). *CD8* T cell clusters included central memory (*SELL*+) and effector (*GZMK*+*/GZMB*+) subsets (**Fig. S3**), with nine total *CD8* states. This included two *CD8+GZMB*+ cytotoxic lymphocyte (CTL) clusters (T2, T3) and a *CD69+ITGAE+CCR7-SELL-S1PR1-CD8+* resident memory-like cluster (T9). We classified *CD4* clusters into memory and naive subsets using *CD28*, *CD69*, *SELL*, *CCR7*, *ITGAE*, and *S1PR1*. We identified two CD4 *CD28*+*CD69+SELL*-*ITGAE+CCR7*-*S1PR1-*resident memory clusters (T13, T14), along with a *CD28+CCR7-ITGAE-SELL-* effector memory-like cluster (T11), and specialized subsets, including *IL2RA*+*FOXP3+* Tregs (T19), *PDCD1*+*CXCR5+* T Follicular helper (TFH/TPH) (T21), and *RORC*+*KLRB1*+*CCR6+* Th17 (T17).

We identified 6 B cell states, expressing *MS4A1*, *BACH2*, and *IL4R*, and 3 Plasma cell states, expressing *SDC1*, *XBP1*, and *PRDM1* (**Table S3, Fig. S4)** We identified a *FOXO1*+*BCL6*+ germinal center (GC)-like B cell cluster (B0) and a *FCRL5*+*ITGAX*+*TBX21*+ autoimmune-associated B cell (ABC) cluster (B5)(*33*), with high and low expression of *IGHG2* and *IGHD*, respectively. We identified a *CD38+CD27*+*MKI67*+ plasmablast-like cluster with low expression of *IGHD* (P8). Plasma cells clustered by *IGG* (P6) or *IGA* (P7) heavy chain immunoglobulin genes.

Within the 21 myeloid clusters, we distinguished macrophages, monocytes, and dendritic cells with marker genes including *APOE* and *C1Q* (**Table S3**, **Fig. S5**). Four monocyte clusters (M0, M1, M2, M3) spanned the *CD14*+ (classical) to *CD16*+ (non-classical) gradient. Several macrophage clusters coexpressed *SPP1, LPL, CD9, TREM2, CTSB,* and *CTSD*. Within this group, we identified two *GPNMB*^high^ macrophage clusters: *GPNMB*^high^*NUPR1*^high^ (M5) and *GPNMB*^high^*NUPR1*^low^ (M11). We also identified three *FABP5*^high^ clusters: *SPP1*^high^*FABP5*^high^ (M7), *SPP1*^low^*FABP5*^high^ (M8), *MERTK*^high^*FABP5*^high^ (M9). We distinguished three macrophage clusters by *SELENOP* and *LYVE1* expression levels: *SELENOP*^inter^*ISG*^high^ (M6), *SELENOP*^inter^*LYVE1*^inter^ (M10) and *SELENOP*^high^*LYVE1*^high^ resident macrophages (M12). Dendritic cell subsets included two cDC2 clusters (DC15, DC17), *CCR7*+*LAMP3*+ DCs (DC16), pDCs (DC20), and a cDC1 *XCR1*+*CLEC9A*+ cluster (DC19).

### Renal Parenchymal Cell States

We identified 5 glomerular (GLOM) clusters, including podocytes (GLOM2), expressing *PODXL*, *NPHS2*, and *WT1*, and parietal epithelial cells (PECs), expressing *VCAM1* and *CFH* (**Fig. S6**, **Table S4**). The three PEC clusters included *BCAM*^high^*AKAP12*^high^ PEC (GLOM0), *SDK1*^high^*HDAC9*^high^ PEC (GLOM1) with high *TNC*, and *CXCL14*^high^*MP7*^high^ PEC (GLOM3). A proliferating cluster (GLOM4) expressed *HELLS* and *CPNK*. GLOM1 notably expresses Epithelial-to-Mesenchymal Transition markers over other PEC subtypes (**Fig. S30A, B)**.

We identified 8 proximal tubule (PT) clusters, using *SLC34A1, SLC22A8,* and *MOGAT1* to delineate anatomical segments (S1, S2, S3) (**Fig. S7**, **Table S4**). (PT1-4) reflected anatomical segmentation, while PT0 and PT5-7 reflected injured populations. Anatomical clusters included two S1/S2 (PT1, PT4) and one S2/S3 cluster (PT3). S1/S2 coexpressed S1 (*PRODH2*, *SLC5A2*, *SLC22A8*) and S2 genes (*SLC34A1, SLC22A7*) while S2/S3 coexpressed S2 and S3 (*MOGAT1, SATB2, ABCC3*) genes. Injury clusters PT5 and PT6 indicated successful redifferentiation into S1/S2 and S2/S3 subtypes, respectively, marked by *LRP2*, *MAF* in PT5 and *LRP2*, *PAX2*, and *HNF4A* in PT6(*34*). PT0 expressed *VCAM1*, *TPM1*, and *DCDC2*(*35, 36*) indicating maladaptive redifferentiation following injury(*37*). PT7 expressed *FOS*, *JUN*, and *EGR1,* associated with a degenerative injury response(*38*, *39*).

We identified 5 loop of Henle (LOH) clusters (**Fig. S8, Table S4**) including descending thin limb (DTL), ascending thin limb (ATL) cells, thick ascending limb (TAL), and macula densa cells, defined by *SLC12A1, CRYAB*, *TMEM25B*, and *EGF* expression (**Fig. S8**). The TAL cluster (LOH0) expressed *CYFIP2*, *MFSD4A, RAP1GAP,* and *ANK2*. Cortical TAL (C-TAL) (LOH2) expressed *TMEM52B, CLDN16,* and *WNK1.* A mixed DTL/ATL cluster (LOH1) expressed *CRYAB, TACSTD2,* and *SLC44A5.* Macula Densa (LOH4) expressed *PAPPA2* and *NOS1*. A low-quality cluster (LOH3) showed high mitochondrial and ribosomal expression.

We identified 8 fine-grain distal nephron (DN) clusters (**Fig. S9, Table S4**), grouped into distal convoluted tubule (DCT), intercalated cells (IC), connecting tubule (CNT), and collecting duct principal cells (CD-PC) based on expression of *CLNK*, *SLC12A3*, *AQP2*, and *SLC8A1* (**Fig. S9**). Two CNT clusters coexpressed *SLC8A1*, *CALB1,* and *SC2NA*, and were distinguished by *DEFB1* (DN3) or *SPP1* (DN4).

Two CD-PC clusters (DN1, DN7), expressed *GATA3*, *SCNN1G, SCNN1B* and *FXYD4*. IC subtypes included IC-beta-like CD-PCs (DN7) expressing *INSRR*, *SLC35F3*, and *TLDC2* and IC-alpha cell states resembling CNT cells (DN5), expressing *SC2NA* and *CALB1,* or cortical collecting duct cells (DN2), expressing *ADGRF5* and *LEF1*.

Renal endothelium and stroma (EC/STROMA) yielded 10 cell states: fibroblasts (FIB), endothelial (EC), vascular smooth muscle cells (VSMC), and mesangial cells, distinguished by *PECAM1*, *COL1A1*, *PLVAP*, and *NOTCH3* expression (**Fig. S10**, **Table S4**). Two *PLVAP*^high^ fenestrated EC clusters (EC0, EC1), represented either fenestrated cells of unknown origin or from the ascending vasa recta (EC-AVR). Other EC subsets included descending vasa recta (EC2), ascendant efferent arteriole (AEA) (EC3), glomerular *EMCN*^high^ (EC4), and lymphatic ECs (EC6). VSMCs and pericytes (STR9, EC5) shared markers *MCAM, NTRK3,* and *RGS5.* FIBs (STR8) expressed *DCN, COL1A2,* and *COL1A1*.

Mesangial cells (STR7) expressed *POSTN, GATA3,* and *DAAM3*. Globally, we find that transcript counts per gene between scRNAseq and snRNAseq are consistent **(Fig. S11A-H)**.

### Comparing Population Frequencies in Single Cell and Spatial Transcriptomic Data

We noted the low frequency of some of the immune populations in our data, most notably B cells.

Since emerging spatial transcriptomic technologies do not require disaggregation, we consider cell-type frequencies from such data to likely be accurate. We obtained a spatial dataset of childhood-onset SLE (cSLE) consisting of 7 case biopsies and 4 controls profiled by a 980-gene CosMX panel(*25*). However, spatial transcriptomic data often lacks the high per-cell information content of single cell data: scRNAseq and snRNAseq data capture a median 5,906 transcripts per cell compared to 168 in spatial transcriptomics. **(Fig. S12A, B)**. Spatial transcriptomics however captures more cells per biopsy (median of 1,190 immune cells in spatial data compared to 189 in scRNAseq and snRNAseq). Although we have fewer immune cells collected per biopsy **(Fig. S12C)**, immune cells constitute a greater proportion of scRNAseq than of spatial methods, particularly in case samples (14.5 vs 4.9%, **Fig. S12D)**; in contrast snRNA-seq had a somewhat lower proportion of immune cells (3.6% of case samples, **Fig S12D**). Within the immune compartment, T and NK cells are relatively enriched in scRNAseq data compared to spatial data, whereas B and myeloid proportions remain similar **(Fig. S12E).** In B cells, the sparsest of our immune cell types, the proportions between spatial and scRNAseq data are consistent with the proportions in spatial data **(Fig. S12C, D, Fig. S13A)**. This concordance suggests that the low abundance of B cells (2.2% of total cells and nuclei in LN samples) reflects limited biological infiltration rather than a technical limitation of single-cell profiling approaches. Glomerular cells made up a lower proportion of total cells collected compared to the spatial dataset (12.6% vs 1.7%, **Fig. S13G**), which may reflect the possibility of lower glomerular cell yield in single cell data than other proportions. Other parenchymal cell types were concordant between our data and spatial data **(Fig. S13A, B)** except for loop of Henle cells, which were overrepresented in scRNAseq and snRNAseq relative to spatial data, while endothelial/stromal cells were underrepresented.

### Defining Shared and Distinct Immune Populations in Blood and Tissue

We profiled 324,995 PBMCs from 118 LN patients and 19 healthy controls and annotated cell states with canonical marker genes (**Fig. S14**). We obtained 114,376 T cells, 52,324 NK Cells, 99,878 monocytes, 5,450 dendritic cells, and 52,967 B Cells (**Fig. S14A**). To identify immune cell states shared between kidney and blood, or distinct to either tissue type, we integrated both kidney and PBMC immune populations into a joint nearest neighbor embedding (**Fig. 3**). In this joint embedding, we preserved the original cell annotations from the respective separate tissue and blood cluster analyses. We use this embedding to illustrate the correspondence between tissue and blood clusters (**Fig. S15D-F**). Using this joint embedding, we developed a per-cell tissue specificity metric (TSM), defined as the proportion of a given cell’s k=50 nearest neighbors derived from renal tissue, weighted for differences in dataset size (**Fig. 3A**). A TSM of 1 indicated a tissue-specific cell state, 0 indicated a blood-specific state, and 0.5 reflected equal representation in tissue and blood. TSM remains consistent across all cell types for varying choices of k (**Fig. S15A-C**). We find that per-cluster median TSM rankings remain stable following downsampling the highly abundant blood single cell data **(Fig. S16A-C)**

**Fig. 3.**
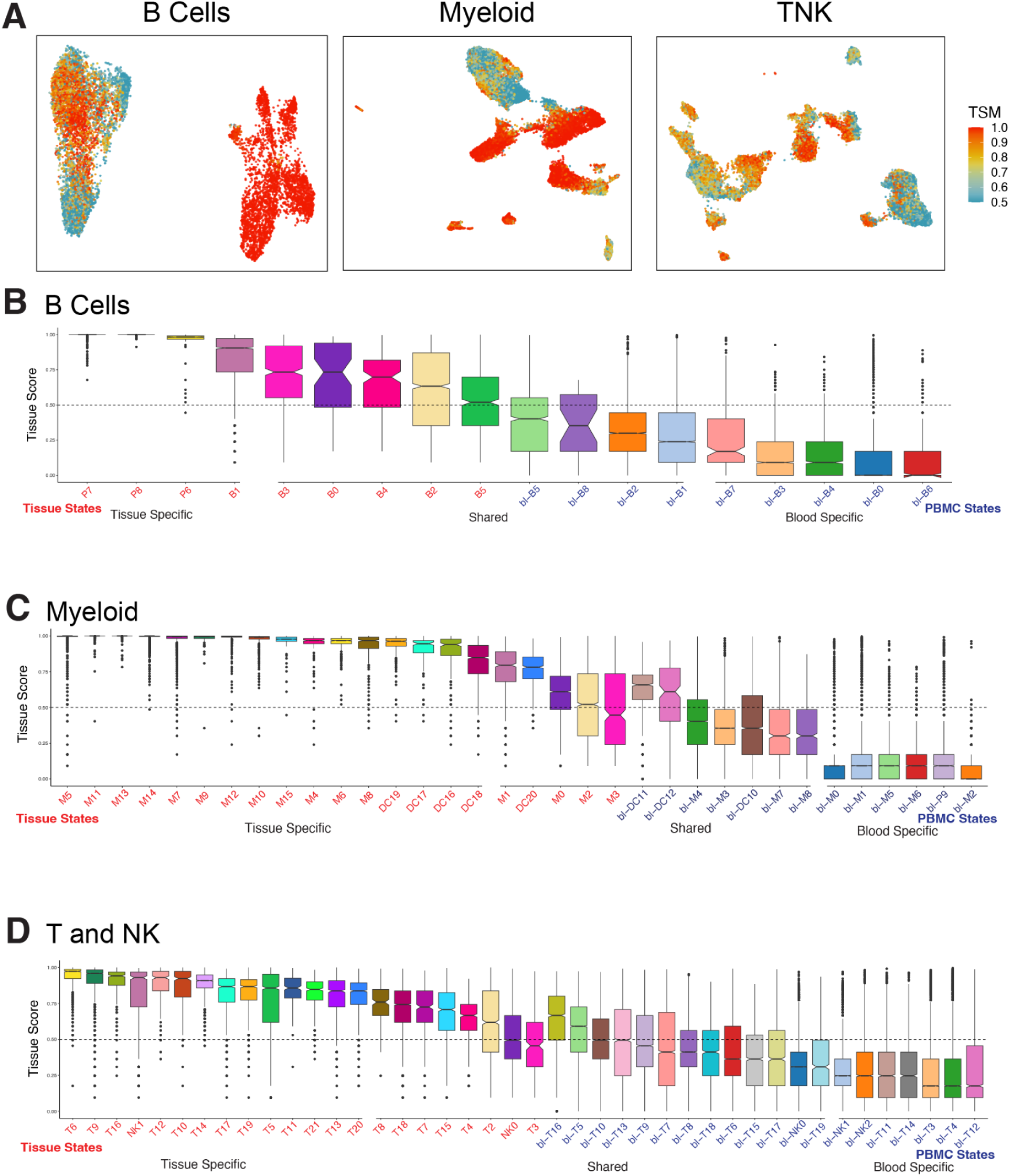
Tissue Specificity Metric. **(A)** UMAPs of TSM score for immune cells from kidney tissue for B cells (left), myeloid cells (center) and T and NK cells (right). (**B-D)** Tissue specificity metric of tissue and blood immune cells stratified by cluster for B cells **(B),** Myeloid cells **(C)**, and T and NK cells **(D)**.

Plasma cell and plasmablast clusters (P6-P8) were tissue-specific, with TSM scores near 1 (**Fig. 3A, B**). The ABC−like cluster from tissue (B5) co-embedded with blood-derived ABCs (bl-B2) (**Fig. S15D**), both with a median TSM ∼0.5, consistent with previous findings that ABCs spanned both blood and tissue(*33*, *40*). We wanted to understand B cell differentiation in our data. Hence, using our matched PBMC and tissue modalities, we re-embedded B cells in a joint Destiny(*41*) space and applied Slingshot(*42*) to infer trajectory pseudotimes to reflect B cell lineage. We specified blood-specific naive B cells (bl-B0. CXCR5^high^ Naive) as the starting point in our trajectories. Within this trajectory, pseudotime progresses through memory states toward B5. FCRL5+ ITGAX+ TBX21+ ABC-like and bl-B2 FCRL5⁺ ITGAX⁺ ABCs **(Fig. S17A).** Genes correlated with pseudotime include ITGAX (**Fig. S17B**), indicating that the trajectory may capture continuous differentiation of the ABC cell state.

In myeloid cells, macrophage populations (M5-12, M14, M16) were tissue-specific (**Fig. 3C**). Conversely, monocyte populations (M0-2) were shared, likely reflecting migration from the blood. Classical DCs (DC16-19) were tissue-specific, while plasmacytoid DCs (DC20) were shared with blood (**Fig. 3C**), consistent with prior observations(*43*, *44*). Non-classical monocytes (bl-M3, bl-M11) were shared between tissue and blood, while classical monocytes (bl-M0-2, bl-M5) were blood-specific. To understand myeloid cell infiltration and differentiation we applied trajectory analysis. Pseudotime inference on Destiny re-embeddings identified two trajectories. The first trajectory proceeds from blood monocytes through tissue macrophages to *GPNMB*^high^ macrophage clusters M5, M7, and M9 **(Fig. S17C)**.

This trajectory may indicate that *GPNMB*^high^ macrophage are derived from blood monocytes, with a tissue monocyte and other macrophage cell states as intermediaries. Future fate mapping experiments may be more informative as to the direct lineage of these populations. The second trajectory involves only blood and inflammatory tissue monocytes. Both monocyte-to-*GPNMB*^high^ macrophage and within-monocyte trajectories correlate with our TSM metric (R = 0.506 and R = 0.511, respectively).

T/NK cell TSM scores had fewer strictly blood- or tissue-specific populations. Tissue-skewed clusters included *CD8*+*GZMK*+ populations (T5, T6, T12) (**Fig. 3D**), consistent with previous work(*45*). TFH/TPH populations (T21), implicated in autoimmune rheumatic disease(*46*), and resident memory populations (T9, T10, T13-14) were tissue-skewed. Both CTL clusters (T2, 3) were shared. Some central and effector memory T cells were tissue-skewed (T11, T20), but most (T7, T15, T18) were shared, likely reflecting tissue infiltration from blood. We observed a shared blood regulatory T cell population (bl-T16), and a tissue-skewed Treg cluster (T19), consistent with our previous analyses(*47*). In trajectory analysis, we observed that blood and tissue CD4+ **(Fig. S17D)** and CD8+ **(Fig. S17E)** T cells have a shared naive memory trajectory. Genes correlated with pseudotime are enriched for interferon alpha and gamma response pathways.

In B/plasma cells, differential gene expression between donor-matched tissue and PBMC samples revealed an upregulation of plasma cell markers (*IGH3*, *IGLC2*, *IGHCHG1*) in tissue (**Fig. S18A, B**). In the B/plasma clusters identified in tissue, the tissue-specific (high TSM) plasma clusters were enriched for these differentially expressed tissue markers. Conversely, the shared cell states (TSM around 0.5) had comparatively higher expression of genes differentially expressed by PBMCs.

Differential gene expression in Myeloid cells revealed similar patterns. In tissue, genes associated with macrophages (*C1QA*, *C1QB*, *C1QC*) and anti-inflammatory properties (*GPNMB*) were upregulated (**Fig. S18C, D**). Markers high in blood monocytes (*SELL*, *LYZ*) and pro-inflammatory properties (*S100A8*, *S100A9*) were downregulated. Within the tissue compartment, tissue-specific Myeloid clusters, including all macrophage states, were relatively enriched for the differentially expressed tissue markers. Shared monocyte states were relatively enriched for differentially expressed blood markers.

Differential gene expression in T/NK cells revealed increased T cell activation in tissue, indicated by upregulation of activation markers (*CD40LG*, *PDCD1*) and cytokine and chemokine secretion (*IFNG*, *TNF*, *CCL3*) (**Fig. S18E, F**). T cells from tissue were also significantly enriched for markers of tissue-residence, including *ITGA1*, *CXCR6*, and *ZNF683* (nominally). While tissue markers were expressed across most renal T cell states, differentially expressed blood markers were relatively enriched in shared T cell states, including *CD56*^dim^ NK (NK0) and *CD8*+*GZMB*+*SYNE2*^bright^ CTL (T3) cells.

### Kidney Case Control Analysis

To understand key differences between LN and unaffected kidney, we performed case-control comparisons with CNA. When we examined all kidney cell types together, we observed expansion of immune populations, reflecting immune infiltration (p<1×10^-4^ for both scRNAseq and snRNAseq) (**Fig. S19A, B**). Parenchymal compartment differences between LN patients and living donors may reflect tissue damage or differences in biopsy tissue procurement (see **Methods)**.

Case-control analysis in scRNAseq data within each cell type revealed fine-grained cell state shifts (**Fig. 4**). We detected shifts in all cell types (p<1e-04), except in B/Plasma cells, where low control cell counts were unable to generate an informative CNA result. We did not adjust for age and sex in our analysis as doing so had minimal impact (**Fig. S20**). Within T/NK cells, the greatest shifts were in tissue-specific populations (**Fig. 4A**) including enrichment of *CD4+IL2RA++FOXP3+* Tregs (T19), *CD4+FOXP3+* Central Memory/Naïve (T20), TFH/TPHs (T21), and *CD56*^bright^ NKs (NK1) and contraction of resident memory populations *GZMK+CD8+* Resident Memory (T9), *CD8+GZMK+CD69+* cells (T12), and *CD4+JUN*^high^ Resident Memory cells (T14). We also observed tissue-specific myeloid population shifts, including enrichment of five macrophage populations: *SELENOP*^inter^*LYVE1*^inter^ Resident Macrophage (M10), *GPMNB*^high^*NUPR1*^low^ Macrophage (M11), *GPNMB*^high^*NUPR1*^high^ Macrophages (M5), *SELENOP*^inter^*ISG*^high^ Macrophage (M6), and *SPP1*^high^*FABP5*^high^ Macrophages (M7) and contraction of two: *APOC*^high^*C3*^high^ Macrophages (M13) and *APOC*^low^*C3*^high^ Macrophages (M14) (**Fig. 4B**).

**Fig. 4.**
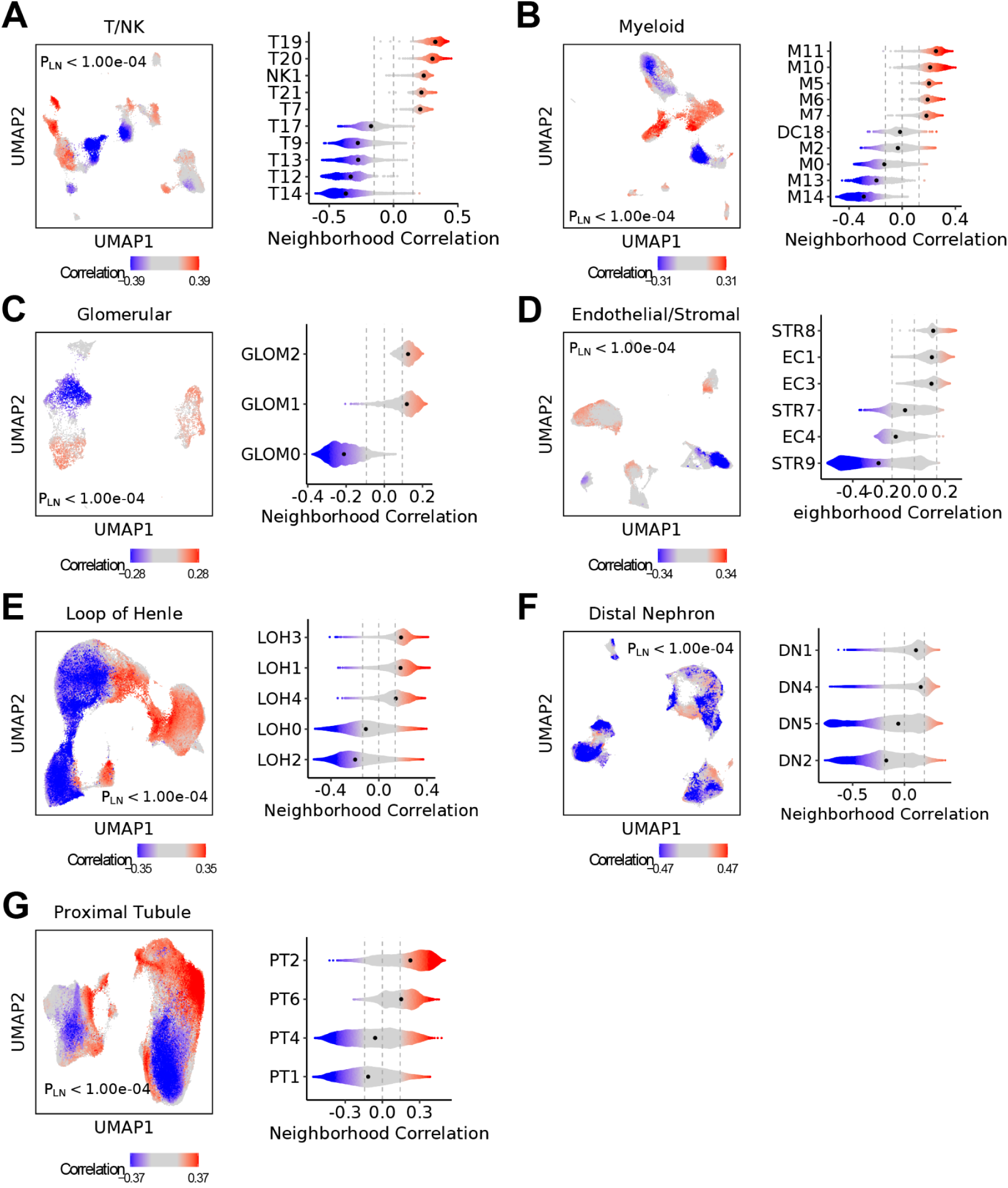
Case-control associations within cell types. **A-G** CNA results for case-control association with no covariate corrections, in scRNA-seq data. Left - UMAP displaying significant per-cell associations with LN, with FDR cutoff of 0.05. Blue and red denote contracted and expanded neighborhoods that pass FDR, respectively. Grey represents neighborhoods that do not pass FDR. Non-significant associations are colored in grey. P-value is the global P-value for cell state associations with LN phenotype. Right - violin plots of clusters containing cells passing FDR significance for LN association. The median is denoted with a black dot. Analysis was repeated for (**A),** T/NK cells, (**B),** myeloid cells, (**C),** B/plasma cells, (**D),** endothelial/stromal cells, (**E),** loop of Henle cells, (**F),** distal nephron, **(G),** proximal tubule cells.

We identified shifts in the renal parenchymal populations. Activated/injury-associated PECs (GLOM1, *SDK1*^high^*HDAC9*^high^) were unsurprisingly expanded in LN, while homeostatic PECs (GLOM0, BCAM^high^AKAP12^high^) contracted (**Fig. 4C**)(*48*, *49*). Podocytes (GLOM2) were relatively expanded in LN, potentially due to depletion of healthy PECs. In Endothelial/Stromal cells, fibroblasts (STR8), EC-AVR (EC1), and EC-AEA (EC3) expanded, while VSMC/pericytes (STR9) contracted (**Fig. 4D**). In the loop of Henle, neighborhoods within TAL (LOH0) and C-TAL (LOH2) demonstrate heterogeneity, with finer subpopulations having positive and negative correlation values, indicating statistically significant expansion and contraction in subpopulations of cells. The median negative correlation indicates that, considered in aggregate, these populations are contracting. Other cell types such as DTL/ATL (LOH1) and Macula Densa (LOH4) expanded in LN samples (**Fig. 4E**). In the Distal Nephron, *SPP1*^high^ CNT (DN4) and CD-PC (DN1) expanded (**Fig. 4F**). Among Proximal Tubule cells, *PKHD1*^high^*SLC3A1*^high^ S2/S3 (PT6) was expanded, while *SLC7A7*^high^*ALB*^high^ S1/S2 (PT1) was contracted (**Fig. 4G**). Contraction of specific states may reflect the loss of vulnerable cells in disease; for example, TAL cells have high metabolic demand and are sensitive to hypoxia(*50*). To confirm that this is not driven by case-control cell number imbalances (**Table S2A**), we subsetted the number of case samples to match controls and further downsampled cells within cases to match cell-type distributions (**Fig. S21A**). Repeating CNA in this reduced data set finds that the associations remain concordant and globally significant (**Fig. S21B, C**).

Differential gene expression in LN and control samples revealed widespread upregulation of genes across cell types (**Fig. S19C**). Unsurprisingly, interferon-stimulated genes (ISGs) were enriched (*IFI44L*, *IF16*, *MX1*) across cell types, as were genes related to immune activation (*STAT1*, *SPP1*)(*51*, *52*). Several genes were broadly downregulated in LN, including innate and innate-like lymphoid cell regulators (*53*) (*ZBTB16*, *NFKBIA*, *PDK4*, *KLF9*, *FKBP5*). Individual cell types revealed enrichment for additional disease-relevant programs, including extracellular matrix (ECM) remodelling (*COL1A1*, *MMP7* in parenchymal cell types), fibrosis and wound healing (*TIMP1* in parenchymal cell types, *CDH11* in Endothelial/Stromal, *HBEGF* in Glomerular), lysosomal activity (*CTSD*, *LGMN*, *PCCA* in Myeloid), class II HLA expression (*HLA-DMB*, *HLA-DRB1* in parenchymal cell types and T/NK), and immune activation and regulation (*TIGIT*, *IL27RA* in T/NK, *LCN2* in loop of Henle) (**Fig. S19C**). Gene set enrichment analysis (GSEA) confirmed upregulation of the well-described interferon response pathway (**Fig. S19D**)(*23*, *54*).

### Defining the Effect of Sample Level Covariates on LN Tissue Cell States

After noting case-control differences, we wanted to understand how cell states changed across LN samples. To understand this, we applied CNA to test clinical variables and pathological variables, including seven key variables capturing LN tissue state. (5 ISN classes, activity, chronicity). Although SLE manifestations differ between sexes (*55*), we did not find any globally significant associations with sex **(Fig. 5A)**. This may be due to the imbalance in our predominantly-female cohort, as is typical of SLE **(Table S1).** We observed that chronicity consistently had the most dramatic effects; changes were significant across all cell types, except B cells, where we were likely underpowered to detect any effects (**Fig. 5A**). We speculated that many alterations due to increased chronicity reflect worsening tissue damage to the kidney.

**Fig. 5.**
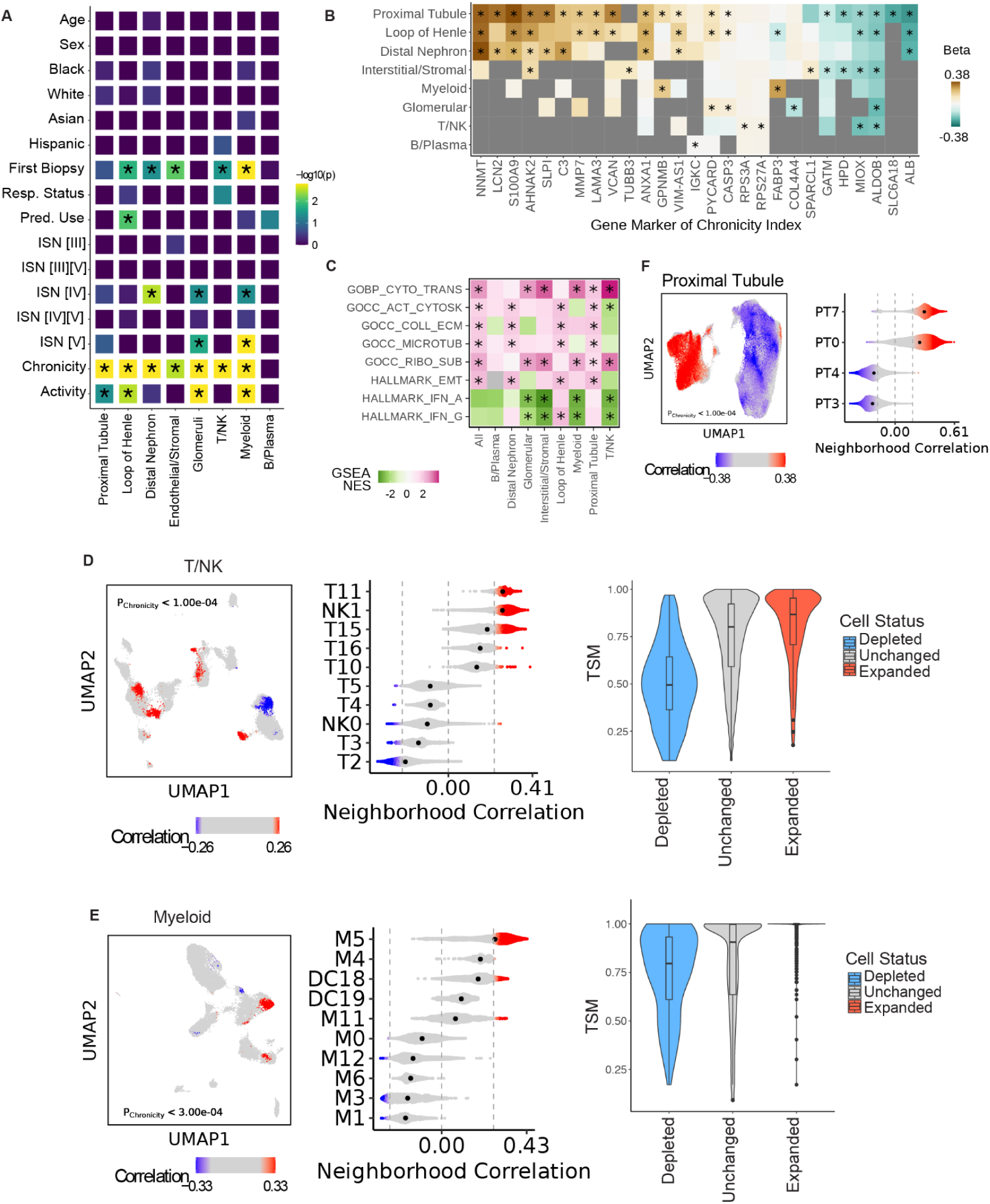
Cell states associated with chronicity index. **(A)**, Univariate CNA of each cell type testing for relevant clinical and demographic variables. * indicates a global p-value < 0.05. (**B),** For each cell type, differential expression for genes associated with chronicity index. * indicates a Bonferroni-adjusted p-value less than 0.05. Grey color indicates the gene was not tested due to low expression in that cell type. **(C),** Normalized enrichment scores (NES) from pathway enrichment of differential gene expression results, using Hallmark and Ontology pathways. * indicates a Bonferroni-adjusted p-value less than 0.05. Grey color indicates the pathway was not tested due to low expression of pathway genes in that cell type. **(D),** Across myeloid cell types, CNA results for chronicity index association while adjusting for first biopsy and site, in scRNA-seq data. Left - UMAP displaying significant per-cell associations with chronicity index, with FDR cutoff of 0.1. Non-significant associations are colored in grey. P-value is the global P-value for cell state associations with chronicity index phenotype. Middle - violin plots of the top 5 and bottom 5 clusters by median cell state correlation containing cells passing FDR significance for chronicity index association. Dashed vertical lines represent the correlation threshold with FDR < 0.1. Right - Within T/NK cells, violin plots displaying per-cell TSM scores for cells found to be expanded, depleted, or neither with chronicity index, as identified using CNA. **(E),** Same as **(D)** for Myeloid cells. **(F),** Same as **(D)** for PT cells, without TSM score derivations.

### Defining Cell State Correlations with Chronicity

The chronicity index reflects irreversible damage to the kidney and is associated with reduced treatment response(*16*, *56*). Differential expression analysis revealed gene programs associated with chronicity index across cell types (**Fig. 5B, C**). This included genes related to ECM remodeling or fibrosis (*VCAN*, *MMP7*, *TUBB3*), inflammation and immune response (*S100A9*, *GPNMB*, *C3*), and apoptosis (*CASP3*). The metabolic enzyme *NNMT*, previously linked to renal injury(*57*, *58*), was highly upregulated in multiple renal cell types. While the most significant changes occurred in parenchymal cell types, immune populations also showed upregulation of immune response genes (**Table S5**). Chronicity index was also correlated with total immune cell proportion in a given sample (sc: R=0.31, P=1.2×10^-4^; sn: R=0.41, P=0.015), suggesting chronic inflammation tracked with increased permanent tissue damage (**Fig. S22C**). GSEA revealed enrichment for ECM remodeling, epithelial-mesenchymal transition, and translation pathways (**Fig. 5C**). ECM remodeling was enriched in endothelial/stromal (nominally) and proximal tubule cells, consistent with prior reports of profibrotic genes in these cell types(*59*), as well as in distal nephron and loop of Henle cells. Interferon response pathways were downregulated in T/NK, B, Myeloid, endothelial/stromal, and glomerular cells (**Fig. 5C**) but weakly upregulated in loop of Henle and proximal tubule cells, suggesting cell type-specific effects.

To understand the relationship with interferon response, we constructed an ISG score from a curated list of interferon-stimulated genes(*23*). LN cases had higher ISG scores than controls in both tissue and immune populations (**Fig. S23A**). Among LN cases, the ISG score declined with increasing chronicity but remained stable across activity index levels (**Fig. S23B**).

We noted that chronicity was correlated with collection variables, including non-first biopsy status and biopsy site (**Fig. S2D, E**). Since chronicity increases with ongoing disease, the chronicity index was lower in the first biopsy (p=1.14e-06) (**Fig. S23F**). Patient recruitment site was correlated with many clinical variables, including first biopsy status (**Fig. S23D**); we suspected site-specific differences reflect both demographic and technical factors. To account for this, we adjusted for both first biopsy status and major recruitment sites as potential confounders in all chronicity analyses. Consistent with prior reports(*16*), chronicity was lower in complete responders than in non-responders (p=0.002) and partial responders (p=0.04) (**Fig. S23F**).

CNA analysis of broad cell types revealed associations with increasing chronicity (FDR<0.05, p<10.00e-04); these associations included expansion of T/NK, myeloid, B/plasma, and glomerular cells, with heterogeneous shifts in proximal tubule and loop of Henle cells (**Fig. S23C**). CNA within each cell type revealed fine cell state shifts **(Table S6)**. In T/NK Cells (FDR<0.05, p=0.006), tissue-specific *CD56*^bright^ NK (NK1) and some *CD4*+ Memory/naive clusters (T11, T15) were strongly expanded (**Fig. 5D**). This expansion was further corroborated at the sample level in NK1 (1.4% at chronicity=0, with +0.46% per unit chronicity; P = 1E-4), T11 (0.32% at chronicity=0, + 0.13% per unit chronicity; P = 8E-4), and T15 (2.2% at chronicity=0, +0.46% per unit chronicity; P = 6.7E-5) **(Table S7)**. In our data set 2.44 points of chronicity correspond to 1 standard deviation. Tissue-specific cell states *GZMK*+*CD8*+*ITGAE*+ (T10) and ILCs (T16) were weakly expanded. Conversely, most contracted cell states were shared with blood, including CTL (T2, T3), *CD56*^dim^ NK (NK0), and proliferating T cells (T4). In Myeloid cells (FDR<0.05, p=2e-4), increasing chronicity tracked with a contraction of monocyte populations shared in the blood and tissue (M0, M1, M3) as well as two tissue resident macrophages (M6, M12) (**Fig. 5E**). Conversely, tissue-specific macrophages (M5, M14) and cDCs (DC18) were expanded with increasing chronicity. These populations were also expanded at the sample-level **(Table S7)**. In both immune populations, cell populations contracted with increased chronicity had a lower TSM score (less tissue-specific) than those that were not.

Increased chronicity was associated with contraction of glomerular endothelial cells (EC4) (FDR<0.05, p=1.1e-3) and expansion of fibroblasts (STR8) in the stromal compartment (**Fig. S24A**). In glomeruli, podocytes were contracted (GLOM2) (FDR<0.05, p<9.99e-05) while injury-associated PEC subsets (GLOM1, GLOM3) were expanded (**Fig. S24B**). In the Proximal Tubule (FDR<0.05, p<9.99e-05), the late-injury PT0 state was highly expanded, while healthy PT1/PT2 states were contracted (**Fig. 5F**). Cell states in the loop of Henle and distal nephron showed both expansions and contractions with chronicity (**Fig. S24C, D**, FDR<0.05, p<10.0e-04 both). Adjusting for treatment response had minimal impact (**Table S8, Fig. S24E**).

Although we were underpowered to detect B cell associations with CNA, we attempted to utilize clonotypes to try and uncover clones expanded in higher chronicity samples. However, clonotyping was limited by low expression of immunoglobulin transcripts **(Fig. S25A)**, a known issue of 3’ sequencing’s ability to determine clonotypic information(*60*). Light chain gene expression was especially sparse in B cells, and of the cells that did have non-zero light chain gene expression, no light chain genes were expanded with higher chronicity **(Fig. S25B).**

While B cells are more clinically relevant, plasma cells had more light chain gene expression **(Fig. S25A)**. However, we did not observe any clones across multiple samples that expanded with chronicity **(Fig. S25C)**. IGKV3-11 and IGKV3D-11 gene usage was expanded in a small subset of cells, but all of these cells originate from a single high chronicity patient sample **(Fig. S25D).**

### Proximal Tubule Injury Drives Chronicity-Associated Cell State Shifts

In CNA analysis, the proximal tubule was the most significantly associated population with chronicity, and had the largest number of significant (FDR<5%) local neighborhood shifts **(Table S9)**. Of the proximal tubule cell states, a key epithelial injury state, PT0 maladaptive/failed repair (R=0.542, P=1.01E-10) was most correlated with at the sample-level with increasing chronicity (**Table S7**) but only weakly associated with treatment response and not associated with ISN class (**Fig. S26A, B**). PT0 had a frequency of 14% at chronicity=0, and expanded 3.93% per unit chronicity (p<4.6E-12).

CNA within each cell type, adjusting for recruitment site and first biopsy status, revealed that increasing PT0 proportion was significantly associated with cell state shifts in the loop of Henle (p<10e-04), distal nephron (p<10.00e-04), endothelial/stromal (p<10.00e-04), glomerular (p<10.00e-04), T/NK (p=2.6e-03), and myeloid compartments (p=2.00e-04) (**Fig. S26C**). The PT0-associated shifts were concordant with those associated with increasing chronicity (**Fig. S26D-I**), and adjusting for PT0 proportion reduced the significance of cell state associations with chronicity. However, PT0-associated cell state shifts remained significant even after correcting for chronicity (**Fig. S26C**). For example, Glomerular shifts strongly associated with chronicity (p<2e-05) became less significant after adjusting for PT0 (p<1e-2), while Glomerular shifts associated with PT0 remained significant even after adjusting for chronicity (p<1e-05 vs p<7e-05). Accordingly, we suspect that proximal tubule injury, marked by the size of the PT0 population, is a proxy for the cellular effects of higher chronicity index and may better reflect tissue damage than chronicity index in LN itself. Within PT0, we find no differentially expressed genes that distinguish between response groups **(Fig. S27A-C).**

### Activity is Associated with Expansion of Myeloid and Glomerular Subpopulations

The activity index is a dynamic pathological measure that reflects inflammation and reversible damage in the kidney(*61*), and cell states associated with this index may represent therapeutic targets. To identify cell state shifts associated with activity, we adjusted for first biopsy status, recruitment site, and chronicity due to the broad effects of these variables on cell states (**Fig. S23D**). Across cell types, we observed significant shifts (p<4.5e-04), driven by shifts in the myeloid (p<1e-5) and glomerular (p<2e-5) populations (**Fig. 6A**). These associations were not confounded by age or sex **(Fig. S28).** Differential gene expression further revealed the largest number of genes associated with activity index in these same compartments (**Table S10**).

**Fig. 6.**
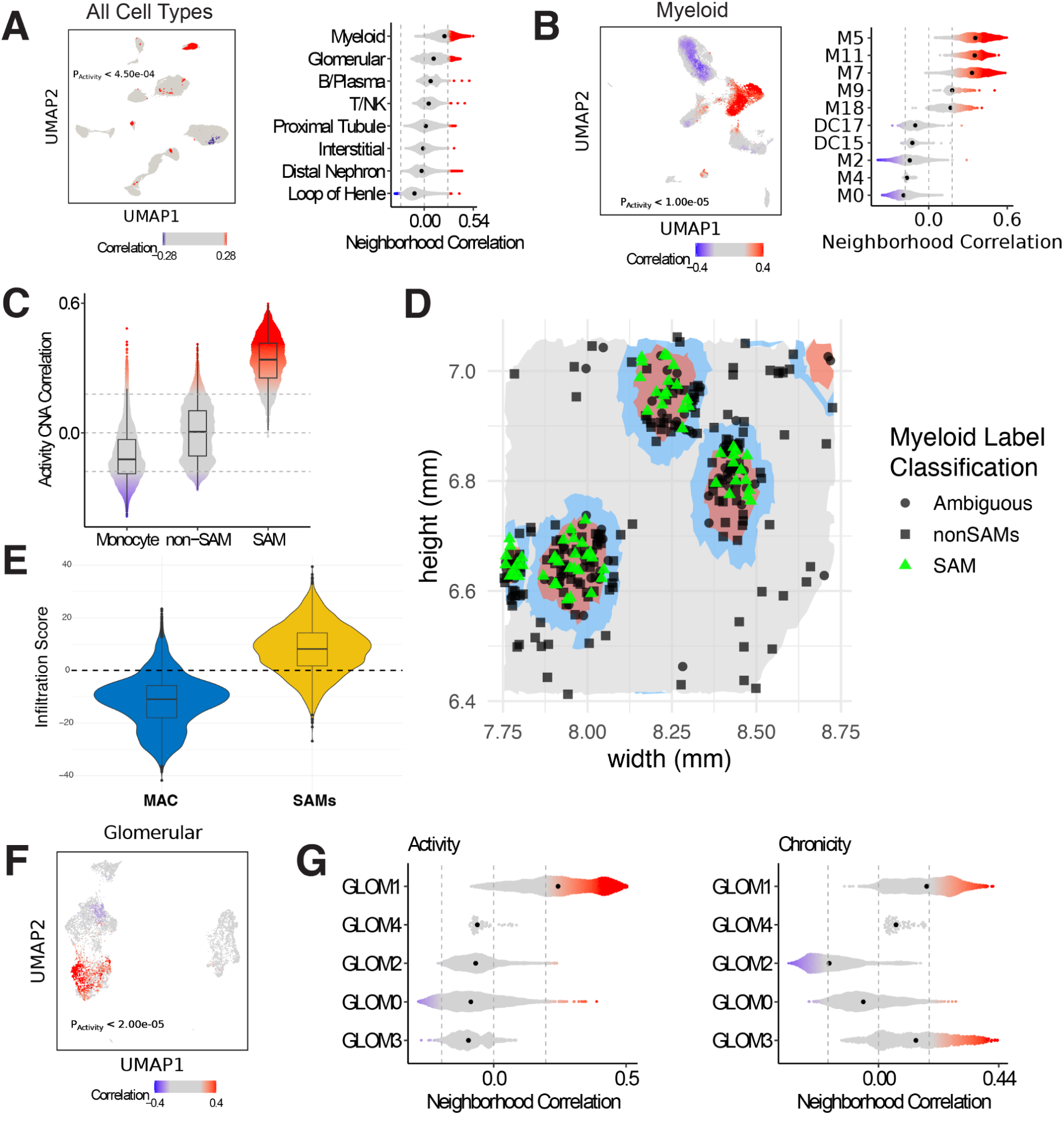
Cell states associated with activity index. **(A)**, Across all cell types, CNA results for activity index association while adjusting for first biopsy, site, and chronicity index, in scRNA-seq data. Left - UMAP displaying significant per-cell associations with activity index, with FDR cutoff of 0.1. Dashed vertical lines represent the correlation threshold with FDR < 0.1. Non-significant associations are colored in grey. P-value is the global P-value for cell state associations with activity index phenotype. Right - violin plots of clusters containing cells passing FDR significance for activity index association. **(B),** Same as **(A)** for myeloid cells. Left - Same as **(A)** for myeloid cells. Right - Within myeloid cells, violin plots of clusters containing cells passing FDR significance for activity index association (adjusting for first biopsy, site, and chronicity). **(C),** Violin plot of activity CAN correlation grouped by broad myeloid category. **(D),** Select case FOV spatial plot of non-myeloid cells colored by spatial niche. Gray indicates tubulointerstitium, blue indicates the periglomerular region and red indicates the glomerular region. Myeloid cells distinguished by shape and color. **(E)** Violin plot of infiltration score applied on scRNAseq and snRNAseq myeloid cells. **(F)** Same as **(A)** Left for glomerular cells. **(G),** Violin plots for CNA correlations in glomerular cell types for activity (left) and chronicity (right).

Although activity index correlates with ISN class and prednisone use (**Fig. S29A**) and has been associated with treatment response in previous studies(*62*), adjusting for these variables did not obviate associations (**Fig. S29B-C**), suggesting that activity independently drives these cell state shifts.

CNA on myeloid cells showed expansion of macrophages (**Fig. 6B**), notably *GPNMB*^high^*NUPR1*^high^ (M5), *GPNMB*^high^*LYVE1^low^* (M11), *C1Q*^low^ *SPP1*^high^, *SPP1*^high^*FABP5*^high^ (M7), and *MERTK*^high^*FABP5*^high^ (M9) (**Fig. 6B**). Populations shared with blood, including *TPSB2*+ MAST cells (M4) and *CD16*+*CXC3CR1*+ Monocytes (M0), contracted with increasing activity. While M5 was also expanded with increasing chronicity, all other activity-expanded populations did not expand with increasing chronicity. For instance, *CCL2*+*CCL3*+ Monocytes (M3) and *CLEC10A*^high^ cDC (DC18) were depleted with increasing chronicity, but expanded with increasing activity. This suggests that activity-associated shifts are primarily driven by tissue-specific macrophages with effects independent of chronicity.

We observed that marker genes for Scar Associated Macrophages (SAMs) defined populations expanding with increasing activity. *CD9* and *GPNMB* were highly correlated (R>0.7) with CNA activity associations in myeloid cells, differentially expressed with activity index, and highly expressed in M5 (**Fig. S29H**). Cells invading the glomeruli in patients with focal segmental glomerulosclerosis (*FSGS*) and crescentic glomerulonephritis express *CD9* and the prevalence of such cells reflect different degrees of podocyte injury(*49*). *GPNMB* (R=0.713) is a known biomarker of SLE previously found in the urine and exosomes in a patient with acute LN(*63*). Both genes are known markers for SAMs(*64*, *65*) which were detected in patients with liver or lung fibrosis. Other SAM genes, including *FABP5* (R=0.68), *TREM2* (R=0.63), and *SPP1* (R=0.55) were also highly correlated and were highly expressed in 4 out of 10 macrophage clusters (M5, M7, M9, M11) (**Fig. S29D**). While a small subset of SAM macrophages expand with increasing chronicity (**Fig. S29E)**, almost all SAMs are strongly expanding with activity (**Fig. 6C**). SAM proportions of the myeloid compartment at the sample level similarly showed expansion (6.4% at 0 activity; +2.00% per unit activity; P = 5.2E-14) **(Fig. S30A),** which is consistent with the CNA results. Three out of four SAM clusters show significant sample-level expansion, specifically M5 (1.2% at 0 activity; +0.77% per unit activity; P = 1.3E-8), M7 (3.4% at 0 activity; +0.87% per unit activity; P = 1.2E-9), and M11 (1.3% at 0 activity; +0.26% per unit activity; P = 4.4E-7) **(New Table S11).** As with activity, SAM proportions of the myeloid compartment at the time of biopsy do not appear to be indicative of treatment response in our cohort **(Fig. S30B)**. To test whether the expansion of M5 or other SAMs was driven by tissue-resident populations, we developed a proliferation score from 93 cell cycle genes but found no evidence of proliferation (**Fig. S29F**).

Additionally, differential gene expression showed activity index was associated with lipid metabolism (*APOC1* - enriched in M5/M11, *FABP5*, *PLIN2* - enriched in M5) and phagocytic/lysosomal activity genes (*TREM2* - enriched in M5/M7/M9/M11, *CTSD* - enriched in M5, *CTSB* - enriched in M5/M7/M9). Cytoskeleton and motility-related genes (*TUBB6* - enriched in M7, *VIM*) were also enriched with respect to activity index. Differential gene expression between SAM and other myeloid populations reveals upregulation of pro-inflammatory markers such as the receptor Integrin Beta-1 (ITGB1) (**Fig. S30C**), which belongs to a family of integrin receptors known to bind to and activate latent TGFB(*66*). Consistent with this, integrin receptors and other upregulated genes are enriched for the EMT pathway, which is the most significantly enriched in SAMs (**Fig. S30D**). EMT has been shown in previous work as a necessary prerequisite in murine organ fibrosis(*67*).

To better understand where SAMs localize, we examined a spatial data set of 11 samples (see section **Comparing Population Frequencies in Single Cell and Spatial Transcriptomic Data**)(*25*). To identify SAMs in this dataset, we trained a logistic regression classifier to distinguish SAM versus non-SAM myeloid cells using 258 genes that are present in both our data set and the spatial transcriptomic data set, and are highly expressed in the Danaher et al. dataset. **(Methods).** Testing the classifier on an unseen subset of across 10 folds myeloid cells in our data demonstrated an average AUC of 0.933 **(Fig. S31A).** Applying this classifier to spatial data, we find that macrophages and SAMs localize to glomerular niches **(Fig. 6F, Fig. S31B, C)**. Across all spatial samples, SAMs comprise a higher percentage of total cells in SLE cases over control samples **(Fig. S31D)**. Within case samples, SAMs are present in the glomeruli at a higher rate than non-SAM myeloid cells **(Fig. S31E).**

Conversely, SAMs obtained by scRNAseq and snRNAseq resemble macrophages previously identified in spatial data to be localized to the glomeruli. From spatial transcriptomic data, we defined a Glomerular Infiltration Score (GIS) consisting of the 196 differentially expressed genes previously reported upregulated or downregulated in the glomeruli-localized macrophages over interstitial macrophages (FDR < 0.05)(*25*). Applying this score on our dataset we find that SAMs have a higher infiltration score over other non-SAMs **(Fig. 6G)**. Samples with higher median infiltration score have higher activity index **(Fig. S31F)**. This suggests that our SAMs are not only associated with glomerular damage as measured by the activity index but are also infiltrating the glomeruli at a higher proportion than other non-activity associated macrophage populations.

Among glomerular cells, cell states associated with increasing activity index were similar to those associated with chronicity (**Fig. S29G,** R=0.487), including strong expansion of *SDK1*^high^*HDAC9*^high^ PEC (GLOM1) (**Fig. 6B, C**). This expansion was also observed at the sample level (15.0% at 0 activity; +1.90% per unit activity; P = 6.6E-7) **(New Table S11).** GLOM1 displayed a transcriptional program consistent with activated, injury-associated PECs, characterized by the expression of genes involved in extracellular matrix production (*FN1*, *COL8A1*, *COL6A2*, *LOXL1*), epithelial-to-mesenchymal transition (*SOX4*, *CDH6*), cytoskeletal remodeling (*ACTN1*, *TPM1*), and matrix degradation/remodeling (*MMP7*, *TNC*); reflecting a pro-fibrotic phenotype observed in glomerular injury and crescentic glomerulonephritis(*10*, *64*, *68*, *69*). Notably, GLOM1 selectively expresses *TNC* (Tenascin C), an extracellular matrix glycoprotein that is transiently induced during tissue injury and repair(*70*). We have previously shown that urinary Tenascin C levels correlate with histological activity—and, to a lesser extent, chronicity—in patients with LN(*71*), mirroring the associations observed here.

Differential expression in glomerular cells revealed enrichment of genes associated with ECM remodeling and fibrosis (*FBN1*, *PLOD2*) and inflammation and interferon signaling (*SPP1*, *STAT2*) (**Fig. S29H**). *MFGE8*, previously implicated in SLE disease activity and risk(*72*, *73*), was also enriched. All are markers of the activity-associated cluster GLOM1. Differential expression analysis within PECs **(Fig. S32A)** suggests that GLOM1 is enriched for the epithelial to mesenchymal transition pathway **(Fig. S32B)**. To test whether activity-associated expansions are due to proliferation, we developed a proliferation score from 93 cell cycle genes and found that GLOM1 had a higher proliferation score than other PEC populations (**Fig. S32C)**.

Given the localization of SAMs to the glomeruli, we wondered if they interacted with glomerular cells. We employed receptor-ligand analysis between glomerular cells and tissue immune cells. We observed that the glomerular-myeloid pairing has the most interactions among the cell types **(Fig. S32F).** Statistically significant glomerular and myeloid receptor-ligand pairings implicate SPP1 and TNC. SPP1, a known SAM marker interacts with cell adhesion molecule CD44 and 4 integrin receptor heterodimers **(Fig. S32E)**. Two of these heterodimers show strong interaction with SPP1 in GLOM1 **(Fig. S32F, G)**.

ITGB1, which forms a heterodimer with ITGAV, is upregulated in GLOM1 compared with other PECs **(Fig. S32A)** and is also part of the EMT pathway. Another integrin heterodimer, ITGB3 + ITGAV, is strongly enriched for SPP1 and GLOM1 interactions; ITGB3 is not significantly expressed in GLOM1 over other PECs (p = 0.99). Although not previously characterized in SLE, secreted SPP1 promotes cell migration as part of the EMT pathway (*74*, *75*).

GLOM1 upregulated Tenascin C (TNC) which has two main binding partners: SDC4 and the ITGAV+ITGB3 heterodimer. TNC expressed by GLOM1 is found to interact with the SDC4 receptor found on SAMs and proliferating myeloid cells **(Fig. S32I).** Furthermore, GLOM1-expressed TNC interacts with the aforementioned pan-glomerular ITGAV+ITGB3 receptor **(Fig. S32J)**. As previously mentioned, our group has found TNC as a potential urine biomarker correlated with activity in SLE. While the integrin-TNC link has not been explicitly studied in SLE, it is well-known to trigger the EMT pathway in cancer(*76*, *77*); the EMT pathway has been shown to contribute to renal fibrosis(*78*).

Given that the chronicity index is divided into indices representing glomerular or tubulointerstitial damage and 21 out of 24 points in the activity index are based on measures of glomerular inflammation, we examined whether applying CNA to the individual subcomponents could reflect different aspects of LN pathology.

### Activity and Chronicity Index Subcomponents

Activity and chronicity indices are comprised of subindices representing glomerular or tubulointerstitial damage. For example, 21 out of 24 points in the activity index are based on measures of glomerular inflammation. We examined whether applying CNA to the individual subcomponents could reflect different aspects of LN pathology.

In chronicity, however, apart from fibrous crescents, most indices correlate with each other, including measures of glomerular or tubulointerstitial damage **(Fig. S33A).** CNA on fibrous crescents reveals new neighborhoods in distal nephron and myeloid cells **(Fig. S33B, C)** that were not previously identified as associated with chronicity **(Table S11).** While DN4 primarily expands with higher chronicity, higher fibrous crescents are marked exclusively by expanded distal convolute tubule (DN0) populations **(Fig. 33D, E).** In the myeloid compartment, fibrous crescent CNA has a lower correlation with chronicity CNA (R = 0.442) than it does with activity CNA (R = 0.715) **(Fig. S33F)**. While M5 remains the most significant with fibrous crescent as it does with chronicity, dendritic cells are no longer significantly expanded **(Fig. S33G)**. Other macrophage populations M11 and M7 remain significantly expanded. Previous pathological studies suggest that fibrous crescents may indicate early lesions that eventually lead to glomerulosclerosis (*79*). Furthermore, our group has previously reported that the urine proteomic signature in high fibrous crescent index is more correlated with activity rather than chronicity (*71*). So we felt that quantification of fibrous crescent may capture a mixture of chronicity and activity.

In activity, three components correlated with overall activity at the sample level index with R < 0.6, fibrinoid necrosis, wire loops, and interstitial inflammation **(Fig. S34A)**. Notably, interstitial inflammation is the lone component that does not reflect glomerular damage. However, these associations were only significant in myeloid cells **(Table S12)**. Within myeloid cells, neighborhood-level associations for these indices were highly correlated with activity **(Fig. S34B)** and did not identify many novel neighborhoods not previously identified as passing FDR significance in activity CNA **(Fig. S34C-E).**

### Activity Associations are Independent of ISN Classification

We were interested to see if these activity changes could be better explained as a difference between membranous (ISN class V) vs proliferative nephritis, since membranous nephritis is clinically defined as having low activity.

Class V was significantly associated with Myeloid (p<9.99e-05) and glomerular (p=0.0029) populations (**Fig. S36A, B**), even after adjusting for first biopsy status, chronicity, and collection site. In both compartments, populations expanded in ISN Class V were contracted with higher activity. Expanded populations included *TPSB2*+ MAST cells (M4) and *CD16*+*CXC3CR1*+ Monocytes (M0), while contracted populations included *SPP1*^low^*FABP5*^high^ (M8), *GPNMB*^high^*LYVE1*^low^ (M11), *SELENOP*^inter^*LYVE1*^inter^ Resident (M10), *SPP1*^high^*FABP5*^high^ (M7), and *MERTK*^high^*FABP5*^high^ (M9) macrophages (**Fig. S35A**). In the Glomeruli, *SDK1*^high^*HDAC9*^high^ PEC (GLOM1) were also contracted (**Fig. S35B**). Results were unchanged after adjustment for prednisone use and Black race, both of which correlated with ISN Class V (**Fig. S23D**, **Fig. S35C**).

However, after adjusting for activity index, glomerular associations were no longer globally significant, and myeloid associations were much less significant. While activity-associated CNA results remained consistent after correcting for ISN V, ISN V associations did not (**Fig. 35D, E**). These findings suggest that cell state shifts observed in Class V can be explained by its reduced activity index.

### Clinical Phenotype Associations with Immune Cells from Blood

Given the easier access to blood than kidney tissue, we examined blood cell state changes associated with case-control status, chronicity, and activity indices. After correcting for age and sex, we identified cell state shifts in B (p<7e-5), monocytes and DCs (p<3e-4), and T/NK cells (p<5e-5) (**Fig. S36A-C**). LN cases had expansion of ISG-high monocytes (bl-M6), ISG-high naive B cells (bl-B4), and ISG-high T cells (bl-T17), corroborating previous reports of interferon-inducible gene upregulation in LN blood (*80*). Dendritic cells (bl-DC10, bl-DC11, bl-DC12) were contracted, consistent with previous studies of SLE blood(*81*, *82*). ABCs (bl-B2) were expanded, aligning with prior human patient and murine lupus models(*40*, *83*). CXCR5^high^ Naive (bl-B0) and CRIP1+ Mature B cells (bl-B1) were contracted. The T/NK compartment showed reduced *CD4*+*IL7R*^high^*VIM*^high^ Central Memory (bl-T5), CD4+ Naive (bl-T10), and *CD8*+*GZMK*+ TEMRA T cells (bl-T7), with strong depletion of *TRGC1*+ Gamma/Delta T cells (bl-T12). Differential expression further showed enrichment of interferon-stimulated genes and depletion of ribosomal genes in LN (**Fig. S36D**).

Given the strong association between renal pathology indices and tissue cell states, we examined whether there were corresponding associations with blood cell states. After adjusting for age and sex, we observed only modest associations with chronicity in Myeloid (p=0.013) and T/NK cells (p=0.0019) (**Table S13, 14**). In Myeloid cells, few neighborhoods (n=46/106,436) were significant (FDR<0.10). In T/NK cells, *CD8*+*GZMK*+ TEMRA T cells (bl-T7) were expanded with increasing chronicity but were depleted in cases over controls (**Figure S36E**). Notably, after correcting for age, sex, there were no significant associations with activity. The limited activity index associations in blood does not rule out the possibility of immunophenotype associations that may be present with different patient subgroups; for example, in a recent study on the same cohort, statistical differences were seen in B cell states across patient subgroups (*84*).

Similarly, differential expression analyses revealed only weak gene associations with chronicity index (**Fig. S36F,G**), including downregulation of interferon-stimulated genes in T/NK and B/Plasma cells. No strong gene associations with activity index were observed. Together, these results suggest that, while blood signals may help identify SLE cases and modestly reflect chronicity, blood cell state populations are unlikely to reliably capture renal disease activity.

## Discussion

Clinicians have long recognized the heterogeneity of clinical manifestations in LN and the clinical value of renal biopsy. Here, we present a large cross-sectional study comparing kidney biopsies from LN patients and healthy controls, with matched PBMCs for many samples. We included both single-cell and single-nucleus data to robustly define tissue cell states independent of technical effects. This publicly available dataset is a valuable resource for future integration efforts into lupus and renal biology. Our goal was to characterize key cell states in renal tissue and blood, and how these states shift across pathological subsets. We also assessed whether those tissue state shifts manifested in the circulation.

We identified a spectrum of immune, epithelial, and stromal populations relevant to LN, including autoimmune-associated B cells in both blood and tissue (bl-B2, B5), consistent with previous findings in the blood of SLE patients(*85*), as well as Proximal Tubule epithelial cells with a degenerative injury profile (PT0). We also observed tissue-specific immune states such as *GZMK*+*CD8*+ T cells, *GPNMB*^high^ macrophages (M5, M11), and *SPP1*^high^*FABP5*^high^ macrophages (M7). To characterize how these populations changed with clinical subphenotypes in LN, we adjusted for potential confounders such as recruitment sites and whether the biopsy was the first one obtained from a patient, since subsequent biopsies often reflect greater damage due to a greater disease burden. Our results include key observations related to LN that will inform future studies and guide therapeutic decisions.

The high interferon signature in SLE has driven drug development for both LN and non-LN manifestations. While its link to SLE in blood is well established(*23*), associations with clinical disease activity and tissue inflammation are less well understood. In our data, interferon signatures were clearly enriched in LN compared to controls, across blood and tissue. However, connections between interferon signature and tissue inflammation states were more subtle, with interferon signature showing only a modest decline with increasing chronicity in some cell types, and no association with activity.

We observed subtle shifts, such as expansion of *GZMK*+*CD8*+ T cells in blood with increasing chronicity, that paralleled changes in tissue. However, overall, blood expression and cell states had limited associations_with respect to chronicity or activity. This does not preclude the possibility that alternative profiling technologies may reveal cellular or serum biomarkers reflecting LN renal heterogeneity(*48*). In contrast, non-invasive biomarkers, including macrophage-derived proteins in urine, have been associated with proliferative disease and decrease in treatment response(*71*).

We were particularly interested in the activity index, which reflects ongoing renal inflammation and is thought to be modifiable with immunomodulatory therapies. Even brief reductions in activity index has been associated with reduced organ damage(*86*). After adjusting for the substantive effects of chronicity, we observed that key, tissue-specific cell states within the myeloid compartment increased with activity, including *GPNMB*^high^ (M5, M11)(*63*) and *SPP1*^high^*FABP5*^high^ (M7) macrophages. These populations expressed a canonical *SPP1*, *CD9*, *GPNMB*, *FABP5*, and *TREM2* macrophage gene profile, known to occur across multiple chronic inflammatory diseases, often adjacent to fibrotic regions(*87–89*). We leverage a classifier built on SAM transcriptional profiles to find that SAM macrophages identified in a spatial dataset infiltrate the glomeruli at a higher proportion than non-SAMs. In addition, receptor-ligand analysis indicates that SAMs interact specifically with activity-associated glomerular populations through the EMT pathway.

SAM subsets are thought to be heterogeneous, with some populations being pro-fibrotic and others being reparative. (*90–92*). The persistence of the M5 subset in association with both disease activity and chronicity may suggest a particularly pathogenic role. These populations appear to be similar to SAMs reported in other diseases and tissue contexts(*93*). We note that SAMs may not be specific to LN and may play a role in other renal diseases. Further comparative studies between other renal diseases may address this in the future.

This data does not definitively establish whether these states drive disease activity, mediate tissue repair, or promote fibrosis but rather highlights distinct, disease-associated states with therapeutic potential. Furthermore, while our trajectory analysis suggest that SAMs may infiltrate from blood into the tissue, past work in the brain and liver have both suggested that SAMs may have different origins based on their subset, with some being derived from infiltrating monocytes and others differentiating from tissue-resident macrophage populations (*94*, *95*). Mouse models may be useful in defining causal functions of this and other myeloid populations (*96*). Current SLE therapies primarily target B and T cells (*5*, *97*). While B cell targeting has been effective, our study was underpowered to detect activity associations with B cell states. We recognize that single cell analysis requires tissue disaggregation and droplet-based assays, which limits the amount of cells profiled. In addition, there may be biases in terms of cell type proportions. We find that, compared to spatial datasets, our single cell data has a lower proportion of glomerular and endothelial/stromal cells. We find that immune populations are not significantly different between spatial and single-cell modalities, and our lack of B cells may reflect limited infiltration rather than technical collection limitations. The low frequency of B cells in LN is consistent with the idea that B cells are acting elsewhere in other tissues to produce damaging immune complexes. The data presented here supports the potential for targeting myeloid cells in LN in conjunction with B cells.

While we are, to our knowledge, the largest single-cell LN tissue dataset, spatial methods collect many more cells per biopsy and may result in even larger atlases. Single cell data, however, still offers much greater depth of information for each individual cell with over >30-fold transcripts per cell enabling fine cell state definitions. The next phase of this project, AMP AIM, is currently working towards building a large data set of spatial transcriptomics for LN to complement the information in this dataset(*98*).

Renal biopsy chronicity index scores are thought to reflect cumulative, irreversible kidney damage from ongoing disease. Consistent with this, we observed that chronicity closely tracked with a shift from healthy to injured proximal tubule cells. These injured epithelial cells show cellular dedifferentiation, senescence, and a canonical transcriptional signature linked to tubulointerstitial fibrosis and chronic kidney disease (CKD) progression across diverse renal diseases (*99*). These differences may not be specific to LN. Our results suggest that these changes may very closely reflect the level of chronic kidney injury, and may serve as a molecular proxy for chronicity.

Within renal tissue, chronicity had the most dramatic impact on cell types and states, including immune populations, compared to other clinical parameters. Notably, with increasing chronicity, *SPP1*^high^*FABP5*^high^ macrophages (M7) and *GZMK*+ T cells were greatly expanded within the myeloid and T cell compartments, respectively. Most immune cell states that expand with increasing chronicity are tissue-specific and largely absent in the blood, including *GPNMB*^high^*NUPR1*^high^ Macrophage (M5) and a *CD4*+ Effector Memory T cell state (T11). These findings demonstrate the importance of accounting for chronicity in molecular studies of LN. For instance, associations between immune and tissue cell states and treatment response disappear after adjusting for chronicity, a well-known predictor of response(*16*). A key finding is that differences in renal tissue cell states are better explained by activity and chronicity indices than by ISN classes, which are commonly used for informing therapies. For instance, ISN class V patients by definition have a decreased activity index (which is focused on lesions specific to class III and IV), it is the low activity index index and not the ISN class that explains the cell state shifts.

This study has several limitations. Intriguingly, B cells were still relatively rare in the tissue for our study, despite the large number of cells and samples assayed. This was surprising considering the importance of B cells as an SLE drug target. Given the similar proportions of B cells in our atlas and an external spatial dataset, it is likely that this is not a technology issue. B cells are pathogenically active in other tissues, such as lymph nodes, and are less active in LN kidneys. It is possible that there are important rare cell states that track with disease activity in the kidney, but our study would lack the power to identify this given the few cells assayed.

However, other pathogenically important cell types, such as glomerular cells, are still relatively depleted over spatial data where tissue architecture is not as disturbed. This is a known limitation of single-cell data, but glomerular cells have been rescued in other studies through a Percoll gradient(*100*, *101*). Additionally, renal immune cell states were defined transcriptionally, without surface protein markers, which can improve immune cell state annotation accuracy(*102*). Our study design was also highly heterogeneous, including a large number of samples from a range of recruitment sites but also individuals at different disease and biopsy points, and with different treatment histories. While the diversity of the cohort is a strength, its heterogeneity complicates disentangling medications and cohort variables on biopsy metrics. Longitudinal followup samples were also not performed. Finally, while our tissue-specificity metric (TSM) identifies tissue and blood resident populations, it cannot distinguish whether tissue specificity is due to migration from blood or development within the tissue.

In conclusion, this comprehensive study defines the key tissue and blood cell states in SLE, and demonstrates how they shift with the development of LN, and within the renal tissue of LN cases. This useful reference will serve as a key resource in understanding the pathogenesis of LN, and devising therapies to mitigate its long-term effects on patients.

## Methods and Materials

### Study Design

We have described the details of enrollment in the AMP SLE cohort elsewhere(*16*, *103*, *104*). In brief, patients with lupus undergoing kidney biopsies as part of standard of care were eligible to enroll in the prospective AMP LN study. The decision to biopsy was at the discretion of the treating physician to confirm suspected LN *de novo*, assess persistent disease activity not responding to treatment, or diagnose disease recurrence. Inclusion in AMP required: 1) age ≥ 18; 2) fulfilment of the revised American College of Rheumatology(*105*) or the Systemic Lupus Erythematosus International Cooperating Clinics(*106*) classification criteria for SLE; 3) a urine protein/creatinine ratio (UPCR) > 0.5 at the time of biopsy. This study began prior to the publication of the ACR/EULAR criteria. Analyses regarding responder status were restricted to patients with baseline random or 24-hour Urine Protein Creatine Ratio (UPCR) ≥ 1.0 since for patients with UPCRs between 0.5 and 0.999, we did not define proteinuria response. We only considered patients with renal biopsies that demonstrated International Society of Nephrology/Renal Pathology Society (ISN/RPS) classes III, IV, V or combined III or IV with V read in this analysis(*13*, *107*). Where possible, ISN Class and histopathological indices were sourced from the central pathologist scoring. For patients without a central read, we used the read from the pathologist at the recruitment site. Exclusion criteria included: 1) a history of kidney transplant; 2) rituximab treatment within 6 months of biopsy; 3) pregnancy at the time of biopsy. Informed consent was obtained from all participants, and the study protocol was approved by the institutional review boards and ethics committees of participating sites in adherence with the Declaration of Helsinki. One sample was later discovered to have chronic lymphocytic leukemia and was excluded from downstream analyses.

Baseline demographics from a predetermined set of categories, including self-reported race (Asian, Black, White, Other)/ethnicity (Hispanic, non-Hispanic) as required for NIH-funded studies, and clinical characteristics were recorded at the time of biopsy. Laboratory tests and medications were documented at each visit (baseline, week 12, week 26 and week 52) and were performed at the participating sites. For steroids, the higher dose at either baseline or week 12 was considered the induction dose for similar reasons. Pulse steroids were also captured separately.

### Defining Fine grain Renal Cell Tissue and Immune Cell States

Within each cell type identified (see above), we employed a fine grain clustering approach consistent with the methods used in broad cell type identification. Briefly, we normalized raw counts, scaled genes to have a mean of zero and variance of one across cells, and employed a weighted PCA approach within each cell type to balance principal component contribution from each modality. We applied an additional level of quality control within each cell type and removed cells of exceptionally low quality (high mitochondrial count and low total read and unique gene counts). We additionally again removed doublets expressing known marker genes from other cell types with a high scrublet score. We identified fine grain cell states using several clustering at high resolution, and identified cell state specific genes using the presto package(*108*). We determined Immune cell subtypes using marker gene lists from the AMP Phase I LN study, the AMP Phase I and II Rheumatoid Arthritis studies, the KPMP project, and a prior in-depth study of memory T cells in tuberculosis patients(*109*). We determined fine grain renal cell types using marker genes sourced primarily from the KPMP project, and supplemented cell state markers using the AMP Phase II Rheumatoid arthritis study as well as sources describing the proximal tubule injury molecular phenotype(*35*, *110*).

### Defining PBMC Fine Grain Cell States

We processed data within each PBMC each obtained cell type. We performed an additional round of doublet removal within each cell type via a combination of high-resolution Louvain clustering, analysis of Scrublet score distribution across clusters, and identification of cluster-specific genes from other cell types. We applied log-normalization to read counts, filtered genes consistently variable in both scRNAseq and snRNAseq data, and then scaled the data. We then applied Harmony^24^ to principal components weighted by scRNAseq to snRNAseq ratio to correct for processing batch, site, sample, and differences in scRNAseq and snRNAseq technologies. We then regenerated harmonized principal components and employed Louvain clustering at several different resolutions. We defined fine grain cell states in T, B, and myeloid cells using markers sourced from the previously described tuberculosis memory t cell study, the AMP Phase 1 SLE study, the AMP Phase 1 and 2 RA studies, Villani et. al(*111*), and Perez et. al(*21*) **(Fig. S14B).**. These markers were matched with clusters via identification of cluster specific gene expression sourced from the presto package(*108*).

Clustering within T and NK Cells obtained 20 clusters spanning CD56^bright^ and CD56^dim^ NK Cells (bl-NK0, bl-NK1, bl-NK2), GZMK+CD8+ (bl-T3, bl-T4, bl-T14, bl-T15), and gamma/delta T cells (bl-T11, bl-T12). TCAT multilabel annotation(*112*) leverages RNA expression data to distinguish between CD4+ central memory (bl-T5) and naive (bl-T10) populations **(Fig. S37A)**. Key marker genes designated between effector memory populations (bl-T9, bl-T13), *GZMK*+*TEMRA* (bl-T7), *CD8*+*GZMB*+proliferating CTLs (bl-T18, bl-T19), Treg (bl-T16), and an interferon-stimulated gene (ISG) high cell state (bl-T17) **(Fig. S37C, D).** CITE-Seq obtained for the PBMC cells further validates these annotations, marking bl-T10 CD4+ naive populations as CD45RA+, CD62L+, and CD45RO- and bl-T5. CD4+ central memory populations as predominantly CD45RO+, CD62L+, and CD45RA-**(Fig. S37B, C)**.

13 monocyte and dendritic cell clusters were delineated as *CD14*+*CD16*-conventional monocytes (bl-M0, bl-M1, bl-M2, bl-M5), *CD14*+ *CD16*+ intermediate monocytes (bl-M4, bl-M8), and *CD14*^dim^ *CD16*++ non-conventional monocytes (bl-M3) (**Fig. S14E, F**). We also found a cluster of ISG^high^ monocytes (bl-M6), CD1C+ DC2 dendritic cells (bl-DC10), *CLEC9A*+ *XCR1*+ DC1 dendritic cells (bl-DC12), *TCF4*+ *CLEC4C*+ pDC dendritic cells (bl-DC11), and platelet cells (bl-P9). B Cells were clustered into 9 specialized substates, including *IGHM*+ or *IGHD*+ naïve B Cells (bl-B0, bl-B4) and *FCRL5*+ *ITGAX*+ auto-immune associated B Cells (ABCs) (bl-B2, bl-B8). (**Fig. S14G, H**). We also found a population of mature B cells expressing low *IGHM* and *IGHD* but high *CRIP1* (bl-B1), a population of *CD79*+*VPREB3*+ pre–B Cells (bl-B3), and *JUN*+*NKFB1*+activated B cells (bl-B6).

### Case-Control Statistical Associations

To understand cell states enriched and depleted in cases, we utilized Covarying Neighborhood Analysis (CNA) (*30*). In brief, CNA divides a given embedding space into granular transcriptional neighborhoods, quantifies the abundance of these neighborhoods per sample, and assesses if the abundance of neighborhoods is associated with sample-level phenotypes. We utilized harmonized principal components to generate CNA neighborhoods. CNA produces a global significance p-value indicating that some neighborhoods are non-randomly associated with the variable being tested. If that value was significant, we visualized CNA outputs by highlighting cells that are at the center of significantly associated neighborhoods (FDR<5%). However, in certain instances where we observed global significance, but did not identify neighborhoods with FDR<5%, we displayed FDR<10% neighborhoods. We applied CNA separately to single-cell and single-nuclear embeddings. When reporting expanded and contracted cell states found by CNA of scRNAseq, we confirmed observed trends with snRNAseq CNA and reported cell states shifts found via both methods. For cell types with less than 10,000 single-nuclei, we confirmed expanded and contracted scRNAseq cell states with MASC and again reported cell state shifts found via both methods. For all PBMC analyses we adjusted for age and sex as covariates.

### Within Case Statistical Associations

We wanted to identify the key clinical variables driving single cell phenotypes within our case samples to better understand heterogeneity within SLE tissues. We selected a mixture of demographic and clinical variables: age, activity index, chronicity index, first biopsy status, ISN Class, prednisone use, race, collection site, treatment response, and sex. Across multiple cell types we found associations across multiple cell types with the key clinical variables of activity index, chronicity index, and ISN Class.

However, we were concerned about confounding variables in the within case analysis. For chronicity index, we computed R-squared coefficients with: age, activity index, biopsy status, ISN Classes, prednisone use, race, treatment response, sex, and site using the lm() functionality in R. For activity index, we computed the R-squared coefficients similarly. For both these analyses we transformed multilevel categorical variables with one-hot encoding. We reported adjusted R-squared coefficients via the lm() functionality in R by fitting the continuous outcome to all one-hot encoded levels but one (as a reference level). For ISN Class we transformed class IV and V into one-hot encoded outcome variables. We employed logistic regression and reported the McFadden’s pseudo-R-squared. For multilevel categorical variables correlated with Class IV and V ISN status, we conducted multiple logistic regression and reported adjusted McFadden’s pseudo R-squared.

To adjust for multiple testing burden, we adjusted correlation p-values with Bonferroni adjustment. However, to ensure robustness of an observed association, we adjusted for variables nominally (p<0.05) associated with the variable of interest. Analysis of differential cell-state abundance and differential gene expression was conducted similarly to steps outlined for case-control analysis above. Differential cell-state abundance testing via CNA and MASC, however, was adjusted for covariates identified through the procedure outlined above.

### Pseudobulk Differential Gene Expression Analysis

To identify differential genes associated with case-control status, we constructed pseudobulk profiles from single-cell RNA profiles within each cell type(*113*). Differential expression analyses were performed separately for tissue and blood single-cell profiles. Pseudo-bulking was conducted via summing the raw gene counts per individual. To reduce false identification of differentially expressed due to sparsity, only genes with non-zero expression in 90% of samples were included. We used two negative binomial models to model the expression of each gene. In the null model for tissue, we modeled gene expression with an intercept and offset for sample library size. In the full model, we performed an identical model with the addition of case-control status as a covariate. All p-values described were calculated using a likelihood ratio test (LRT) between the full and null model. We further corrected for multiple hypothesis testing by adjusting p-values with the Bonferroni method. Identical models were applied to the blood data, with the addition of covariates for age and sex added to both models.

To test for genes differentially expressed with chronicity index, we constructed similar pseudobulk profiles within only case samples, performed gene filtering as in our case-control analysis, and utilized two negative binomial models to model the expression of each gene. Within tissue, we applied a null model for gene expression with an intercept and offset for sample library size and covariates correcting for first biopsy status and site. In the full model, we applied an identical model with the addition of chronicity index as a covariate. Within blood samples, we applied similar null and full models as in tissue, with additional covariates for age and sex added to both models. P-values were calculated with a LRT and adjusted with Bonferroni correction.

To test for genes differentially expressed with activity index, we also constructed pseudobulk profiles within only case samples, performed gene filtering as in our case-control analysis, and utilized two negative binomial models to model the expression of each gene. Within tissue, we applied a null model for gene expression with an intercept and offset for sample library size and covariates correcting for first biopsy status, site, and chronicity index. In the full model, we applied an identical model with the addition of activity index as a covariate. Within blood samples, we applied similar null and full models as in tissue, with additional covariates for age and sex added to both models. P-values were calculated with a LRT and adjusted with Bonferroni correction.

To test for genes differentially expressed between tissue and blood, we subset to only individuals with matched tissue and blood samples. We then constructed pseudobulk profiles from single-cell transcriptomes for each sample. As the sequencing depth between the tissue and blood is very different, we performed stricter quality control prior to modeling and utilized a linear model rather than negative binomial. In addition to including only genes with non-zero expression in 90% of samples, we also filtered out any samples with zero expression of any of these common genes. We then performed library size normalization (TPM) and log10+1 transformation of sample counts. We used two linear models for modeling the expression of each gene. In the null model, we modeled gene expression with an intercept and covariate for individual. In the full model, we performed an identical model with the addition of tissue source as a covariate. P-values were calculated with a LRT and adjusted with Bonferroni correction.

### Statistical Analysis

Correlations between clinical variables were calculated using Pearson correlation for all 155 cases. For correlations with collection sites, we performed multivariate regression correlating clinical variables with all sites. When correlating gene expression with CNA results, only genes that were marked differential using a Wilcoxon rank-sum test via presto with logFC > 0.5 and Benjamin-Hochberg adjusted p<0.01 were used. Correlations were then calculated using Pearson correlation with Bonferroni-corrected p-values. Statistical methods for other analyses are described in-depth in their appropriate sections.

## Acknowledgements

This work was generously supported by the Accelerating Medicines Partnership® Rheumatoid Arthritis and Systemic Lupus Erythematosus (AMP® RA/SLE) Network. AMP is a public-private partnership (AbbVie, Arthritis Foundation, Bristol-Myers Squibb Company, Foundation for the National Institutes of Health, GlaxoSmithKline, Janssen Research and Development, LLC, Lupus Foundation of America, Lupus Research Alliance, Merck & Co., Inc., National Institute of Allergy and Infectious Diseases, National Institute of Arthritis and Musculoskeletal and Skin Diseases, Pfizer, Inc., Rheumatology Research Foundation, Sanofi and Takeda Pharmaceuticals International, Inc.) created to develop new ways of identifying and validating promising biological targets for diagnostics and drug development. Accelerating Medicines Partnership and AMP are registered service marks of the US Department of Health and Human Services.

## Funding

National Institutes of Health UH2-AR067676 (AMP RA/SLE)

National Institutes of Health UH2-AR067677 (AMP RA/SLE)

National Institutes of Health UH2-AR067679 (AMP RA/SLE)

National Institutes of Health UH2-AR067681 (AMP RA/SLE)

National Institutes of Health UH2-AR067685 (AMP RA/SLE)

National Institutes of Health UH2-AR067688 (AMP RA/SLE)

National Institutes of Health UH2-AR067689 (AMP RA/SLE)

National Institutes of Health UH2-AR067690 (AMP RA/SLE)

National Institutes of Health UH2-AR067691 (AMP RA/SLE)

National Institutes of Health UH2-AR067694 (AMP RA/SLE)

National Institutes of Health UM2-AR067678 (AMP RA/SLE)

## Author Contributions

Conceptualization: DAR, AF, TME, J. Gunthridge, PJH, MD, DW, DLK, KCK, RF, MB, PI, RC, DH, ESW, WA, MAM, J.Grossman, CC, HP, JLB, FP-S, CCB, JBH, CP, JHA, SR, NH, JAJ, AD, MAP, JPB, BD

Histology: JBH

scRNA and snRNA Data: TME, NH, MP, RR

Single-cell pipeline: SG, JM, QX, AA, SS, SR

Statistical Analysis: SG, NWS, MC, YZ, SS, SR, AF

Histological Analysis: BR, AF

Website: NWS

Supervision: SR, BD, JLB, JPB, MAP, AD, DAR

Writing – Original Draft: SG, NWS, MC, SR, BD

Writing – Review and Editing: All authors contributed

AMP RA/SLE Network members contributed to this work by managing patient recruitment, curating clinical data, obtaining and processing synovial tissue samples, managing biorepositories, conducting histologic or computational analysis, providing software code, providing website support and/or providing input on data analysis and interpretation.

## Code availability

The code for analyzing this dataset can be found on our GitHub: https://github.com/immunogenomics/AMP_SLE_Kidney.

## Data availability

In accordance with AMP RA/SLE policies the processed count matrices will be made available to the public via the ARK Portal (https://arkportal.synapse.org/) once this study has been accepted for publication in a peer-reviewed journal.

The results published here are in whole or in part based on data obtained from the ARK Portal (http://arkportal.synapse.org). The Accelerating Medicines Partnership® RA/SLE Network data used for this publication are available under:

1. scRNAseq (https://arkportal.synapse.org/Explore/Datasets/DetailsPage?id=54406421)
2. snRNAseq (https://arkportal.synapse.org/Explore/Datasets/DetailsPage?id=54407831)

The ARK Portal hosts data generated by a network of research teams working collaboratively to deepen the understanding of Arthritis and Autoimmune and Related Diseases. It was established by the National Institute of Arthritis and Musculoskeletal and Skin Diseases (NIAMS) and includes data from the Accelerating Medicines Partnership® (AMP®) RA/SLE program.

The specific data used in this publication is available as a controlled-access dataset. Researchers seeking to use these data must submit:

1. A detailed intended data use statement
2. A completed and signed data use certificate

These access requirements ensure responsible and ethical data sharing within the research community.

Instructions for access are available at https://help.arkportal.org/help/data-use-certificate#DataUse&Acknowledgement-Acknowledgement.

An interactive portal with marker gene visualization is available on our website: https://immunogenomics.io/ampsle2/app/?ds=b_pbmcs.

## Supplementary Materials

## Supplementary Methods/Materials

### Outcomes

Complete response (CR) required: 1) UPCR<0.5; and 2) normal creatinine (≤ 1.3 mg/dL) or, if abnormal, ≤ 125% of baseline; and 3) prednisone ≤ 10 mg/day at the time of the study visit. Partial response required: 1) >50% reduction in UPCR; and 2) normal creatinine (≤ 1.3 mg/dL) or, if abnormal, ≤ 125% of baseline; and 3) prednisone dose ≤ 15 mg/day at the time of the study visit. Patients who did not achieve a CR or PR at the specific timepoints were considered non-responders (NR) or not determined (ND) if data were missing. These response definitions were based on the ACCESS Trial (*114*). In agreement with the ACCESS trial, we specifically decided not to include the microscopic review of the urine sediment given the absence of uniformity across sites in assessing urinary sediment and the challenge of attribution especially in a population of young women. The prednisone threshold for CR at ≤ 10 mg prednisone was also based on the ACCESS trial. However, the ≤ 15 mg prednisone maximum for defining PR was agreed upon unanimously by the site investigators.

Although proteinuria was measured by either a UPCR on a spot urine or a timed urine collection, consistency of the method across the study for an individual was required. While determination from a timed urine collection was preferred, if this method was not performed at all time points for an individual participant, calculations from a spot urine were utilized.

### Human Kidney Tissue Collection and Disassociation

Case samples were collected via needle biopsy at the time of their initial visit as part of the AMP program. Control samples were taken from living donors following kidney removal and perfusion. Both case and control kidney biopsies were processed and cryopreserved in batches of 3 following a previously described protocol(*23*) with modifications. Cryovials containing tissue samples were warmed in a 37 C water bath until partially thawed. The contents of each vial were then decanted into one well of a 24 well plate containing 500uL warm RPMI (ThermoFisher) supplemented with 10% FBS (Avantor) and 10mM HEPES (ThermoFisher). Using tweezers, each biopsy was then transferred to a second well in a 24 well plate containing 2mL warm RPMI/FBS/HEPES and allowed to rest for 1 minute, after which it was placed in 100uL of RPMI supplemented with 10 mM HEPES and 0.04% BSA (Millipore Sigma) in a Petri dish. The biopsies were cut into segments approximately 2 mm in length using a clean razor blade. For biopsies analyzed by single nucleus RNA-seq, two noncontiguous segments were removed for nuclei isolation. The remainder were enzymatically dissociated as follows. Biopsy segments, along with the 100uL RPMI/BSA/HEPES, were transferred to a 1.5 mL conical tube containing DMEM/F12 (Corning) and 0.5 mg/mL Liberase TL (Roche) pre-warmed to 37 C. Samples in digestion medium were then agitated for 6 minutes using an orbital shaker set to 300 rpm and 37 C. Samples were then gently mixed 5 times by pipetting with a large-bore tip (Rainin), after which they were returned to the shaker for a second 6 minute 300 rpm/37 C incubation. After the second incubation, samples were placed on ice. 600 uL cold RPMI/FBS/HEPES was added to stop digestion. The remaining tissue fragments were gently resuspended, and then decanted onto a 70 um strainer (Miltenyi) sitting on a 15 mL conical tube on ice. 4 mL cold RPMI/BSA/HEPES was used to rinse the digestion tube and was strained through the 70 um filter to collect single cells released during digestion. The biopsy fragments were then mechanically dissociated by pushing them against the strainer mesh with a cell strainer pestle (CELLTREAT). The pestle and strainer were then rinsed with a further 5 mL cold RPMI/BSA/HEPES to collect dissociated cells. The strained samples were centrifuged at 200 rcf for 8 minutes at 4C, after which cell pellets were resuspended in 50 uL cold RPMI/BSA/HEPES and filtered again through a 40 um strainer (Pluriselect).

Cells in the resulting suspension were counted by trypan blue exclusion.

### Single cell RNA-seq Library Preparation and Sequencing

Single cell RNAseq was performed on the Chromium platform (10x Genomics), using the single cell gene expression 3’ V3 kit. Up to 10,000 live cells from each kidney sample were loaded into separate 10x channels. Complementary DNA amplification and library construction were carried out by the Broad Institute Genomics Platform according to the manufacturer’s instructions.

To reduce sequencing requirements, we removed DNA fragments derived from mitochondrial transcripts using CRISPR-Cas9. We modified the DASH(*115*) method for compatibility with 10x libraries to reduce mitochondrial content. We selected a panel of 61 guide RNA sequences (**Table S15**) to tile regions of the mitochondrial genome heavily represented in 3’ 10x libraries from kidney samples using on-target and off-target predictions from the Broad Institute Genetic Perturbation Platform algorithm(*116*, *117*). We used two non-targeted mitochondrial genes: MT-ND5 and MT-ND6 to estimate the mitochondrial content per cell. We find that MT-ND5 and MT-ND6 correlate to total mitochondrial read content in our pilot data (R=0.982 and R=0.861, respectively) **(Fig. S38A, B).** The summation MT-ND5 + MT-ND6 also correlates to overall mitochondrial read content **(Fig S38C).** A 1% increase in MT-ND5 or MT-ND6 corresponded with a 12.7% and 13.7% increase in total mitochondrial content. A 1% increase in the summation MT-ND5 + MT-ND6 corresponds to a 7.3% increase. We removed cells with more than 3% read contributions of total reads by these two mitochondrial genes. For single nuclei, we used a stricter threshold and removed nuclei with higher than 1% contribution to total reads of total mitochondrial reads. Sensitivity analysis of our mitochondrial threshold in scRNAseq suggests that a 1% change in MT-ND5 and MT-ND6 resulted in a 10% change in cell count **(Fig. S38D).**

T7 reverse transcriptase template was generated from oligonucleotides (IDT) as previously described(*115*). Transcription was performed with 100U T7 RNA polymerase (New England Biolabs) in a 30 uL reaction containing 1X RNAPol Reaction Buffer (NEB), 75ng template DNA, and 0.83 mM each of rATP, rCTP, rGTP and rUTP (NEB). T7 reactions were incubated for 4 hours at 37C. Template DNA was then removed by digestion with 4U DNAse I (NEB) for 15 minutes at 37C. RNA was then purified using a Monarch® RNA Cleanup Kit (NEB), quantified by absorbance of 260 nm light, and aliquoted prior to storage at −80C. Cas9:sgRNA complexes were formed by mixing 1.5 pmoles Cas9 nuclease (NEB) with 1.5 pmoles of the sgRNA pool in a 15 uL solution containing 2uL 10 X NEBuffer™ r3.1 (NEB) and incubating at 25C for 15 minutes. 10ng of individual samples of sequencing-ready 10x gene expression libraries in 5uL water were then added to the ribonucleoprotein complexes and incubated for 2h at 37C to make double strand breaks.

To enable direct amplification of the digested libraries(*118*), the samples were then incubated with 1ug RNAseA (Ambion) and 0.8U Proteinase K (NEB) for 15 minutes at 37C to remove RNP complexes, followed by a second incubation for 15 minutes at 95C to inactivate the enzymes. The libraries were then amplified by 6 PCR cycles using P5/P7 oligonucleotides (AATGATACGGCGACCACCGA, CAAGCAGAAGACGGCATACGA) and KAPA HiFi HotStart ReadyMix (Roche). This step enriched the samples for intact uncut DNA fragments with non-mitochondrial inserts. The PCR products were purified using SPRIselect paramagnetic beads (Beckman Coulter) at a 0.8x volumetric ratio. Finally, the purified libraries were quantified and sequenced according to manufacturer’s guidelines on a Novaseq S4 (Illumina).

### Single nuclear RNA-seq Library Preparation and Sequencing

Nuclei were isolated from a subset of the cryopreserved human kidney biopsies used for single cell RNA-seq using a protocol previously described(*119*), with modifications. We found that ∼4mm from each 18G biopsy provided enough cells or nuclei for transcriptional profiling. Therefore we selected biopsies that were >7mm in length to perform both single cell and single nucleus sequencing. Two non-contiguous ∼2mm sections of each biopsy were placed in a 2mL dounce tissue grinder (Kimble) containing cold lysis buffer (20mM Tris pH7.5, 1M sucrose, 5mM CaCl2, 3mM MgCl2, 1mM EDTA, 0.1% Triton-X100). Biopsy fragments were homogenized with 5 strokes using pestle A and 20 strokes with pestle B, then incubated for 10 minutes on ice. The sample was then strained through a 30um mesh (Celltrix) and diluted with 6mL cold PBS with 1mM EGTA. Nuclei were then pelleted by centrifugation at 500g for 5 minutes at 4C. Supernatant was removed, and the pellet resuspended in 100uL cold PBS + 1% BSA (manufacturer) and a 1:1,000 dilution of RNase inhibitor (Takara). The resulting nuclei suspension was then strained through a 10um mesh (Pluriselect) before loading up to 8,000 nuclei for gene expression profiling using the Chromium 3’ V3 kit (10x genomics).

### PBMC Collection and Processing

Blood was collected in heparin tubes and processed within 2 hours. PBMC were isolated by centrifugation through a density gradient medium in 50mL SepMate tubes (Stemcell Technologies) following manufacturer guidelines, and then cryopreserved in CryoStor CS10 (Stemcell Technologies). Frozen samples were shipped to Oklahoma Medical Research Foundation Biorepository for centralized storage. Samples were randomized and shipped frozen to the Broad Institute where they were processed in batches of up to 16 to generate single cell RNAseq data. Samples were thawed at 37 degrees C, then diluted in cold RPMI 1640 (Gibco) supplemented with 0.5% BSA (Miltenyi Biotec), 1X Glutamax (Gibco), and 10mM HEPES (Corning). Samples were thereafter maintained on ice or at 4 degrees C whenever possible. Cells were filtered through a 40um strainer (Pluriselect), then pelleted by centrifugation. Pellets were resuspended in autoMACS Running Buffer (Miltenyi Biotec) containing human FcR Blocking Reagent (Miltenyi Biotec) and fluorescent antibodies (Biolegend) targeting CD15 (clone W6D3 conjugated to BV510) and CD45 (clone HI30 conjugated to one of: FITC, PE, PE/Cy7, AF700), then incubated for 20 minutes on ice. Samples were then washed, and live cells were counted using ViaStain AOPI Solution (Nexcelom Bioscience) and a Cellaca MX instrument (Nexcelom Bioscience). Pools of 4 samples were created, with the CD45 fluorochrome unique to a single sample in each pool. Each pool contained 600,000 live cells. Pools were incubated for 30 minutes at 4 degrees C in a volume of 50uL Cell Staining Buffer (Biolegend) containing oligo barcode-tagged antibodies for surface protein detection (Totalseq-A Human Universal Cocktail, V1.0 from Biolegend; ¼ test). Pools were then washed twice and resuspended in autoMACS Running Buffer containing 1mg/mL DAPI (Biolegend) before filtration through a 40um strainer. Equal numbers of live, CD45 positive, CD15 negative cells from each PBMC donor were FACS sorted (Sony MA900) into RPMI 1640 (Gibco) supplemented with 0.04% BSA (Miltenyi Biotec), 1X Glutamax (Gibco), and 10mM HEPES (Corning). Cells derived from each donor in the pool were distinguished by the CD45 antibody fluorochrome. After sorting, all samples in the batch were pooled, and 32,000 cells were loaded in each of 3 Chromium chips (10x Genomics) to generate single cell gene expression libraries using manufacturer’s protocols. The libraries were sequenced on Novaseq S4 flow cells (Illumina) to an average depth of 60,000 reads per cell.

### PBMC processing, quality control, and profiling

We processed PBMCs from raw 10X read data with cellranger pipeline (version XXX). Briefly, we generated fastq files from the bcl file output from the 10X sequencer using the cellranger makefastq function and generated bam files from fastq files using the cellranger count function. We employed cellranger version 6.1.1 for this preprocessing. PBMC samples were not hashed for sample of origin and required computational matching. We employed demuxlet(*120*) to match samples to donors using genotyping information collected at time of enrollment to the AMP study. We removed cells which could not be attributed to a single donor confidently via demuxlet’s functionalities. Next, we imported the data into the Seurat package, and using Seurat quality control functionalities we removed: cells with less than 500 genes per cell and over 20% contribution to total reads from mitochondrial reads.

We then normalized and scaled the PBMC dataset using the Seurat pipeline’s inbuilt functions and performed PCA. We used the Seurat implementation of harmony to mitigate non-biological effects from the batch of origin and sample. We employed the Seurat implementations of the UMAP algorithm for low dimensional visualization and clustered the harmonized pcs using Louvain clustering at low resolution. We further employed Scrublet to identify inter cell-type doublets and removed clusters with high scrublet scores and expression of marker genes from multiple cell types. We repeated the PCA, harmonization, and clustering procedures post doublet removal.

We used commonly identified markers of broad immune cell types to define PBMC transcriptomic profiles post clustering in the whole PBMC space. We used *CD4, CD8*, *and CD3* for T Cellls, *CD56* and *TYROBP* for NK Cells, *CD14, CD16,* and *CCR2/L2* for monocytes. We further used *MS4A1* and *CD19* for B cell identification. We sourced markers for cell states from a previously described study of memory t cells in tuberculosis patients, the AMP Phase I LN and RA studies, the AMP Phase II RA study, and the KPMP project. Within cell types, we applied the same strategy used for tissue cells to eliminate likely doublet and low-quality clusters, and defined cell states with the sources and software packages described above.

To understand the relationship between tissue and blood immune cell states, we also sequenced 327,326 PBMCs from 119 LN patients and 19 healthy controls. We defined broad cell types using marker genes such as *CD3D* for T cells, *CD14* for monocytes, *MS4A1* for B cells, *NCAM1* for NK cells, and *CD1C* for dendritic cells (**Fig. S9B**).

### Processing and Quality Control for Kidney Transcriptomics Data

We converted raw sequencing reads to count matrices with cell ranger (version 5.0.1), first converting sequencing output to FASTQ files using the cellranger mkfastq function, We aligned reads to GRCh38 and generated count matrices with cellranger count. Quality control for the scRNAseq and snRNAseq data involved several common steps in single cell/nuclear transcriptomic sequencing analysis. Initially, we filtered cells removing cells and nuclei with less than 1000 reads per cell or nucleus, and less than 500 unique genes detected. Since we employed DASH to reduce mitochondrial read contribution to the sequencing, we could not define mitochondrial read content per cell or nucleus with the entire list of mitochondrial genes. However, we used two non-dashed mitochondrial genes: *MT-ND5* and *MT-ND6* to estimate the mitochondrial content per cell and nucleus. We removed cells with more than 3% read contribution total reads by these two mitochondrial genes. For single nuclei, we used a stricter threshold and removed nuclei with higher than 1% contribution to total reads by these two mitochondrial genes.

### Defining Broad Renal Cell Types and Doublet Removal

We employed standard scRNAseq and snRNAseq preprocessing steps. We first transformed the raw counts matrix by multiplying with a factor of 10^4^ and log normalizing the count. We then scaled the raw counts to have mean zero and variance 1. We were concerned that the list of variable genes would be dominated by the scRNAseq modality, which had many more samples. Additionally, we did not want to obtain variable genes from samples with low cell or nuclear count. We therefore selected the top 500 most variable genes based on dispersion from each sample with more than 25 cells or nuclei within each technology. The final list of variable genes was obtained by taking the union of these sets obtained from these patients. We were additionally concerned about uneven contributions to low dimensional embeddings from each modality due to imbalanced sample numbers. We therefore adapted the weighted PCA approach described in Zhang. et al (*121*) and initially characterized in Korsunsky et. al (*28*) to balance the contribution from single cell and single nuclear cells when computing principal components. We then applied harmony to these principal components to correct for non-biological variation from processing batch, sample collection site, sequencing modality, and samples.

Using the harmonized principal components, we created reduced dimensional representations for visualization using the UMAP algorithm(*122*). We additionally employed density-based Louvain clustering on fuzzy shared nearest neighbor graphs computed using the harmonized principal components. We clustered at low resolution to obtain broad cell type groupings. We were concerned about the possibility of doublets formed by multiple cell types, so we employed Scrublet(*123*) to identify possible sources of doublets. After manual inspection, we removed clusters of cells expressing markers for multiple cell types and possessing a qualitatively higher-than-average Scrublet score. After manual inspection, we also removed clusters of low-quality cells, marked by higher-than-average mitochondrial content and a lower number of genes and reads per cell.

To define major immune and tissue cell types, we employed markers identified by several previous works, including the Kidney Precision Medicine Project(*36*), the AMP Phase II Rheumatoid Arthritis publication(*124*), and the AMP Phase I LN publication. Broadly we employed known markers for immune cells, including: *CD3, CD4,* and *CD8* for T cells, *CD56* and *TYROBP* for NK cells, *CD14* and *CD16* for macrophages, and *XCR1, SDC1, MS4A1,* for B and Plasma cells. For tissue cell types, we used kidney specific marker genes such as: *CUBN* and *SLC34A1* for proximal tubules, *SLC12A1* and *CRYAB* for cells from the loop of henle, *CLNK, SLC12A3, AQP2,* and *SLC8A1* for cells from the distal nephron, *PECAM1, COL1A1,* and *NOTCH3* for cells from the endothelial/stromal compartment, and *CFH* and *PODXL* from cells from the glomerulus.

### Blood-Tissue Immune Integration

Tissue and blood immune cells were integrated into a single embedding using weighted PCA to account for the larger amount of tissue-origin immune cells than PBMC. Variable genes were found independently between the tissue and the blood and the union was used to compute the principal components. Further batch correction was done through Harmony with theta=0 with a maximum of 50 iterations to mitigate non-biological variation arising from factors such as individual sample differences, processing batches, and the original source of the cells (tissue or blood). Only samples present in the blood and tissue were taken.

The Tissue Specificity Metric (TSM) was calculated per-cell on the basis that, for a given cell, what fraction of its 50 nearest neighbors are from the tissue over the total number of cells. The numerator of the TSM calculation represents the number of these 50 nearest neighbors that originate from the same tissue as the index cell. The denominator of the TSM equation is designed to normalize this count using a weighted factor that accounts for the inherent differences in the abundance of different cell types, particularly the often disproportionately high number of immune cells present in blood compared to tissue cells. A high TSM value for a given cell indicates that its local microenvironment is predominantly composed of cells from the same tissue, suggesting a strong tissue-specific identity. Conversely, a low TSM value suggests that the cell’s neighborhood is more blood-like in nature. A TSM score of 0.5 indicates that a cell’s neighborhood reflects what the expected proportion of blood to tissue cells should equal mixing occur. The TSM can be given through the equation:

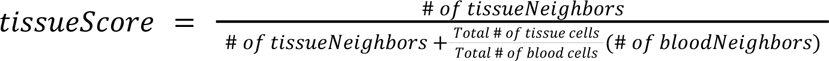

Groups with a median score of above 0.7 were assigned as tissue-specific, while a median score of below 0.3 was designated as blood-specific. All other populations are demarcated as being shared.

### Pseudotime Analysis

Pseudotime lineages were computed by embedding PBMC and tissue cells into a joint embedding using Destiny (*41*) on harmonized Principal Components (PCs). Trajectories were subsequently computed using Slingshot (*42*). Start points were chosen from PBMC clusters that were categorized as blood-specific by TSM.

### Constructing an Interferon Stimulated Gene Score

We constructed a per-cell score reflecting relative expression of interferon stimulated genes using a previously published gene list(*23*) and the commonly used method for single-cell scoring with a gene set used in Seurat’s AddModuleScore and Scanpy’s score_genes functions (*125*, *126*). Briefly, within each cell type for scRNAseq tissue data, we performed library size normalization (TPM) and log+1 transformation of the single cell raw counts matrices, where log refers to the natural log. We then performed z-score scaling of each gene and applied Seurat’s AddModuleScore() function using a set of known interferon stimulated genes. For calculating sample level correlation with chronicity and activity index, we averaged the calculated score across all immune (B/plasma, myeloid, and T/NK) or all tissue (glomerular, proximal tubule, distal nephron, loop of Henle, endothelial/stromal) cell types.

### Receptor-Ligand Analysis

Cell–cell communication analysis was performed using the CellChat R package(*127*). We performed ligand receptor analysis subsetting only to immune and glomerular cells from scRNAseq experiments. We use the full human ligand-receptor interaction database provided to CellChat.

Overexpressed genes and ligand–receptor interactions were identified using CellChat’s native identifyOverExpressedGenes and identifyOverExpressedInteractions functions. Communication probabilities between cell groups were inferred using the computeCommunProb function with the tri-mean method. Inferred interactions were aggregated into signaling networks using aggregateNet.

### SAM Spatial Logistic Regression Classifier

We build our logistic regression classifier using the glmnet package in R. We designate M5, M7, M9, and M11 as SAMs and all other myeloid cells as non-SAMs. Training data was selected from a random 90% of samples with the remaining 10% withheld for testing. Featurized genes were selected from the overlap between the AMP and Danaher et al. dataset and further subsetted to include genes that were expressed in at least 25% of genes in spatial transcriptomic cells. Cross fold validation was performed across all the potential folds and the first fold was arbitrarily selected. To optimize regularization, we performed additional cross-validation using cv.glmnet to optimize lambda. The trained model was applied to all myeloid cells, and cells were classified as “SAMs” if predicted probability >0.8, “non-SAMs” if <0.2, and “ambiguous” if probabilities fell between these thresholds.

---

**Fig. S1.**
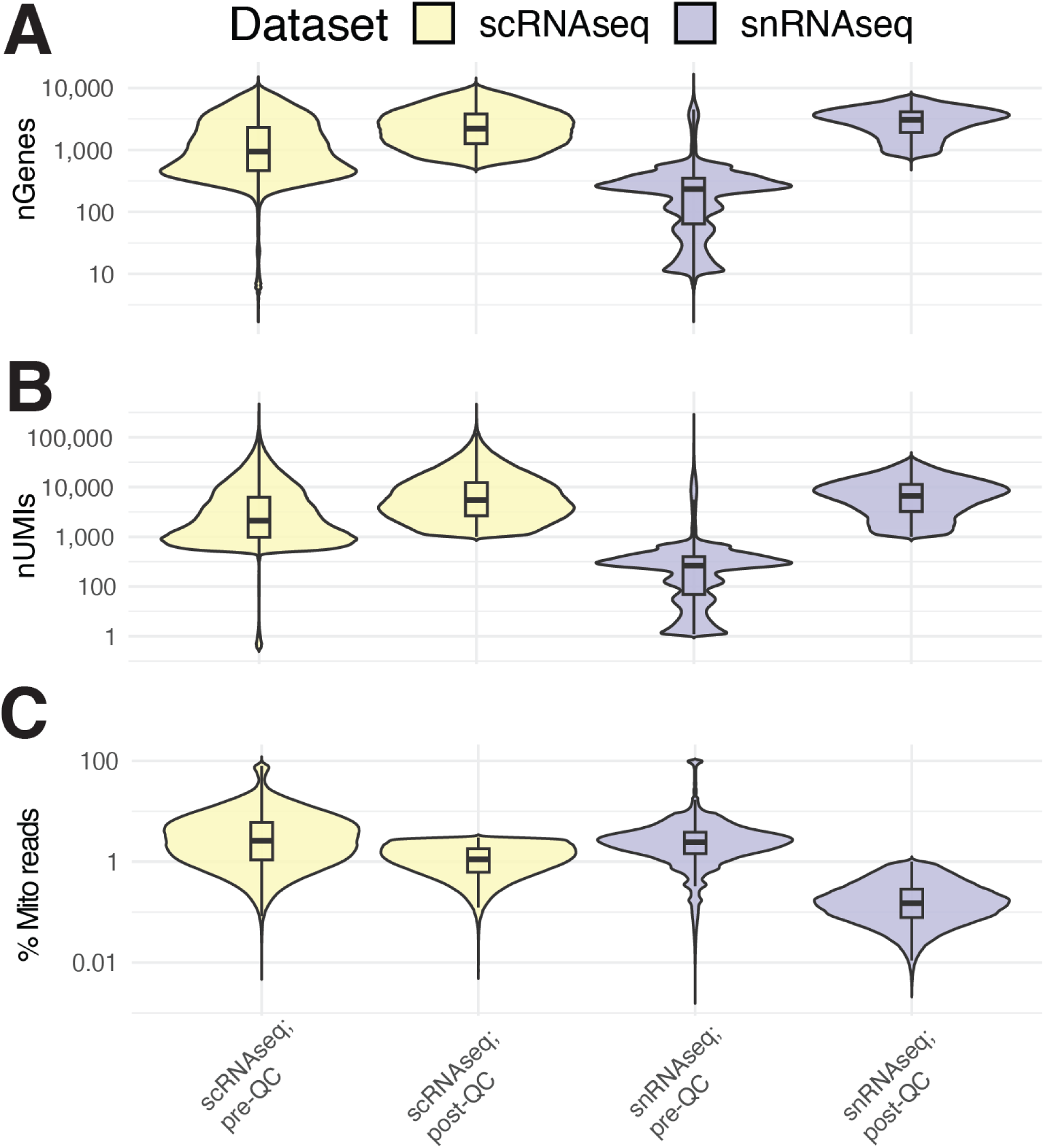
Histogram of scRNAseq and snRNAseq prior to quality control. Histograms of uniquely expressed genes (top), UMIs (center), and % mitochondrial reads (bottom) stratified for scRNAseq (A) and snRNAseq (B) modalities. Vertical lines correspond to post quality control cutoffs. Lines are drawn at > 500 genes, > 1,000 UMIs, and >3% MT reads in scRNAseq and between 500 and 7500 uniquely expressed genes, between 1,000 and 40,000 UMIs, and less than 1% mitochondrial reads in snRNAseq.

**Fig. S2.**
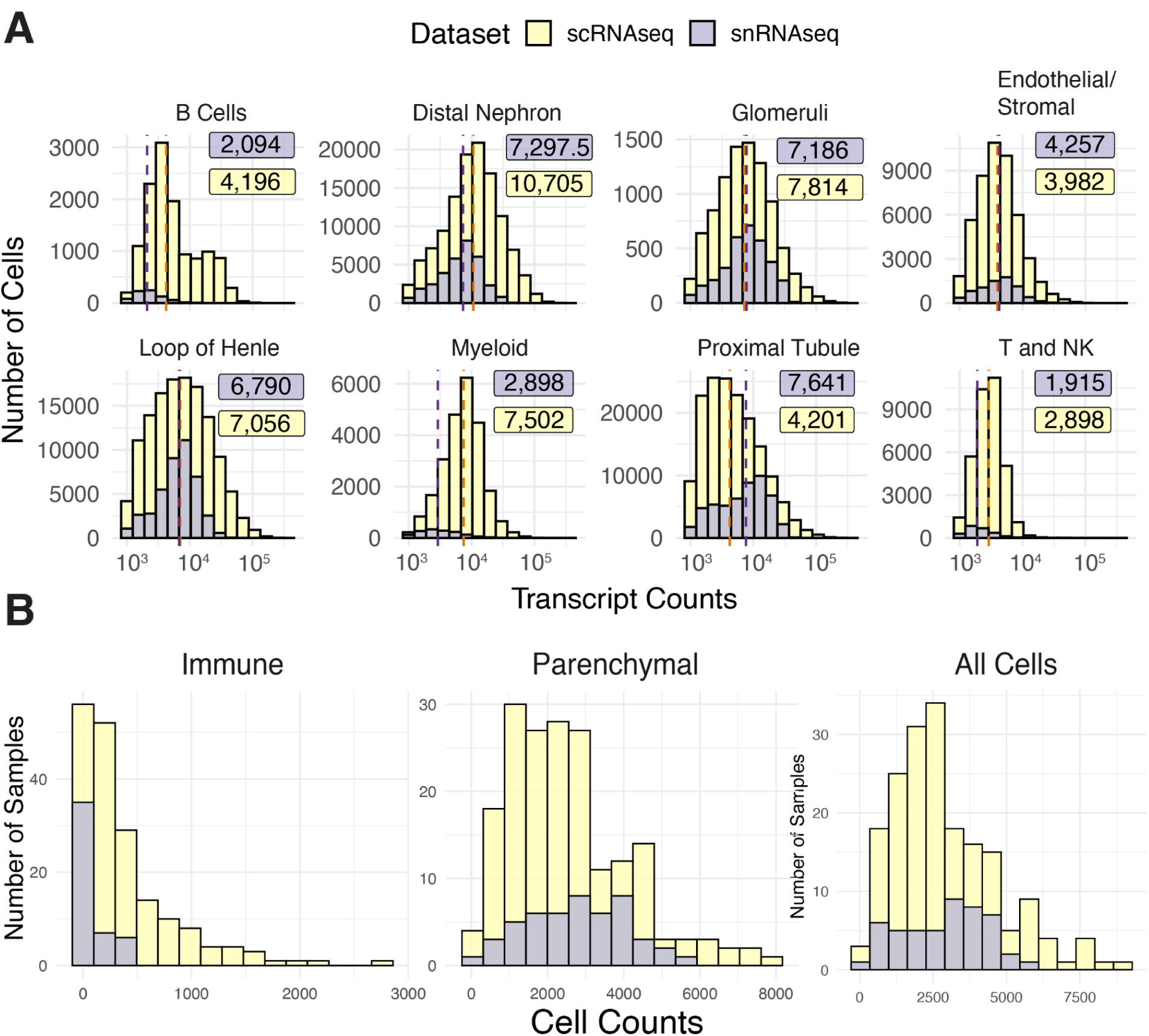
Transcript Count Histograms. (A) Histogram of transcript counts per modality, faceted by cell type. Vertical lines are drawn at each modality’s median in scRNAseq (deep orange) and snRNAseq (deep purple). Medians for scRNAseq (yellow box) and snRNAseq (purple box) are included for each cell type. (B) Histogram of cell counts per sample in each modality, faceted by immune cells (left), parenchymal cells (center), or both immune and parenchymal cells (right).

**Fig. S3.**
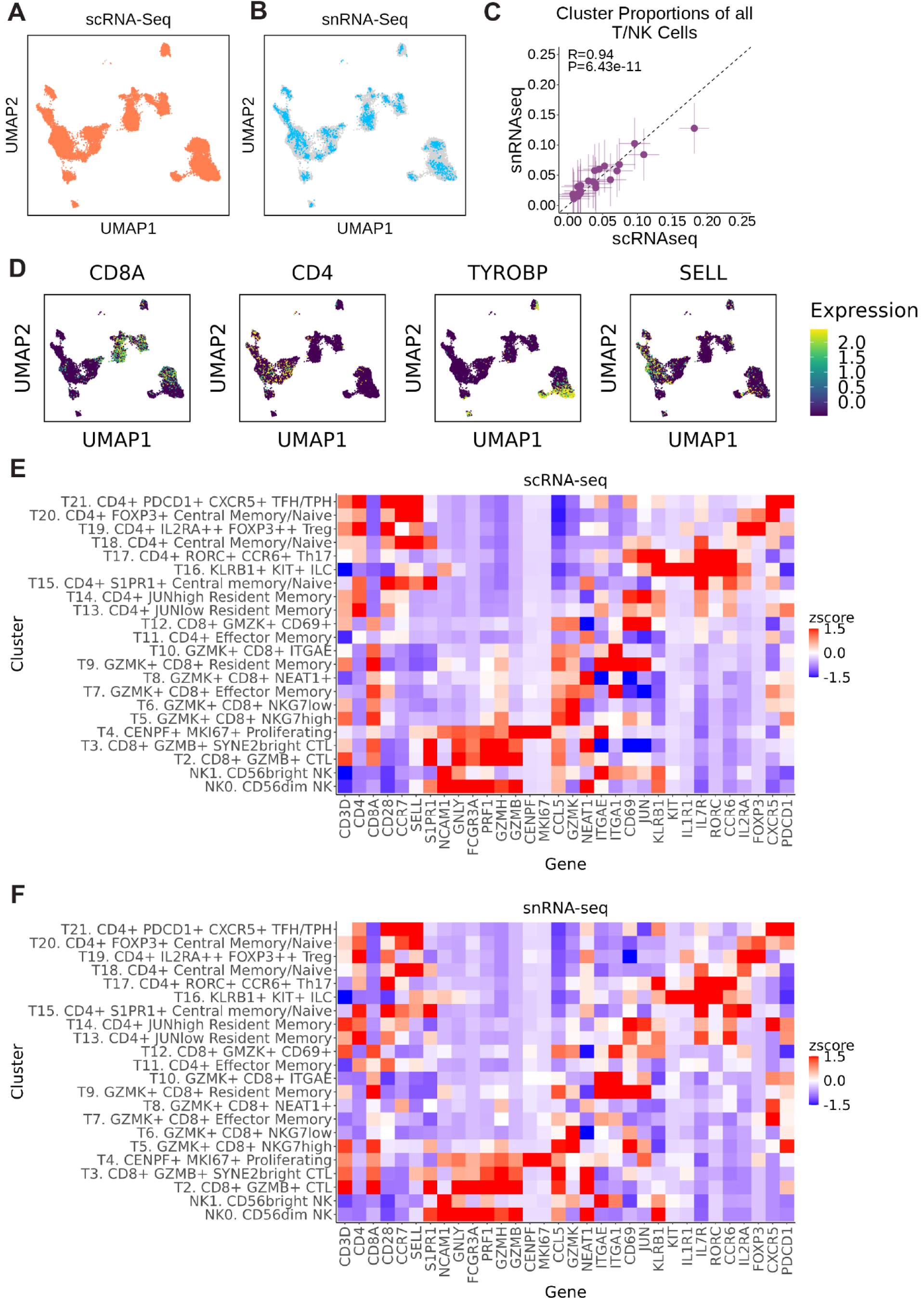
T/NK. **(A)** UMAP depiction of cell states captured from scRNA-seq. (**B)** UMAP depiction of cells from snRNA-seq. (**C)** Scatterplot displaying each cell state cluster’s proportion out of total T/NK cells. X-axis reflects cell state proportions calculated in scRNA-seq and Y-axis reflects snRNA-seq. Proportions were calculated as the proportion of total cell state annotations across all donors with both matched scRNA-seq and snRNA-seq (n=22). Error bars reflect the standard error of that cluster’s cell state proportion across donors, where horizontal lines reflect per-sample variability in scRNA-seq proportions, and vertical lines reflect per-sample variability in snRNA-seq proportions. (**D)** Expression of marker genes (log-normalized, Z-scored) across all T/NK cells. (**E-F)** Heatmaps depicting average expression of marker genes (log-normalized, Z-scored after averaging) within each cell state annotation. Shown in scRNA-seq **(E)** and snRNA-seq **(F)**.

**Fig. S4.**
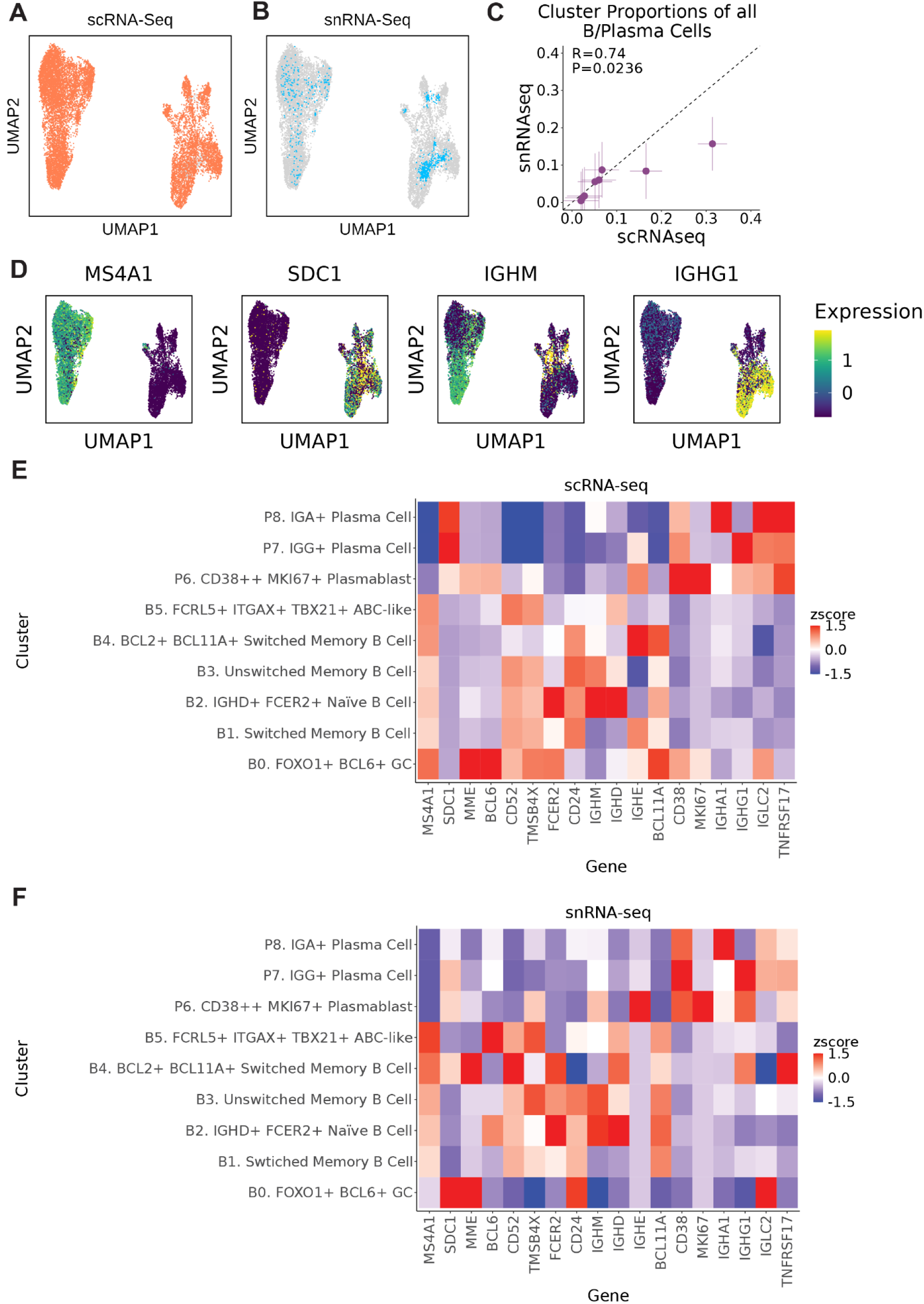
B/Plasma. **(A)** UMAP depiction of cell states captured from scRNA-seq. **(B)** UMAP depiction of cells from snRNA-seq. **(C)** Scatterplot displaying each cell state cluster’s proportion out of total B/Plasma cells. X-axis reflects cell state proportions calculated in scRNA-seq and Y-axis reflects snRNA-seq. Proportions were calculated as the proportion of total cell state annotations across all donors with both matched scRNA-seq and snRNA-seq (n=22). Error bars reflect the standard error of that cluster’s cell state proportion across donors, where horizontal lines reflect per-sample variability in scRNA-seq proportions, and vertical lines reflect per-sample variability in snRNA-seq proportions. **(D)** Expression of marker genes (log-normalized, Z-scored) across all B/Plasma cells. **(E-F)** Heatmaps depicting average expression of marker genes (log-normalized, Z-scored after averaging) within each cell state annotation. Shown in scRNA-seq **(E)** and snRNA-seq **(F)**.

**Fig. S5.**
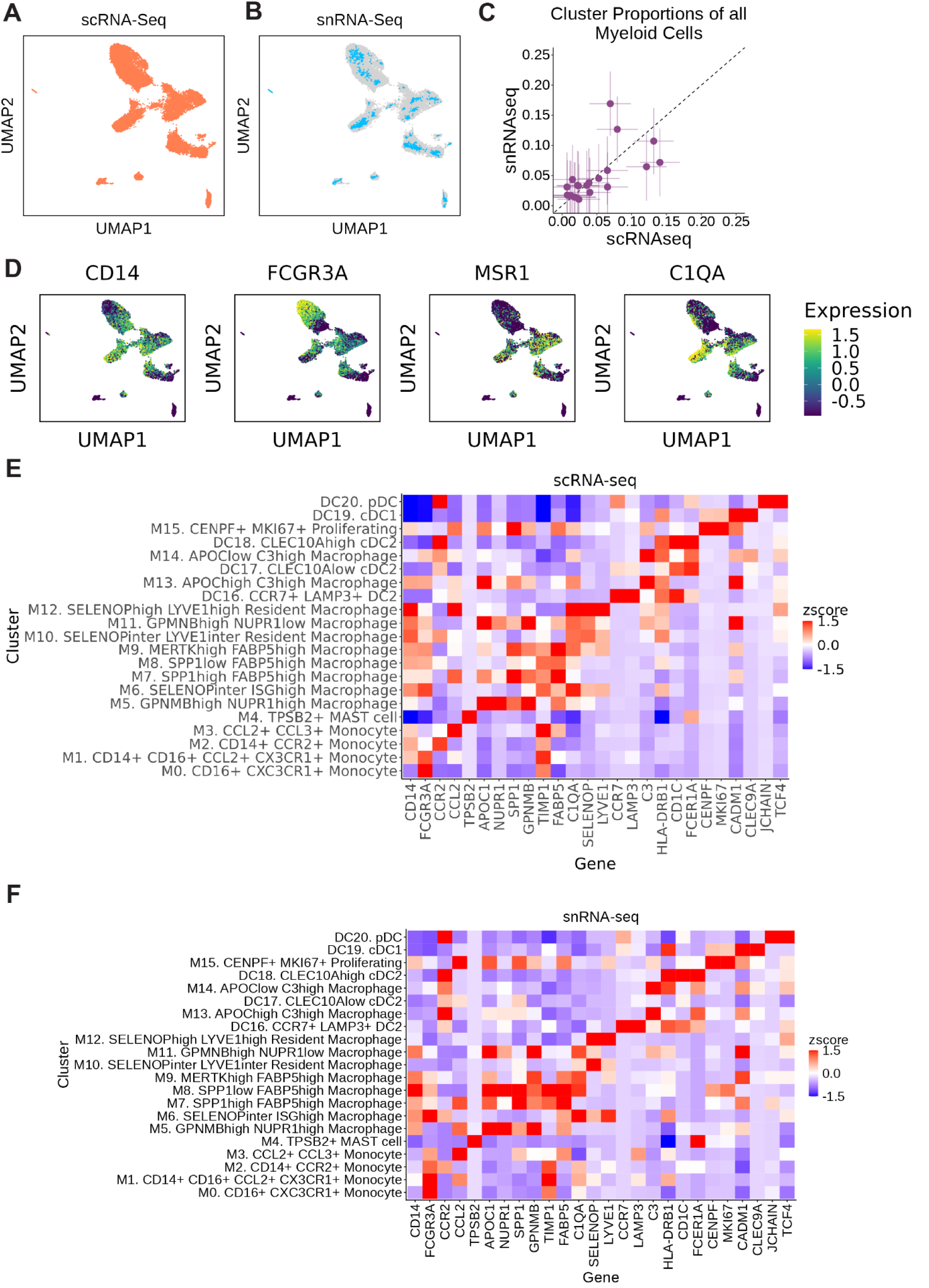
Myeloid. **(A)** UMAP depiction of cell states captured from scRNA-seq. **(B)** UMAP depiction of cells from snRNA-seq. **(C)** Scatterplot displaying each cell state cluster’s proportion out of total Myeloid cells. X-axis reflects cell state proportions calculated in scRNA-seq and Y-axis reflects snRNA-seq. Proportions were calculated as the proportion of total cell state annotations across all donors with both matched scRNA-seq and snRNA-seq (n=22). Error bars reflect the standard error of that cluster’s cell state proportion across donors, where horizontal lines reflect per-sample variability in scRNA-seq proportions, and vertical lines reflect per-sample variability in snRNA-seq proportions. **(D)** Expression of marker genes (log-normalized, Z-scored) across all Myeloid cells. **(E-F)** Heatmaps depicting average expression of marker genes (log-normalized, Z-scored after averaging) within each cell state annotation. Shown in scRNA-seq **(E)** and snRNA-seq **(F)**.

**Fig. S6.**
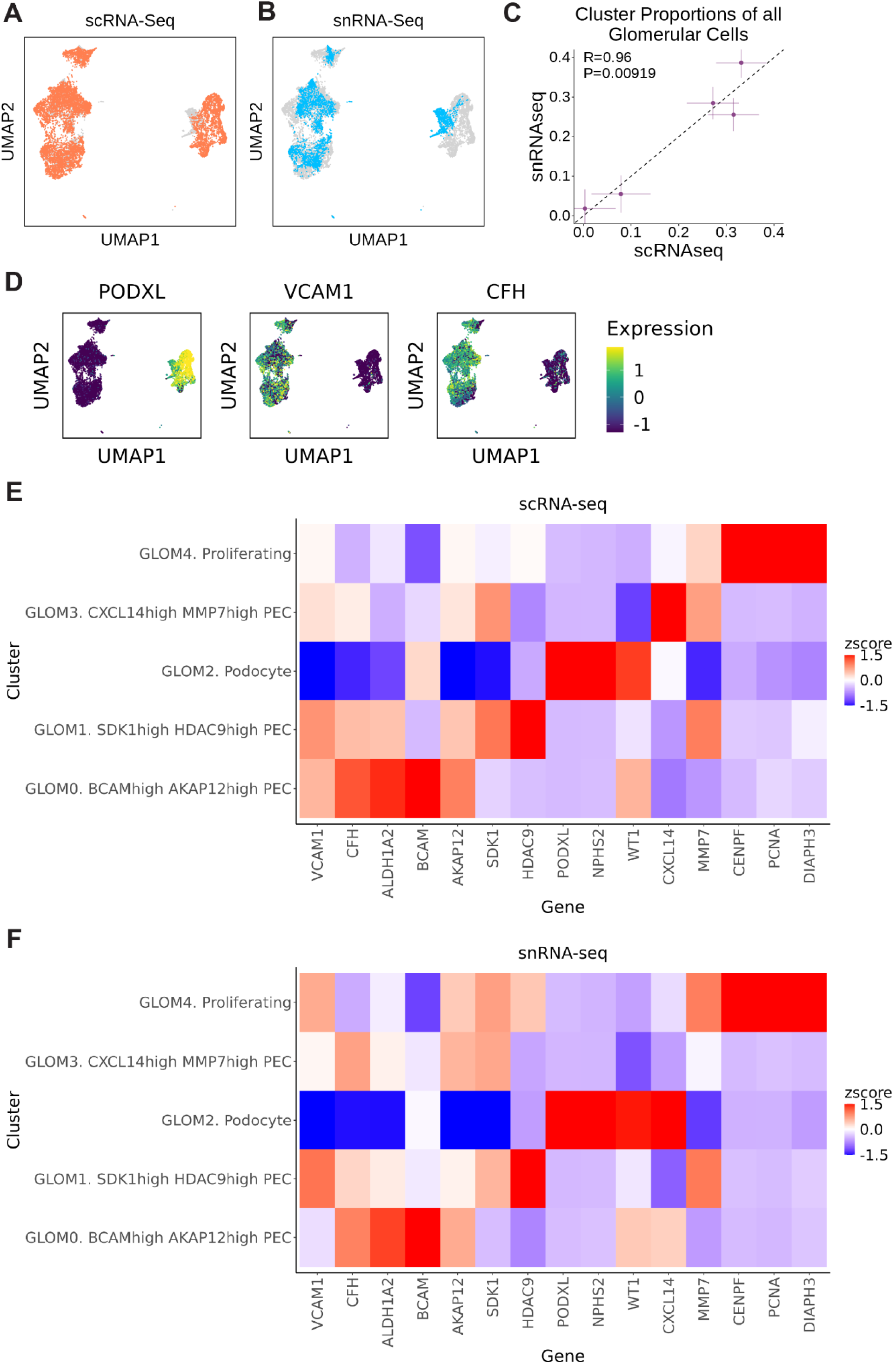
Glomerular. **(A)** UMAP depiction of cell states captured from scRNA-seq. **(B)** UMAP depiction of cells from snRNA-seq. **(C)** Scatterplot displaying each cell state cluster’s proportion out of total glomerular cells. X-axis reflects cell state proportions calculated in scRNA-seq and Y-axis reflects snRNA-seq. Proportions were calculated as the proportion of total cell state annotations across all donors with both matched scRNA-seq and snRNA-seq (n=22). Error bars reflect the standard error of that cluster’s cell state proportion across donors, where horizontal lines reflect per-sample variability in scRNA-seq proportions, and vertical lines reflect per-sample variability in snRNA-seq proportions. **(D)** Expression of marker genes (log-normalized, Z-scored) across all glomerular cells. **(E-F)** Heatmaps depicting average expression of marker genes (log-normalized, Z-scored after averaging) within each cell state annotation. Shown in scRNA-seq **(E)** and snRNA-seq **(F)**.

**Fig. S7.**
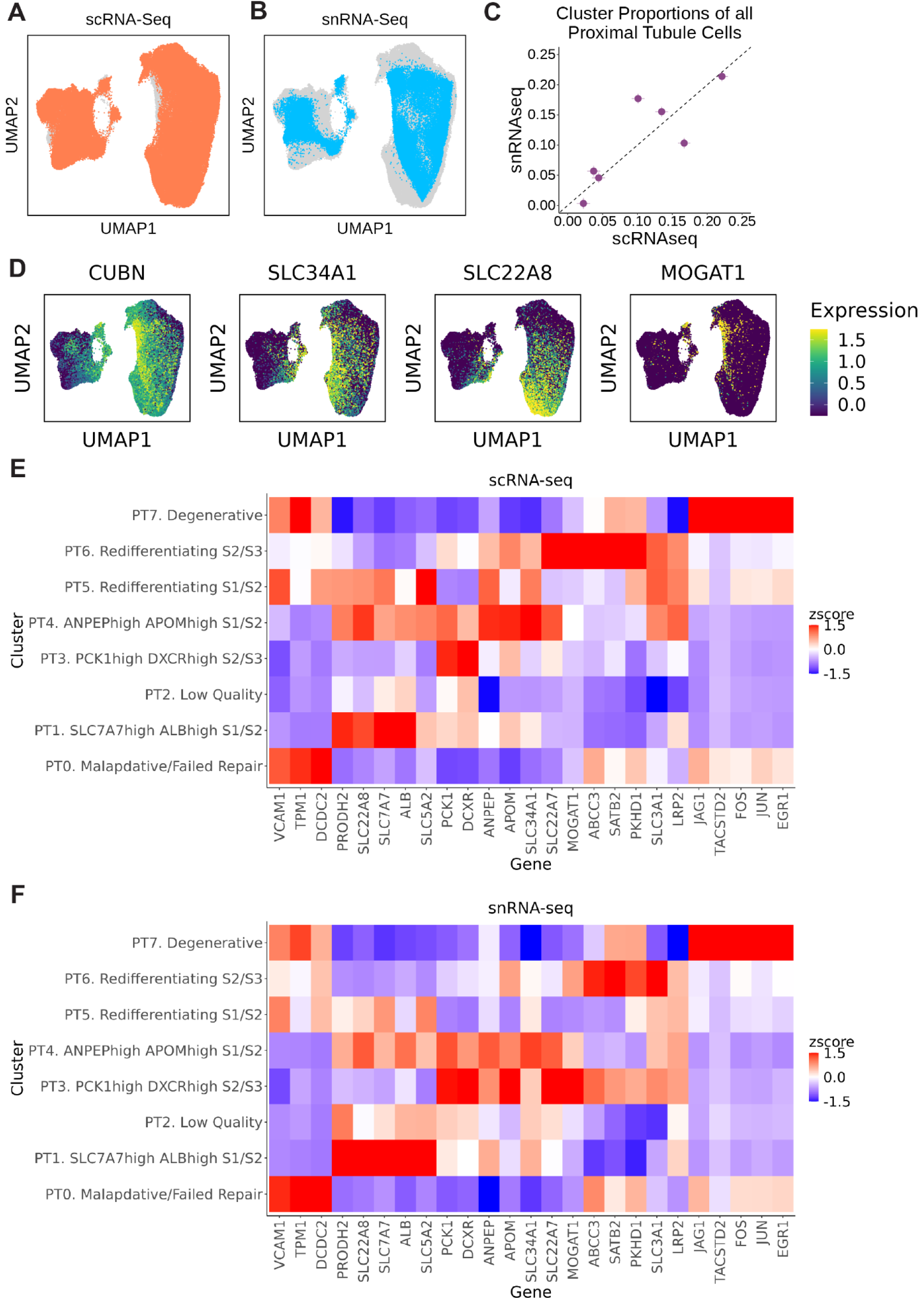
Proximal tubule. **(A)** UMAP depiction of cell states captured from scRNA-seq. **(B)** UMAP depiction of cells from snRNA-seq. **(C)** Scatterplot displaying each cell state cluster’s proportion out of total Proximal Tubule cells. X-axis reflects cell state proportions calculated in scRNA-seq and Y-axis reflects snRNA-seq. Proportions were calculated as the proportion of total cell state annotations across all donors with both matched scRNA-seq and snRNA-seq (n=22). Error bars reflect the standard error of that cluster’s cell state proportion across donors, where horizontal lines reflect per-sample variability in scRNA-seq proportions, and vertical lines reflect per-sample variability in snRNA-seq proportions. **(D)** Expression of marker genes (log-normalized, Z-scored) across all Proximal Tubule cells. **(E-F)** Heatmaps depicting average expression of marker genes (log-normalized, Z-scored after averaging) within each cell state annotation. Shown in scRNA-seq **(E)** and snRNA-seq **(F)**.

**Fig. S8.**
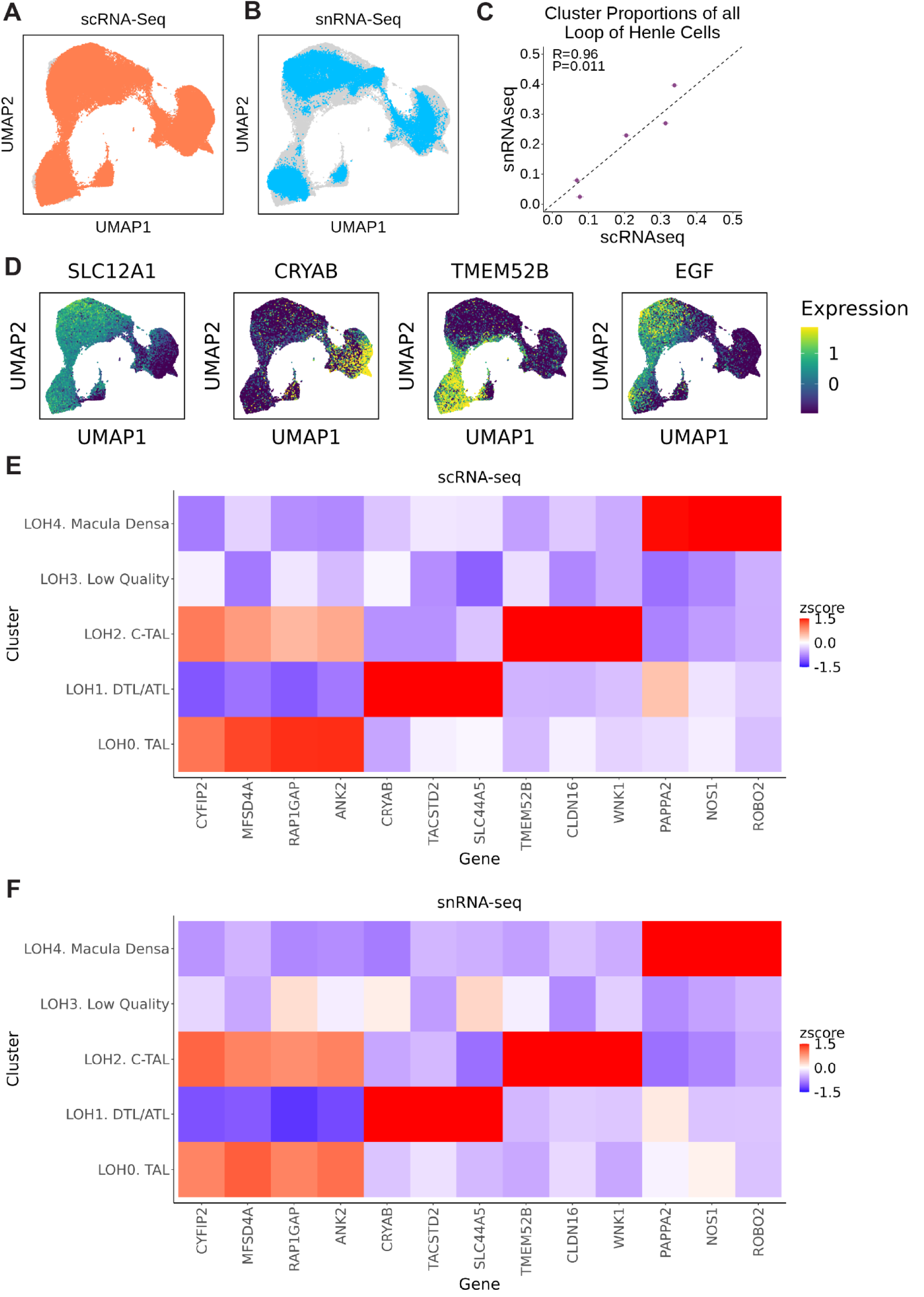
Loop of Henle. **(A)** UMAP depiction of cell states captured from scRNA-seq. **(B)** UMAP depiction of cells from snRNA-seq. **(C)** Scatterplot displaying each cell state cluster’s proportion out of total Loop of Henle cells. X-axis reflects cell state proportions calculated in scRNA-seq and Y-axis reflects snRNA-seq. Proportions were calculated as the proportion of total cell state annotations across all donors with both matched scRNA-seq and snRNA-seq (n=22). Error bars reflect the standard error of that cluster’s cell state proportion across donors, where horizontal lines reflect per-sample variability in scRNA-seq proportions, and vertical lines reflect per-sample variability in snRNA-seq proportions. **(D)** Expression of marker genes (log-normalized, Z-scored) across all Loop of Henle cells. **(E-F)** Heatmaps depicting average expression of marker genes (log-normalized, Z-scored after averaging) within each cell state annotation. Shown in scRNA-seq **(E)** and snRNA-seq **(F)**.

**Fig. S9.**
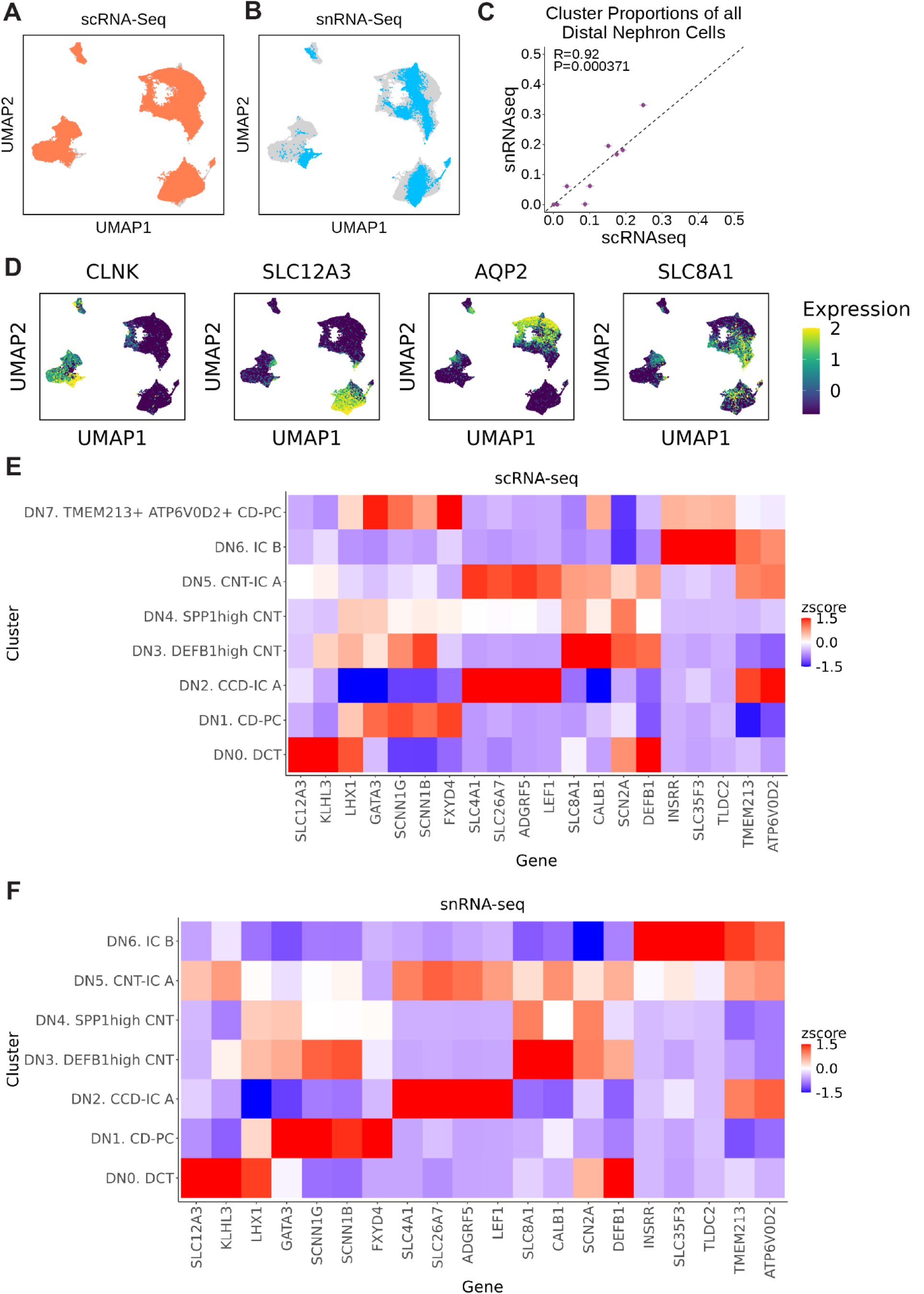
Distal nephron. **(A)** UMAP depiction of cell states captured from scRNA-seq. **(B)** UMAP depiction of cells from snRNA-seq. **(C)** Scatterplot displaying each cell state cluster’s proportion out of total Distal Nephron cells. X-axis reflects cell state proportions calculated in scRNA-seq and Y-axis reflects snRNA-seq. Proportions were calculated as the proportion of total cell state annotations across all donors with both matched scRNA-seq and snRNA-seq (n=22). Error bars reflect the standard error of that cluster’s cell state proportion across donors, where horizontal lines reflect per-sample variability in scRNA-seq proportions, and vertical lines reflect per-sample variability in snRNA-seq proportions. **(D)** Expression of marker genes (log-normalized, Z-scored) across all Distal Nephron cells. **(E-F)** Heatmaps depicting average expression of marker genes (log-normalized, Z-scored after averaging) within each cell state annotation. Shown in scRNA-seq **(E)** and snRNA-seq **(F)**.

**Fig. S10.**
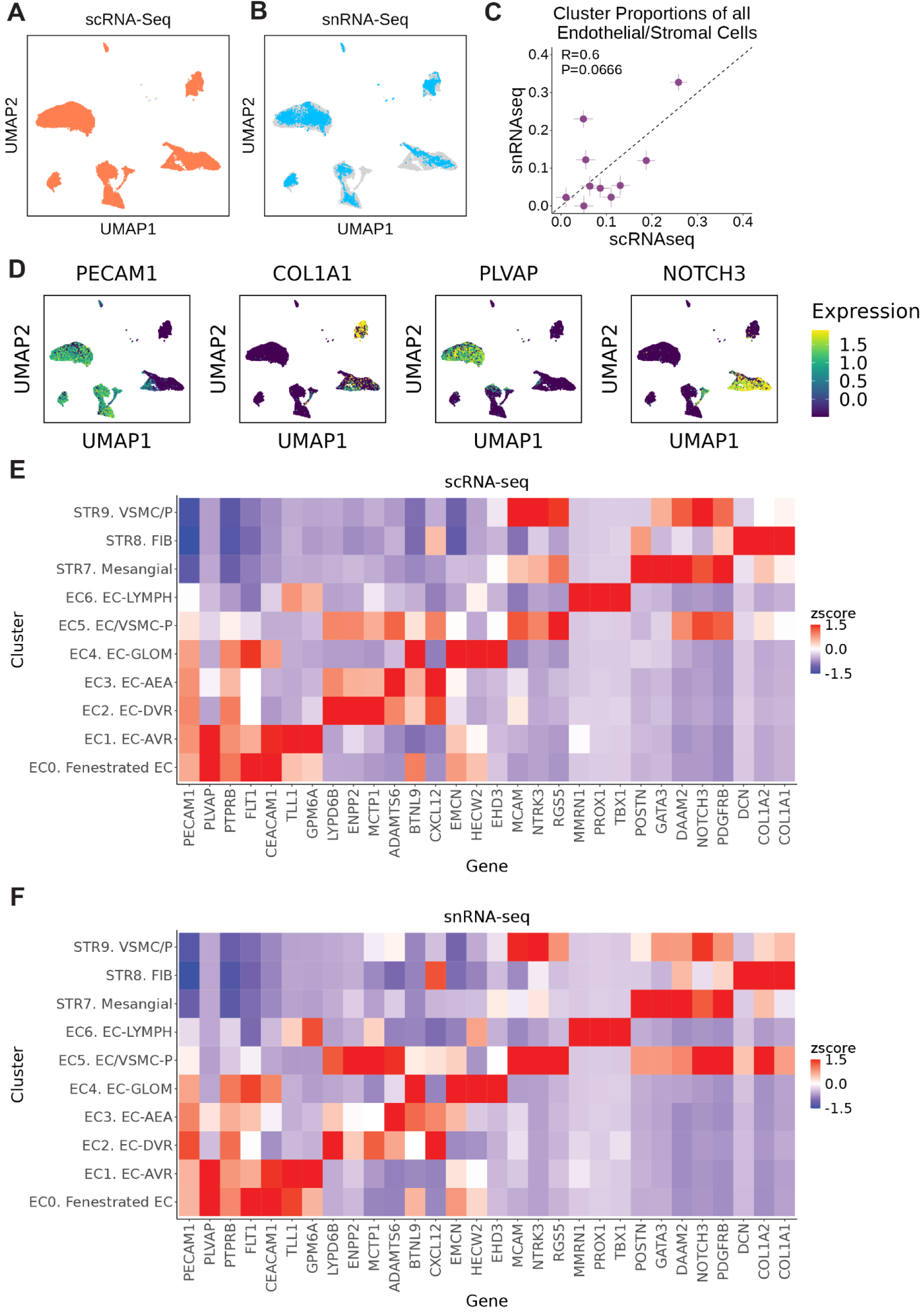
Endothelial/Stromal. **(A)** UMAP depiction of cell states captured from scRNA-seq. **(B)** UMAP depiction of cells from snRNA-seq. **(C)** Scatterplot displaying each cell state cluster’s proportion out of total endothelial/stromal cells. X-axis reflects cell state proportions calculated in scRNA-seq and Y-axis reflects snRNA-seq. Proportions were calculated as the proportion of total cell state annotations across all donors with both matched scRNA-seq and snRNA-seq (n=22). Error bars reflect the standard error of that cluster’s cell state proportion across donors, where horizontal lines reflect per-sample variability in scRNA-seq proportions, and vertical lines reflect per-sample variability in snRNA-seq proportions. (**D)** Expression of marker genes (log-normalized, Z-scored) across all endothelial/stromal cells. **(E-F)** Heatmaps depicting average expression of marker genes (log-normalized, Z-scored after averaging) within each cell state annotation. Shown in scRNA-seq **(E)** and snRNA-seq **(F)**.

**Fig. S11.**
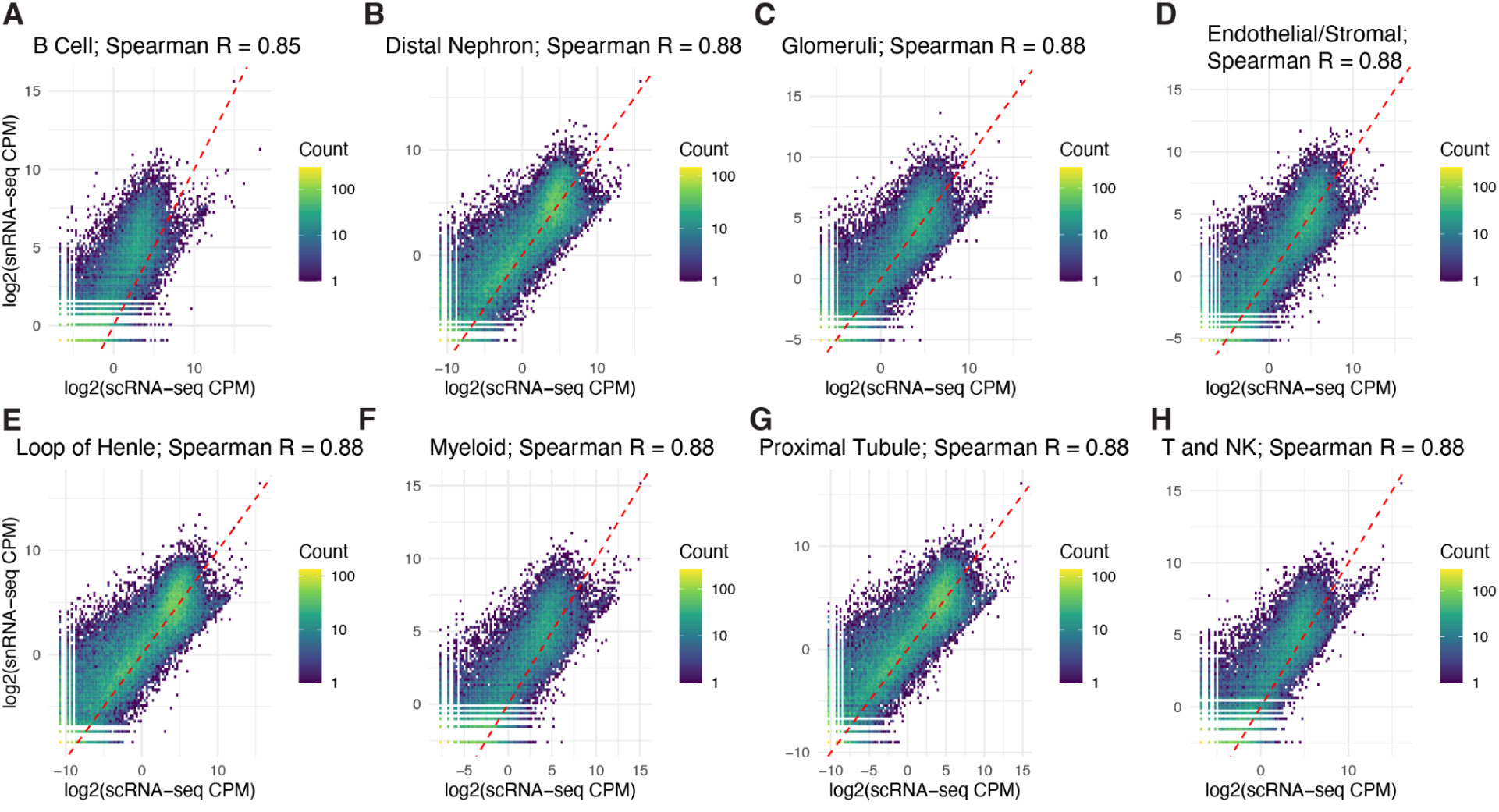
Global transcript correlations within cell type. Density plots of log2 counts per million per gene stratified by B cells **(A),** distal nephron **(B),** glomeruli **(C),** endothelial/stromal cells **(D),** loop of Henle **(E),** myeloid **(F),** proximal tubule **(G),** and T and NK cells **(H).** Spearman correlations are included.

**Fig. S12.**
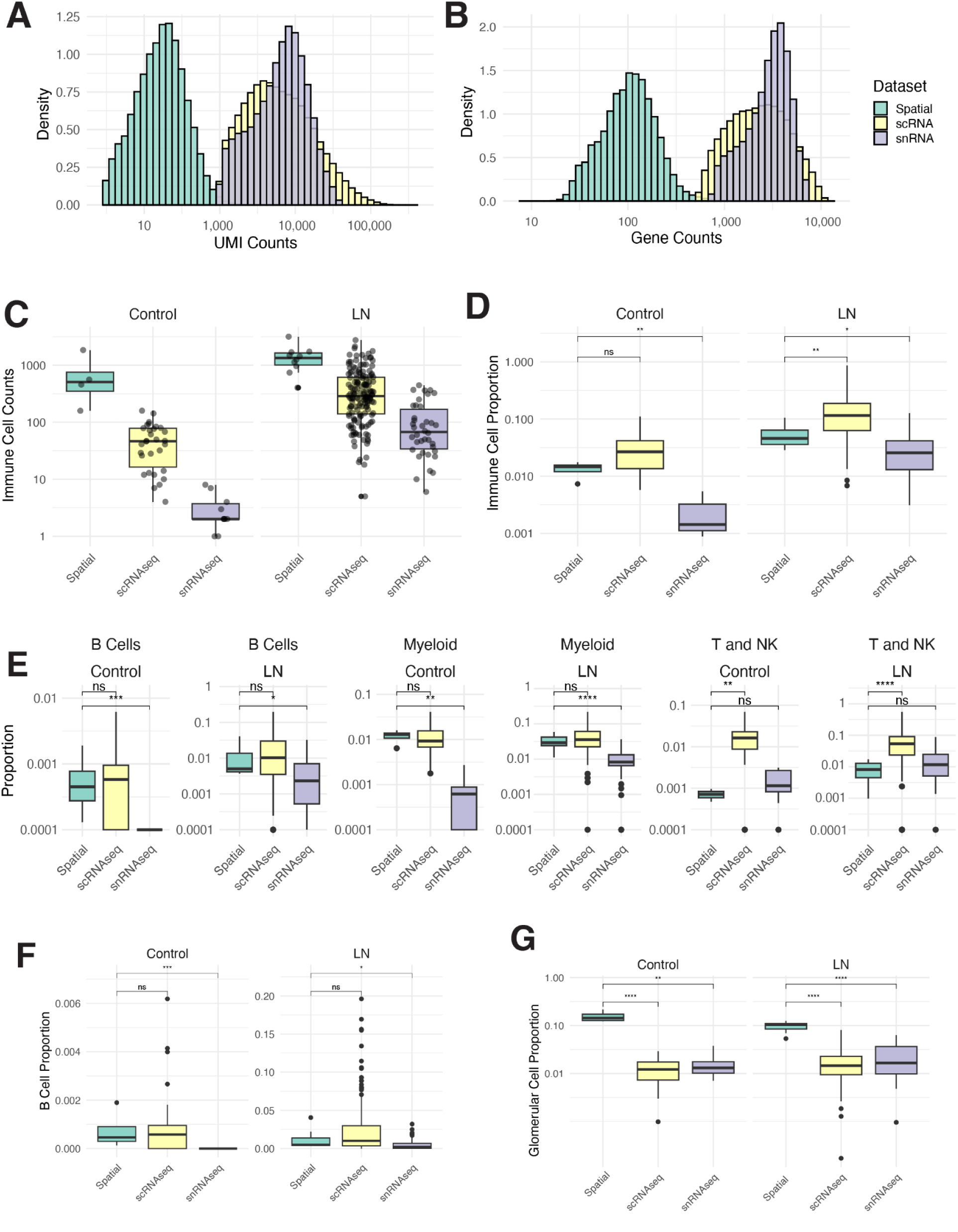
Sparsity of Immune Cells and Glomerular Cells. **(A)** Histogram of total gene counts per cell, stratified by modality. Histograms are normalized such that total area per group is equal. **(B)** Same as **(A)**, but for total genes expressed. **(C)** Total immune cells collected per biopsy, stratified by collection technology. **(D)** Proportion of immune cells per biopsy, stratified by collection technology. **(E)** Proportion of immune cells per biopsy, faceted by B cells (left), myeloid cells (middle) and T and NK cells (right). **(F)** Proportion of B cells of all cells per collection method (G) same as (F), but for glomerular cells as a proportion of all collected cells per sample.

**Fig. S13.**
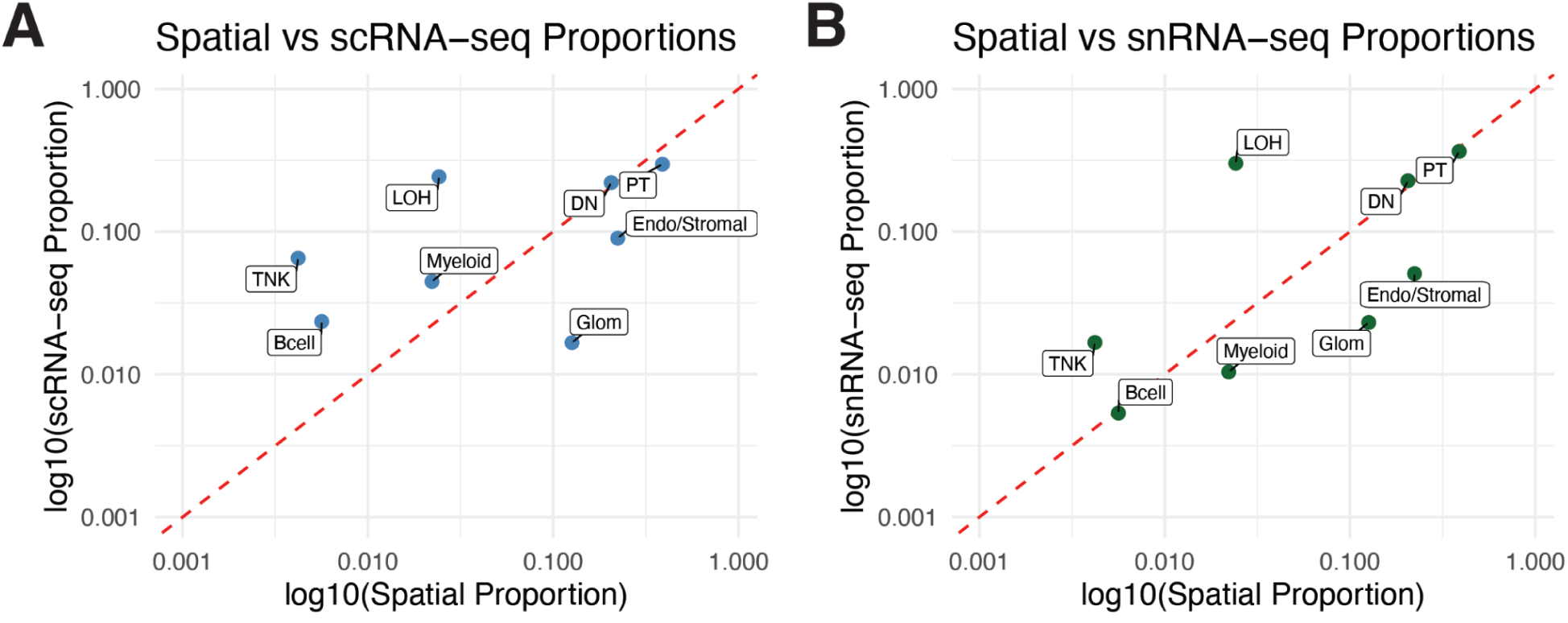
Cell Type Proportions between spatial and single-cell modalities. **(A)** Proportion of broad cell types in Spatial data versus scRNAseq data. **(B)** Proportion of broad cell types in Spatial data versus snRNAseq data.

**Fig. S14.**
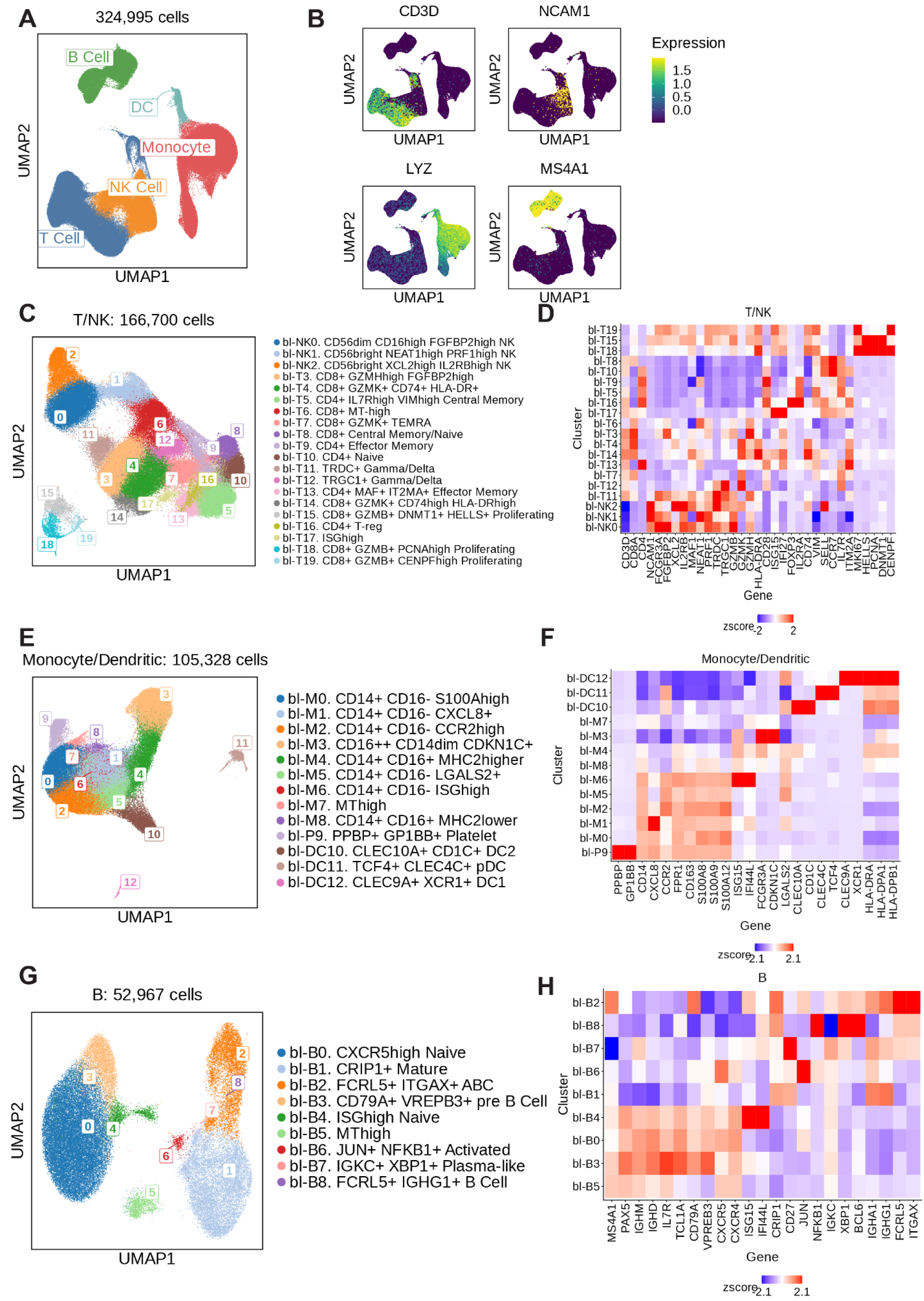
PBMC dataset overview. **(A)** UMAP of all single cells annotated in blood, colored by broad cell type annotation. **(B)** Per-cell gene expression for various broad cell type markers. Gene values are log(TPM+1) normalized and scaled for visualization. **(C)** UMAP of T/NK cell states in blood, colored by fine grain cell state annotations. Annotation boxes are located at the centroid (mean) of each cluster. **(D)** Heatmap depicting marker gene expression in each T/NK cell state cluster. Expression values the log(TPM+1) cluster average of a marker, and z-score scaled by column for visualization. **(E-F),** Same as **(C-D)** for myeloid cells. **(G-H),** Same as **(C-D)** for B/plasma cells.

**Fig. S15.**
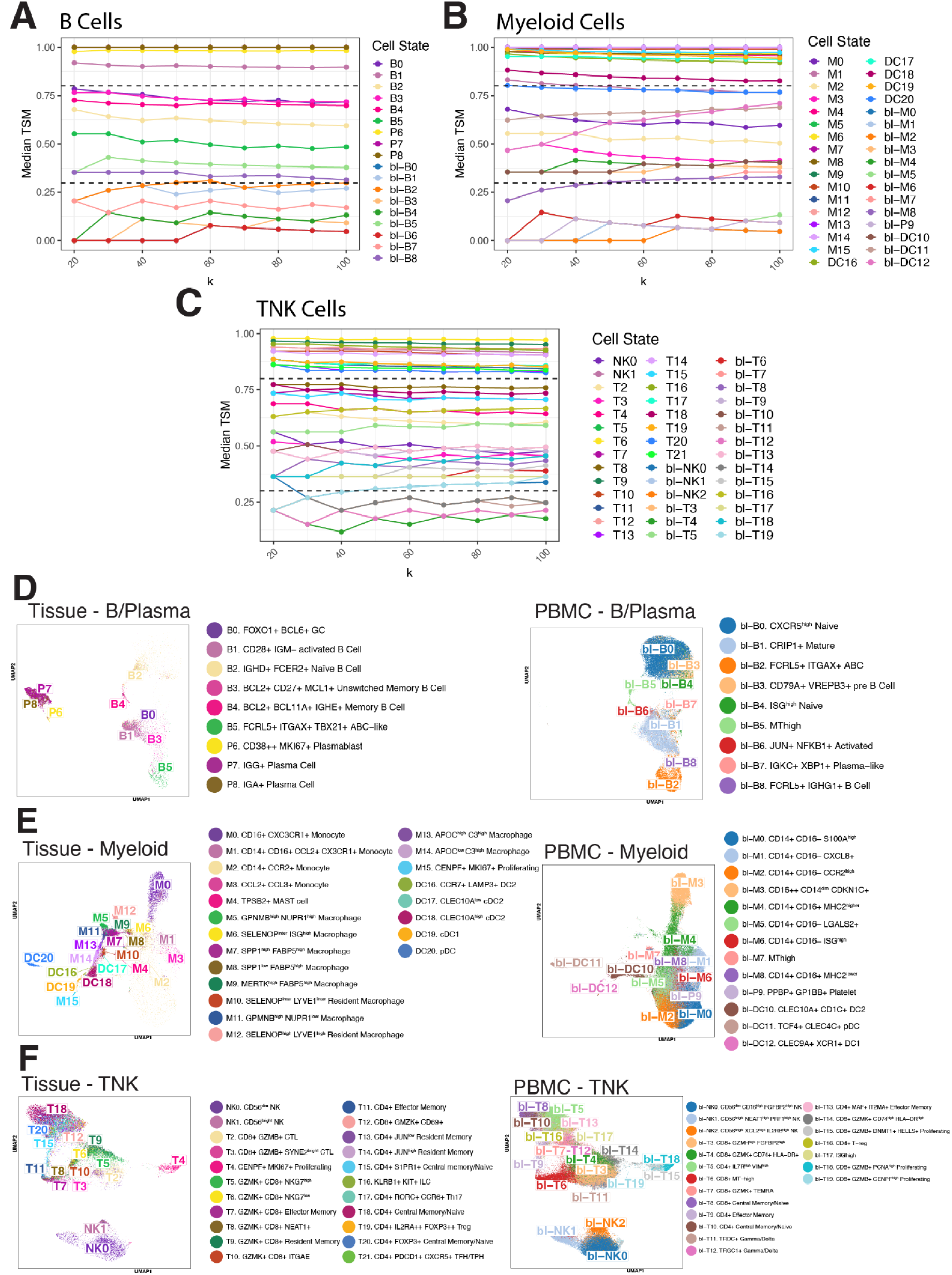
Robustness of TSM. Median TSM scores per cell state for B **(A),** Myeloid **(B),** and TNK **(C)** cell types. Dashed lines denote tissue-specific (> 0.8) and blood-specific (< 0.3) cutoffs. **(D-F)** Joint embedding of tissue (left) and blood (right) immune cells colored by the original clustering for B and plasma cells **(D),** myeloid cells **(E),** and T and NK cells **(F).**

**Fig. S16.**
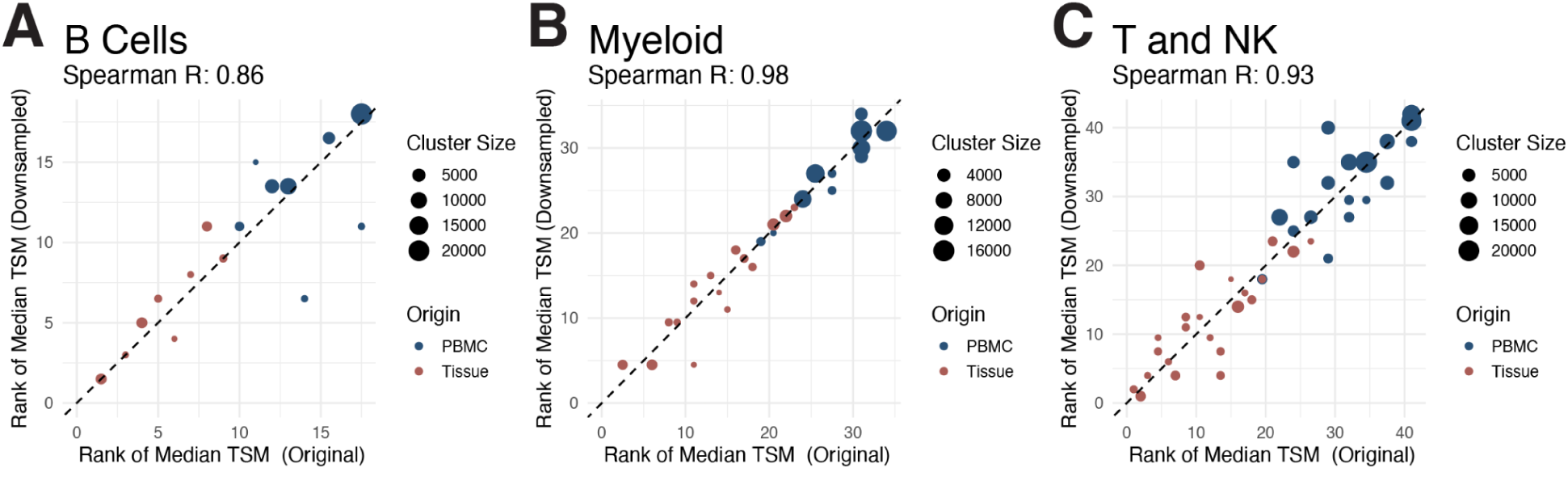
Median cluster TSM rankings before and after downsampling. Correlational plots of per-cluster median TSM rankings for B cells **(A),** Myeloid cells **(B),** and T and NK cells **(C).** Dot sizes are proportional to relative cluster size within that cell type and colored by origin.

**Fig. S17.**
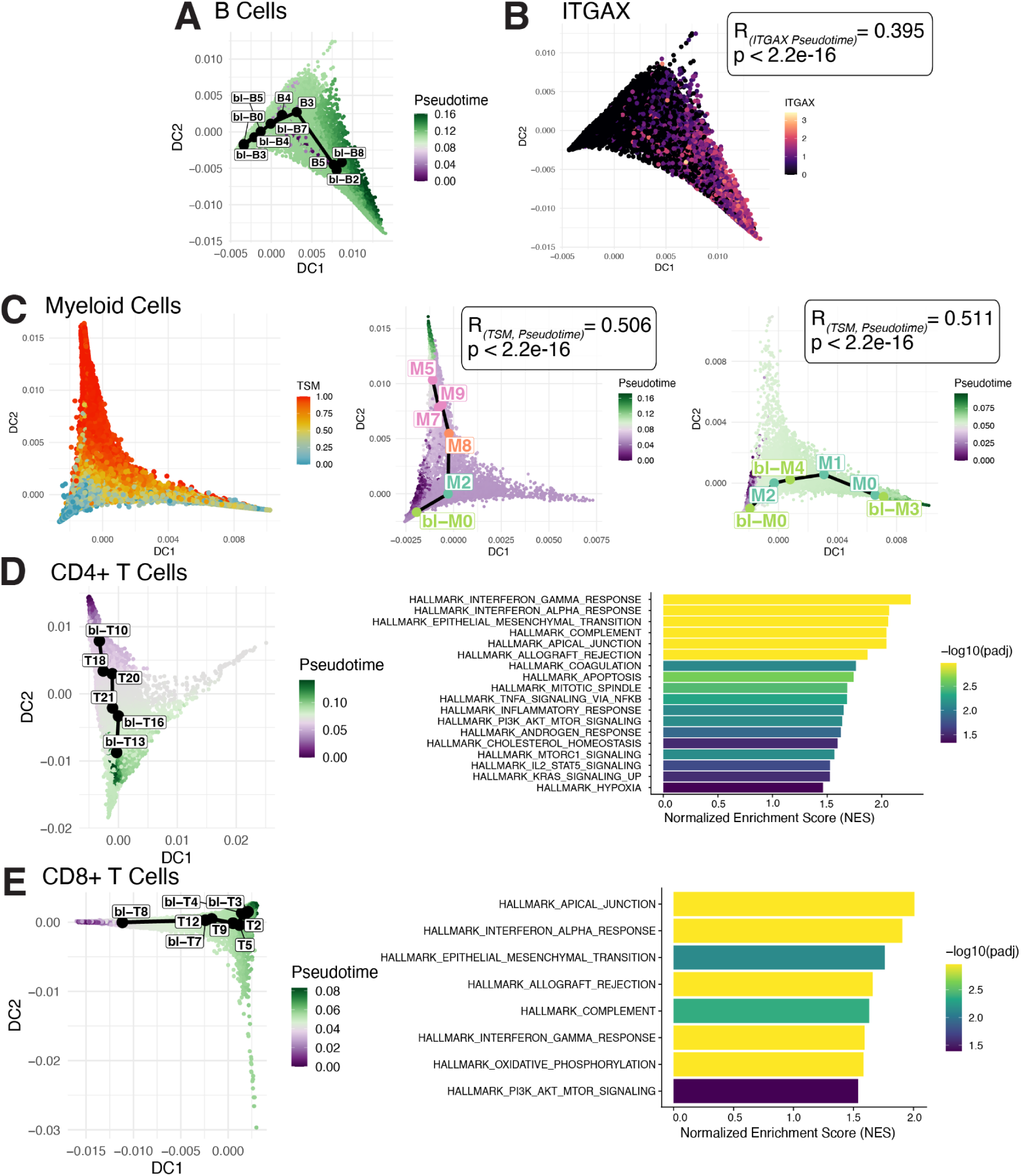
Pseudotime analysis of joint immune cell embeddings. **(A)** Pseudotime trajectory for B cells on destiny embedding. **(B)** ABC-marker ITGAX expression in B cells. Includes R coefficient between ITGAX expression and pseudotime computed by slingshot for the trajectory. **(C)** Left - myeloid cell TSM score colored on destiny embedding. Center - pseudotime trajectory for myeloid cells on destiny embedding. Includes R coefficient between TSM and pseudotime computed by slingshot for trajectory. Right - Same as (center), but for a monocyte-only trajectory. **(D)** Left - pseudotime trajectory for CD4+ T cells. Right - GSEA barplot on genes correlated with pseudotime trajectory. **(E)** Same as **(D)** but for CD8+ T cells

**Fig. S18.**
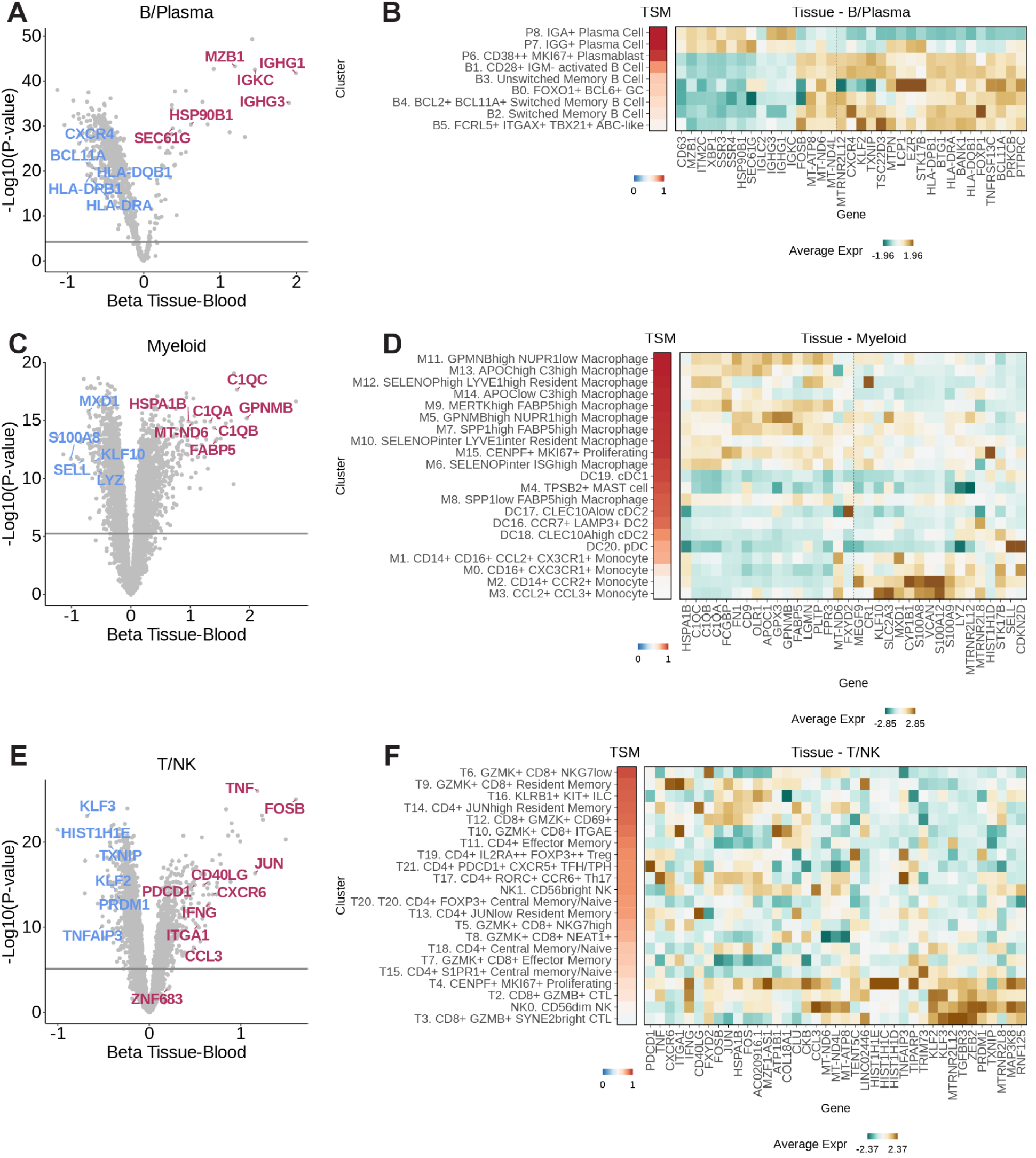
Differential gene expression between immune compartment in blood and tissue. **(A)** In B/plasma cells, differential expression results between tissue and blood samples. Betas and p-values are derived from pseudobulk linear modeling. Genes upregulated with respect to tissue are displayed in red, downregulated genes are displayed in blue. Horizontal lines indicate the Bonferonni corrected significance threshold for p<0.05. **(B)** Heatmap depicting average log(CPM+1) expression of marker genes for each B/plasma cluster in tissue. For visualization, expression values are z-scored by column. Dashed line separates genes identified as enriched (left) and downregulated (right) in tissue via differential expression. Clusters are ordered by TSM score. **(C-D),** Same as **(A-B)** but in myeloid cells. **(E-F),** Same as **(A-B)** but in T/NK cells.

**Fig. S19.**
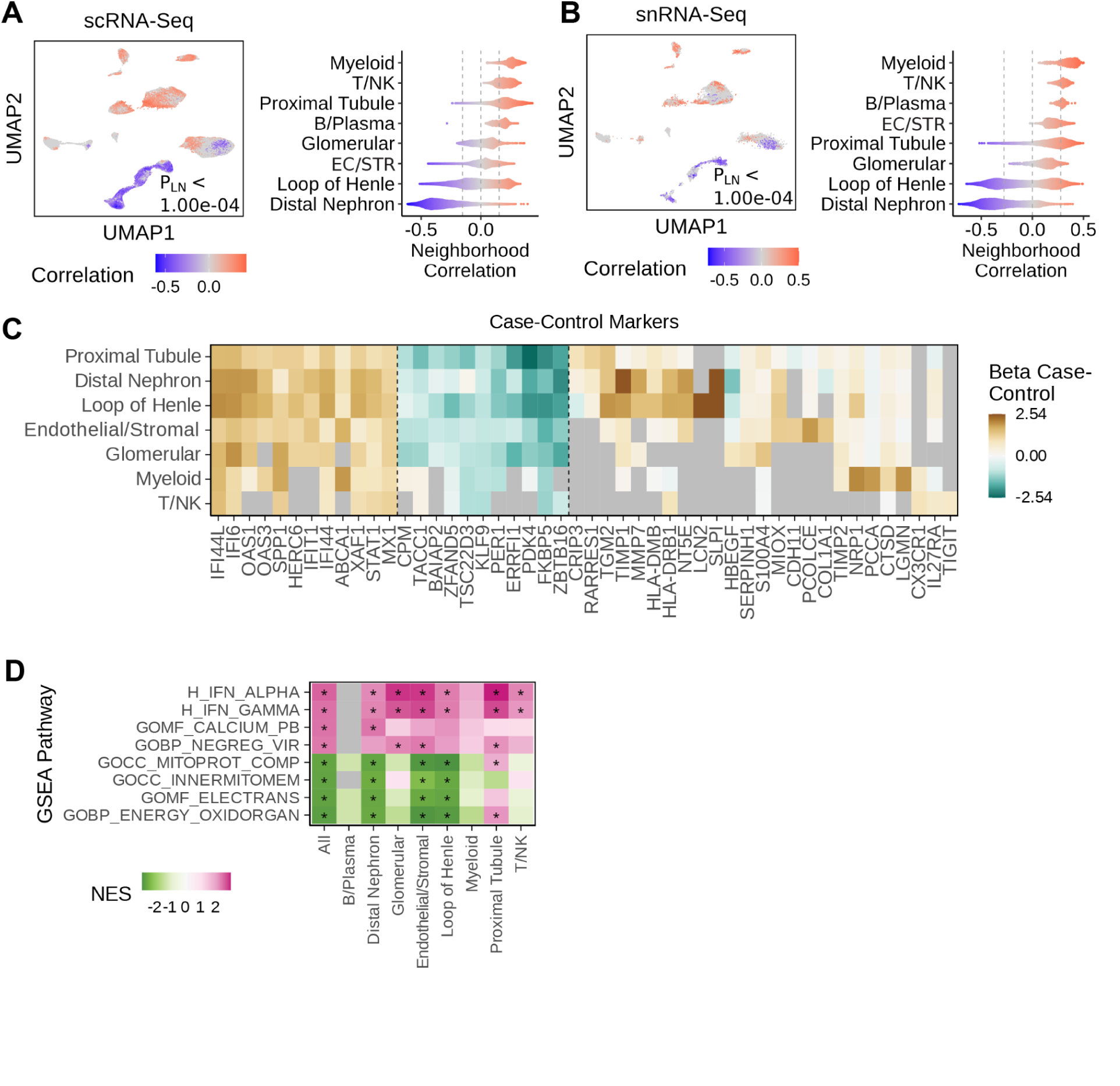
Whole-tissue case-control CNA and differential expression. **(A)** Across all cell types, CNA results for case-control association with no covariate corrections, in scRNA-seq data. Left - UMAP displaying significant per-cell associations with LN, with FDR cutoff of 0.05. Non-significant associations are colored in grey. P-value is the global P-value for cell state associations with LN phenotype. Right - violin plots of clusters containing cells passing FDR significance for LN association. Dashed vertical lines represent the correlation threshold with FDR < 0.05. **(B)** Same as **(A)** in snRNA-seq data. **(C)** For each cell type, differential expression for genes associated with case-control status. Dotted lines separate genes enriched in most cell types and depleted in most cell types. **(D)** Normalized enrichment scores (NES) from pathway enrichment of differential gene expression results, using Hallmark and Ontology pathways. * indicates a Bonferroni-adjusted p-value less than 0.05. Grey color indicates the pathway was not tested due to low expression of pathway genes in that cell type. **(E),** Same as **(D),** where pathway enrichment is on adjusted differential gene expression results, where results have been corrected for all cross-cell-type differential expression signals.

**Fig. S20.**
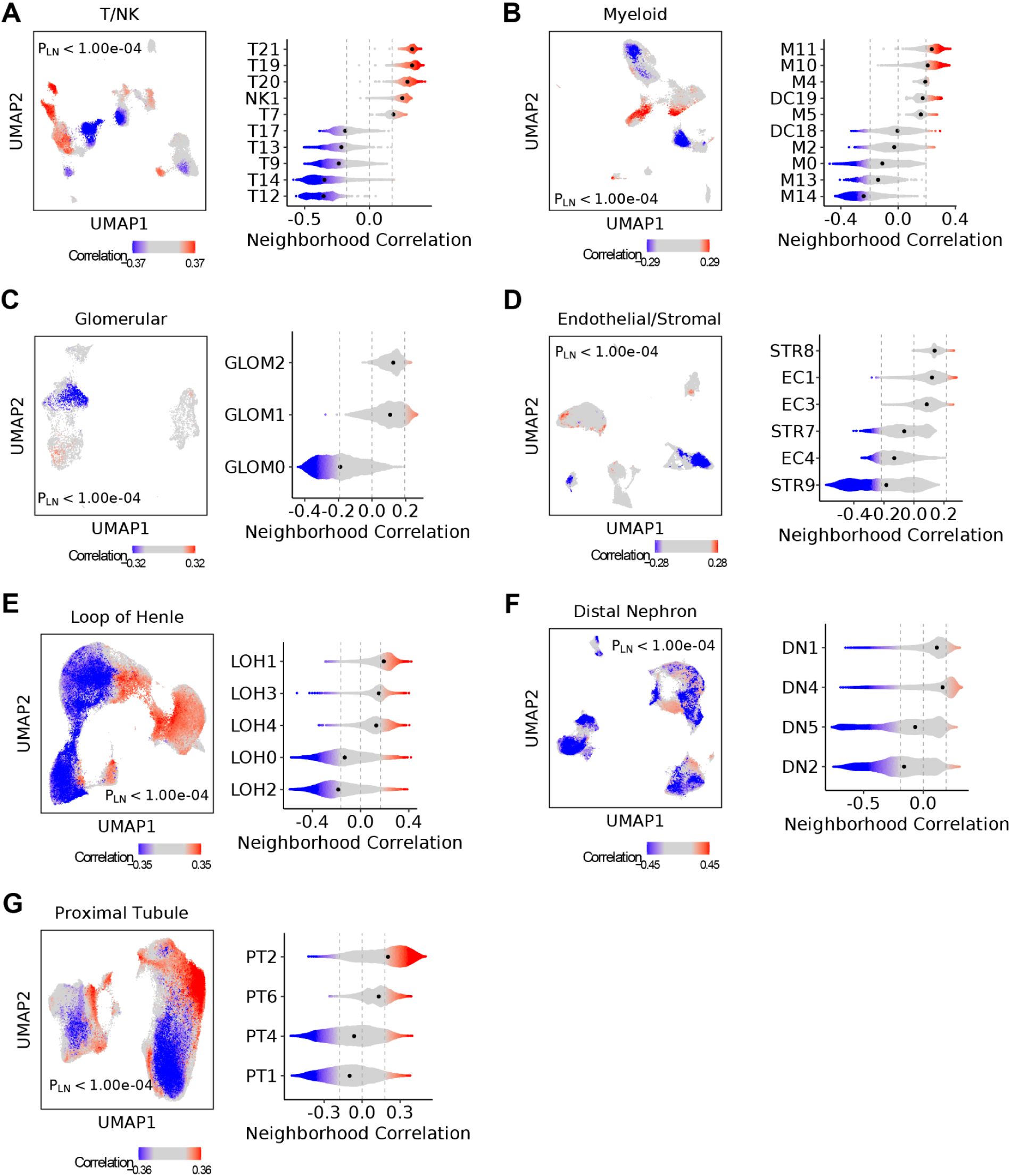
Cell state case-control CNA. **(A)** Within T/NK cells, CNA results for case-control association with covariate corrections for age and sex, in scRNA-seq data. Left - UMAP displaying significant per-cell associations with LN, with FDR cutoff of 0.05. Non-significant associations are colored in grey. P-value is the global P-value for cell state associations with LN phenotype. Right - violin plots of clusters containing cells passing FDR significance for LN association. Dashed vertical lines represent the correlation threshold with FDR < 0.05. **(B)** Same as **(A)** for Myeloid cells. **(C)** Same as (**A)** for B/Plasma cells. **(D)** Same as **(A)** for endothelial/stromal cells. **(E)** Same as **(A)** for loop of Henle cells. **(F)** Same as **(A)** for distal nephron cells. **(G)** Same as **(A)** for proximal tubule cells.

**Fig. S21.**
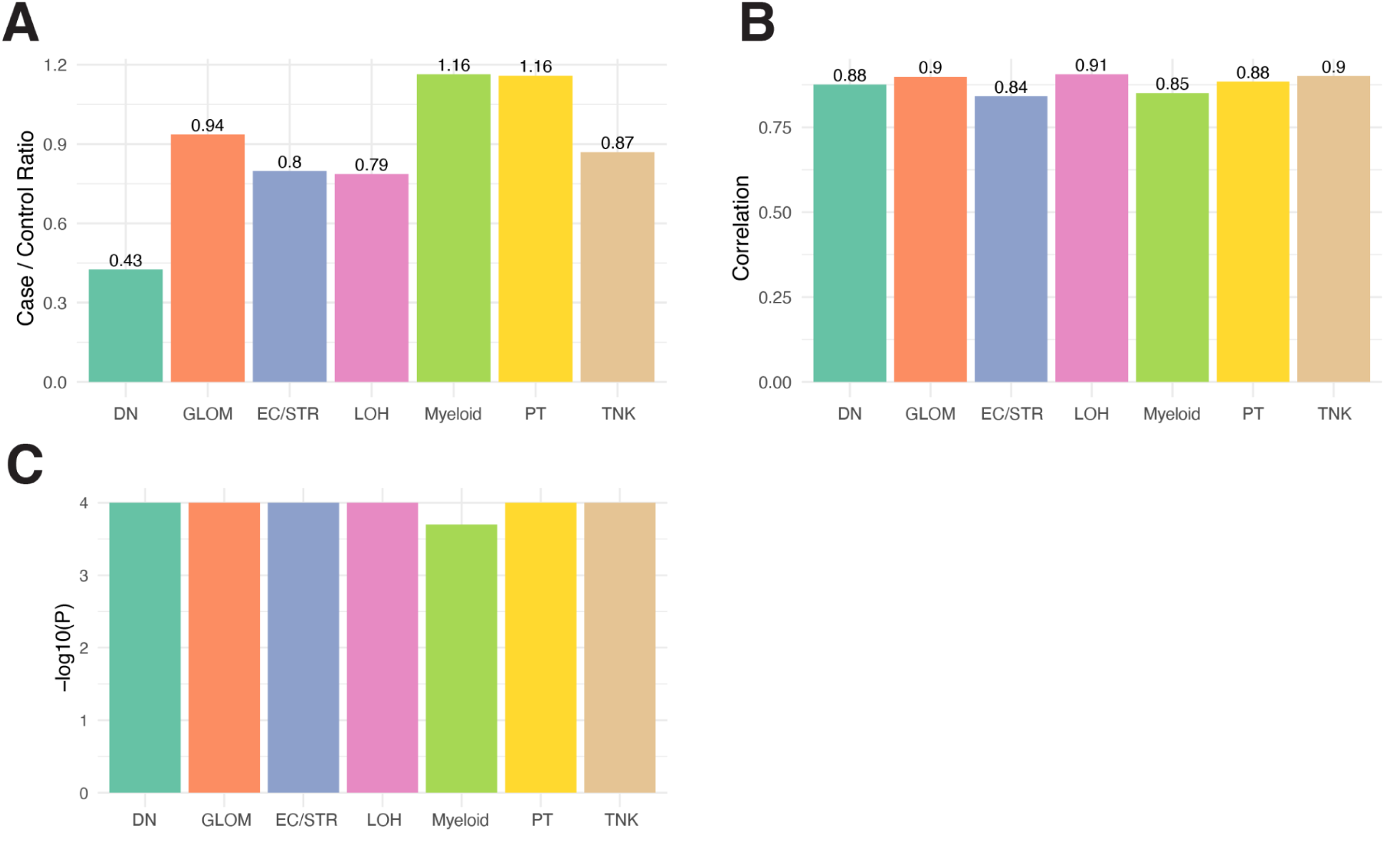
Subsampling of case-control CNA associations. **(A)** Ratio of total case/control cell counts stratified by cell type. **(B)** Neighborhood-level correlations of shared cells between case-control CNA and case-control CNA with case samples subsetted to match control sample count and cell count distributions. These results suggest that similar cell populations are expanded or contracted in both data sets. **(C)** Global P value of subsetted case-control associations within cell type.

**Fig. S22.**
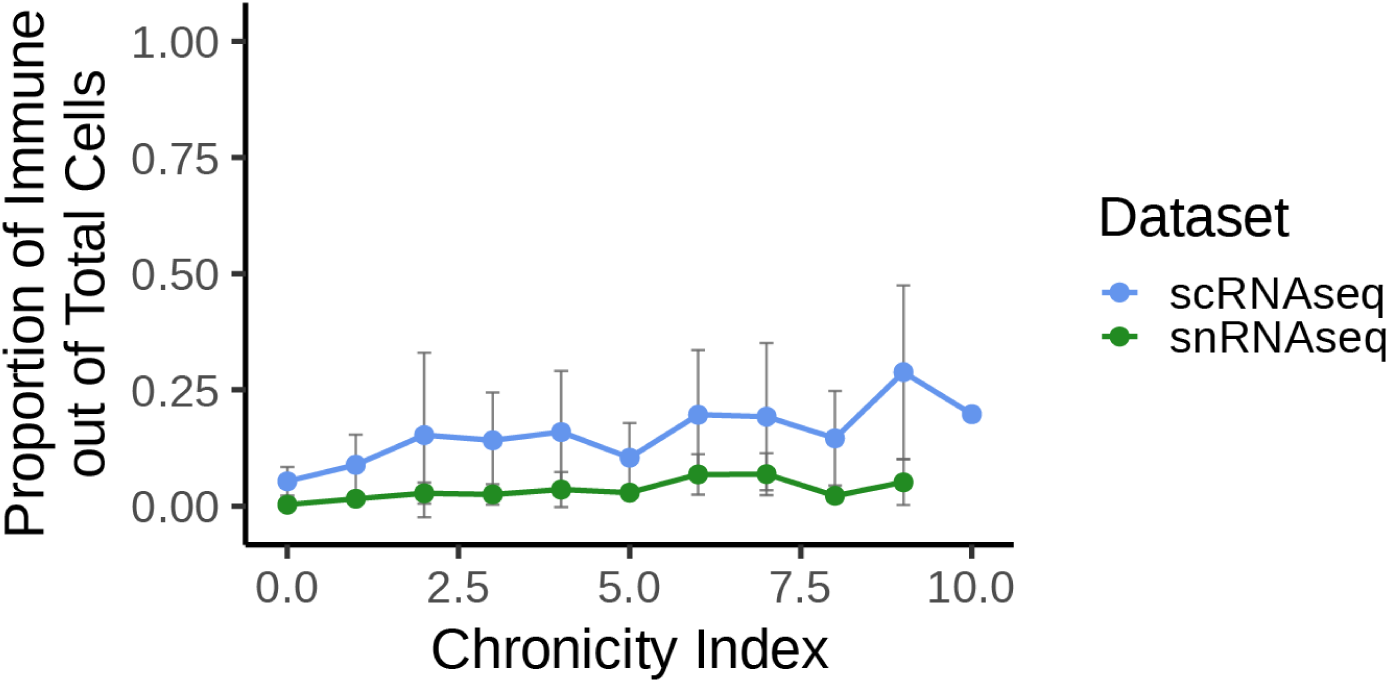
scRNAseq vs snRNAseq proportions. Per-sample proportions of immune cells (T/NK, B/plasma, myeloid) out of all cells in tissue, subset by data modality. Points represent the mean proportion across samples for a given chronicity index value. Error bars represent the standard deviation across samples at that chronicity index value.

**Fig. S23.**
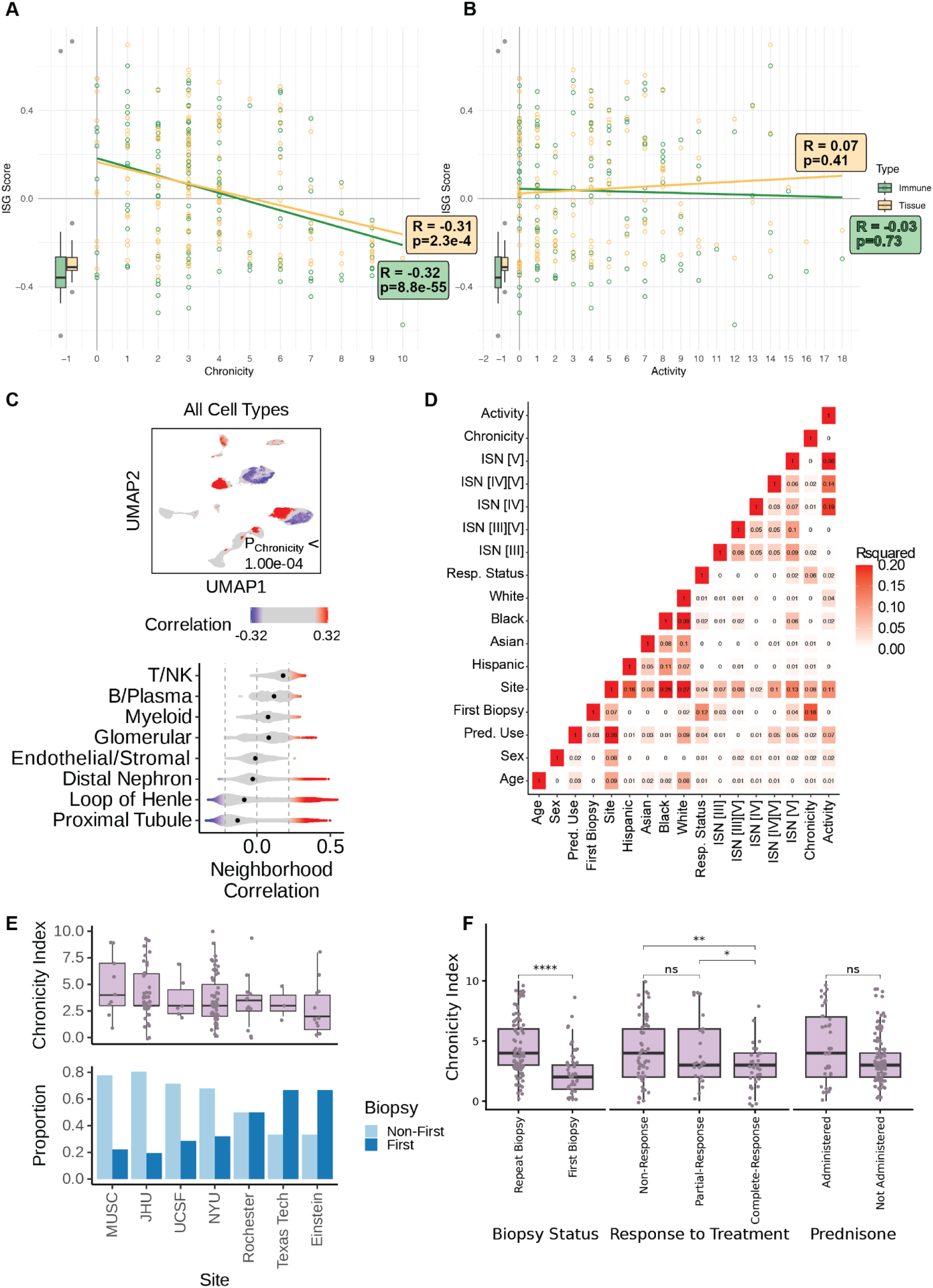
ISG-score and sample-level covariate analysis. **A-B,** Correlation between ISG score (averaged per sample) and Chronicity **(A)**, or Activity Index **(B)**. Controls are displayed as boxplots on the left. Data is stratified between cells originating from immune (green) and tissue (yellow) populations. **(C)** Across all cell types, CNA results for chronicity index association while adjusting for first biopsy and site, in scRNA-seq data. Top - UMAP displaying significant per-cell associations with chronicity index, with FDR cutoff of 0.05. Non-significant associations are colored in grey. P-value is the global P-value for cell state associations with chronicity index phenotype. Bottom - violin plots of clusters containing cells passing FDR significance for chronicity index association. Dashed vertical lines represent the correlation threshold with FDR < 0.05. **(D)** Sample-level correlations of potentially confounding clinical variables. **(E)** Top: Boxplots displaying per-sample chronicity index stratified by patient recruitment sites. Box hinges indicate the first and third quartile of the data, with the center line indicating the median. Whiskers indicate 1.5 x the interquartile range, or the furthest outlier, whichever is less. Bottom: Barplot reflecting the proportion of samples, stratified by site, corresponding to a first biopsy sample or a later (non-first) sample. Color indicates sample first biopsy status. Only sites with a chronicity index measured for more than 1 sample are displayed. **(F)** Boxplots displaying per-sample chronicity index stratified by various relevant clinical categories. Box hinges indicate the first and third quartile of the data, with the center line indicating the median. Whiskers indicate 1.5 x the interquartile range, or the furthest outlier, whichever is less. **** indicates p<0.0001 and ns indicates p>0.05.

**Fig. S24.**
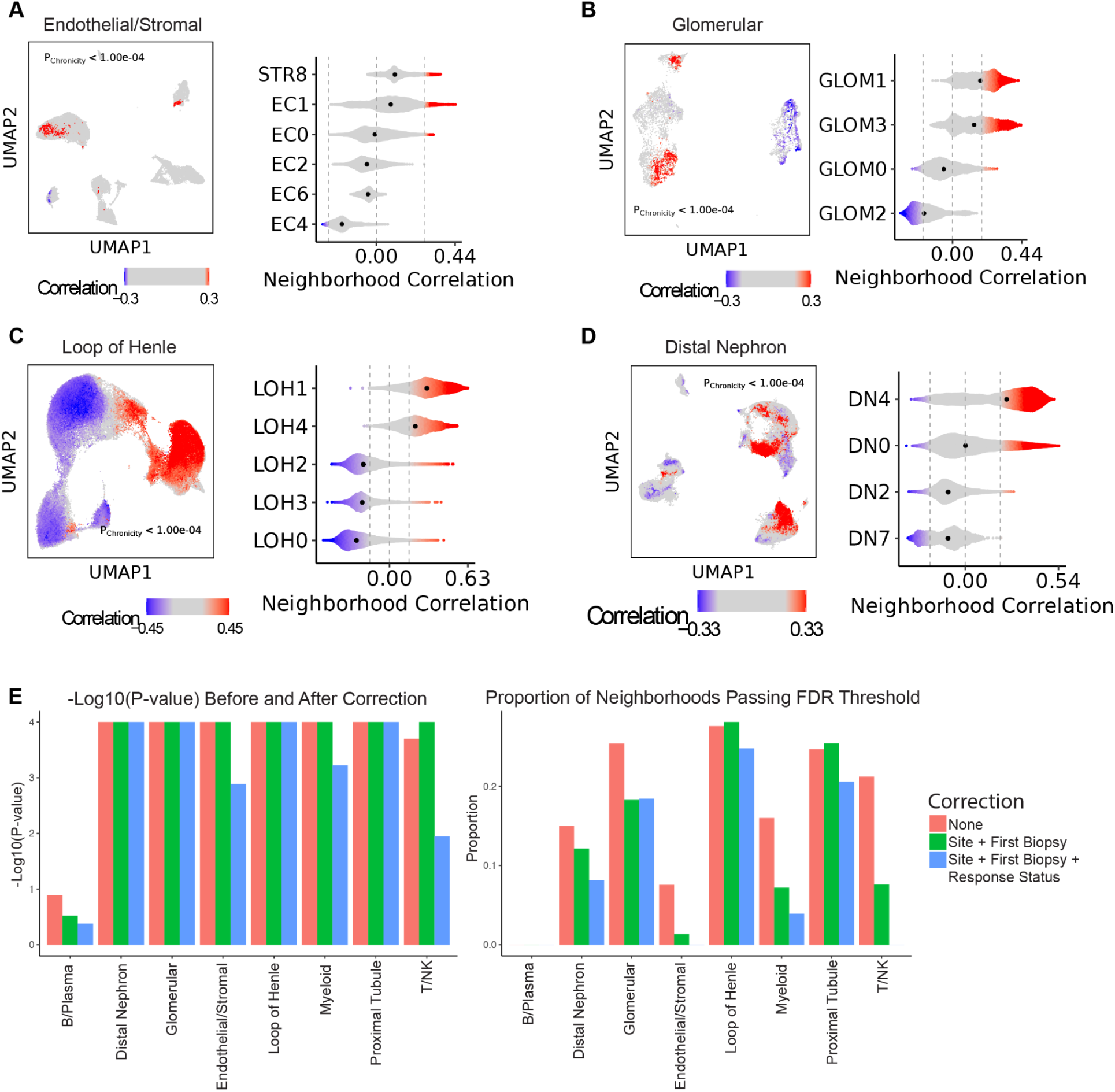
Cell state chronicity CNA analysis. **(A-D)** CNA testing correcting for major collection sites and whether the sample is from the initial biopsy following diagnosis for the endothelial/stromal compartment **(A)**, glomerulus **(B)**, loop of Henle **(C)**, or the distal nephron **(D)**. **(E)** Global p values (left) and the percent of neighborhoods that pass FDR<0.10 (right) for CNA testing of chronicity accounting for either no or additional relevant clinical covariates.

**Fig. S25.**
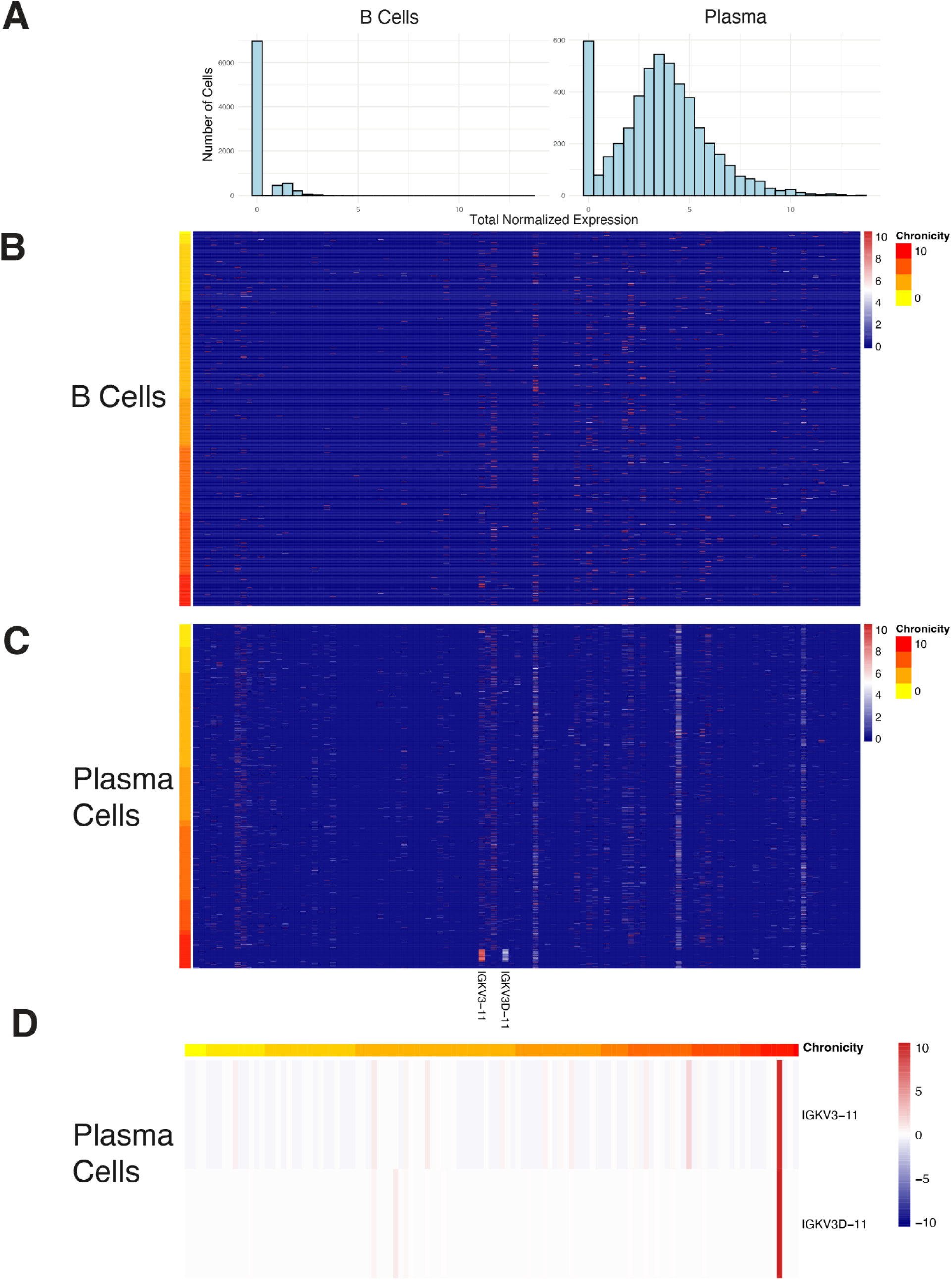
Clonotyping Analysis of Tissue B cells. **(A),** Histogram of cell counts across various levels of total normalized expression of light chain genes stratified by B cells (left) and plasma cells (right). **(B)** Heatmap of light chain expression for each light chain (column) per B cell (row). B cells are ordered from cells originating from low-chronicity samples to high-chronicity samples. **(C)** Same as **(B)**, but for plasma cells. **(D)** Light chain genes IGKV3-11 and IGKV3D-11 pseudobulked across plasma cells per sample.

**Fig. S26.**
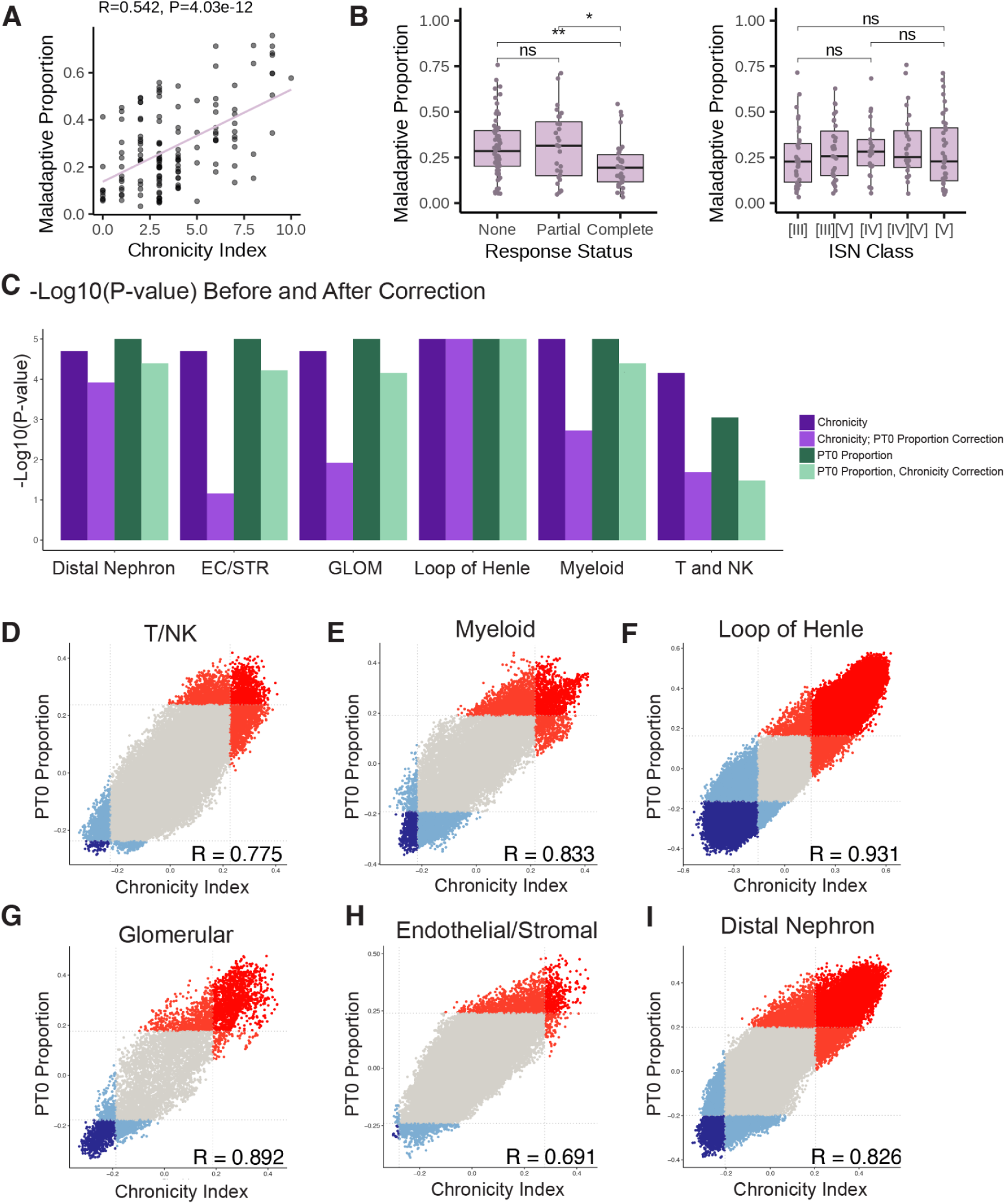
PT0-chronicity analysis. **(A)** Scatterplot of the per-sample correlation of the proportion of maladaptive proximal tubule cells and chronicity index. **(B)** Boxplots indicating differences of the proportion of maladaptive proximal tubule cells between various response statuses (left) and ISN class (right). * indicates p<0.05 and ** indicates p<0.01. “ns” indicates p>0.05. (**C)** Global p values of CNA testing for either chronicity or maladaptive proximal tubule cell proportion, correcting for either no additional covariate or the opposite measure. **D-I**, Scatterplot of per-neighborhood correlations of CNA testing of Chronicity index (x-axis) and proportion of maladaptive proximal tubule cells (y-axis) for cells from either the T and NK **(D)**, myeloid **(E)**, loop of Henle **(F)**, glomeruli **(G)**, endothelial/stromal **(H)**, or distal nephron **(I)** compartments.

**Fig. S27.**
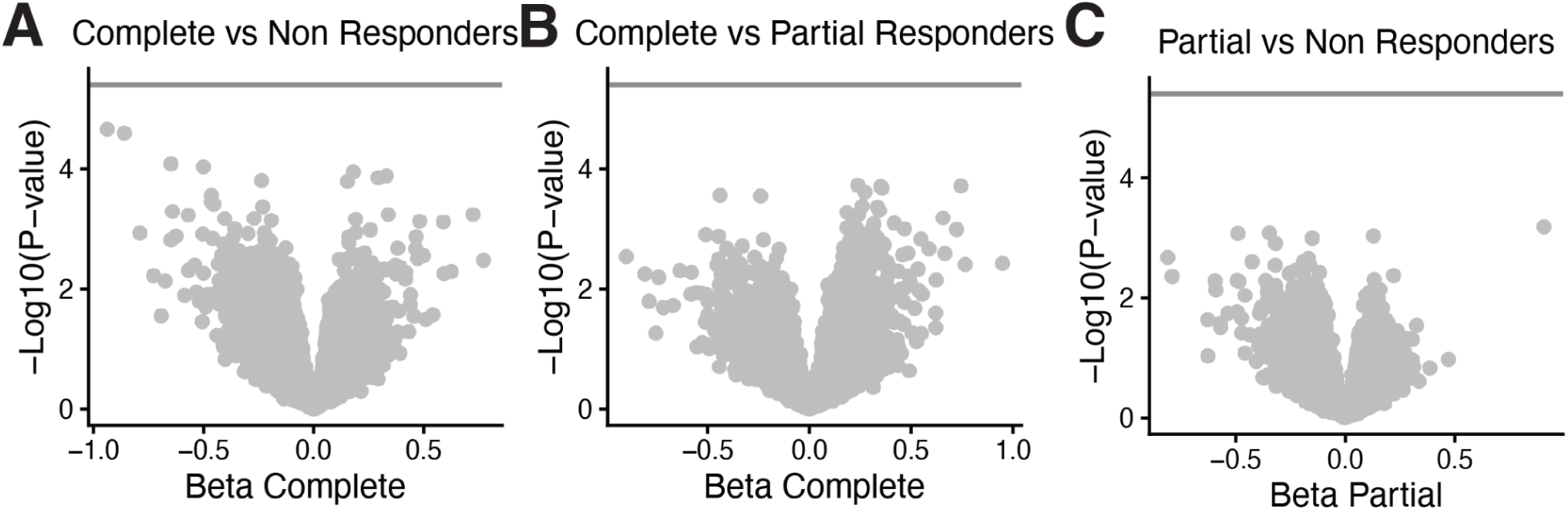
Differential gene expression of PT0 between treatment response groups. Volcano plots of differentially expressed genes within PT0 between treatment groups for complete responders vs non responders **(A)**, complete vs partial responders **(B)**, and partial vs non responders **(C).**

**Fig. S28.**
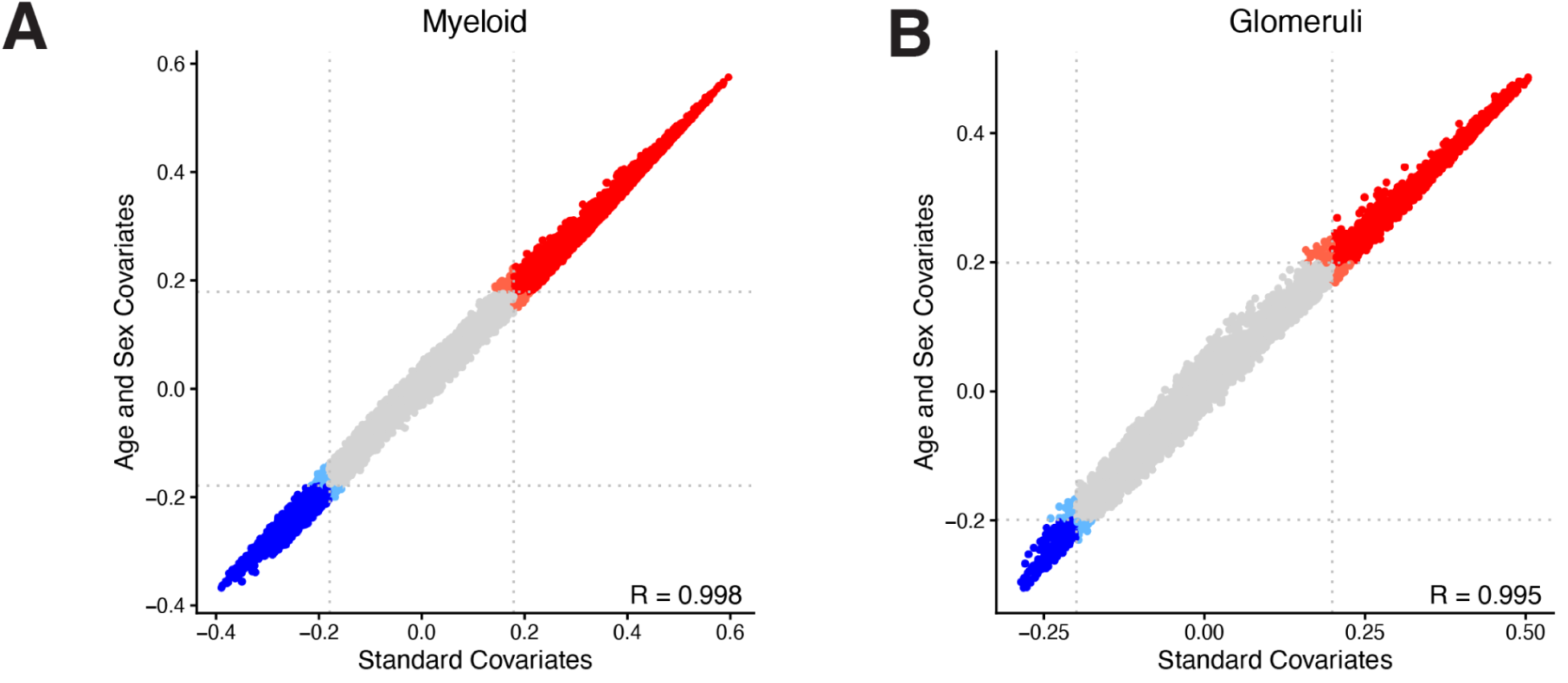
Concordances between standard covariates and additional age and sex covariates in activity index CNA. **(A, B)** Correlations of activity index CNA with standard covariates compared to additional age and sex covariates in myeloid **(A)** and glomerular **(B)** cell types. Horizontal lines are drawn at each individual analyses FDR < 0.10 threshold.

**Fig. S29.**
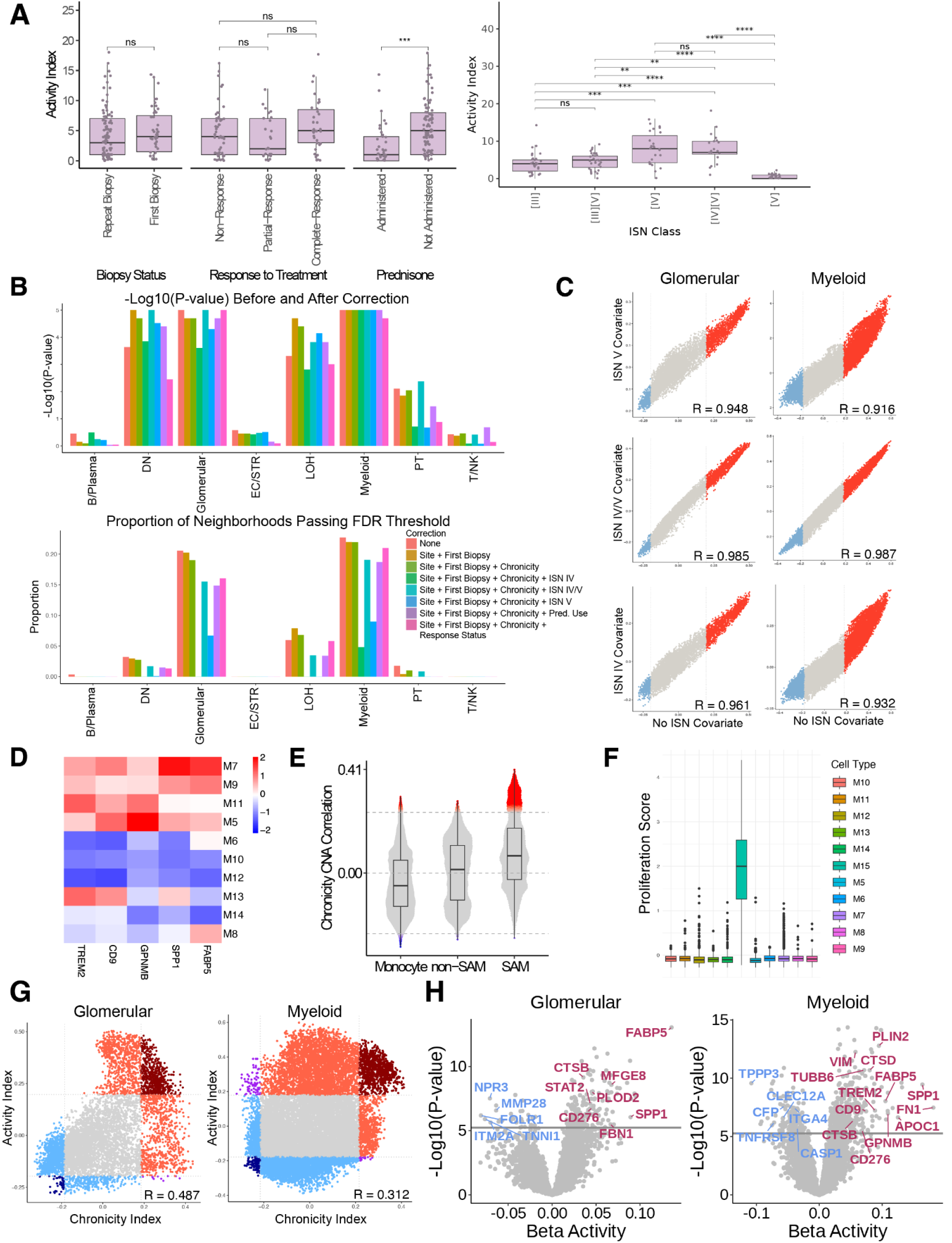
Activity CNA analysis. **(A)** Boxplots indicating differences in sample-level differences in Activity Index for relevant clinical variables (left) and ISN class (right). **(B)** global p values (top) and the percent of neighborhoods that pass FDR<0.1 (bottom) for CNA testing of activity accounting for either no or additional relevant clinical covariates. **(C)** Scatterplot of per-neighborhood correlations for glomeruli and myeloid populations between CNA testing of activity (x-axis) and ISN class IV, IV/V, or V (y-axis). **(D)** Gene expression for SAM markers in tissue macrophage populations. **(E)** Violin plot of Chronicity CNA correlations for monocyte, non-SAM, and SAM populations. Significant neighborhoods are colored red and blue for expanded and contracted populations, respectively. **(F)** Box and whisker plots of proliferation scores stratified to macrophage cell populations and proliferating myeloid cells **(G)** Scatterplot of per-neighborhood correlations for glomeruli and myeloid populations between CNA testing of chronicity index (x-axis) and activity index (y-axis) for glomerular (left) and myeloid (right) cells. **(H)** In glomerular (left) and myeloid (right) cells, differential expression results for genes associated with activity index. Betas and p-values are derived from pseudobulk linear modeling. Genes upregulated with respect to activity index are displayed in red, downregulated genes are displayed in blue. Horizontal lines indicate the Bonferonni corrected significance threshold for p<0.05.

**Fig. S30.**
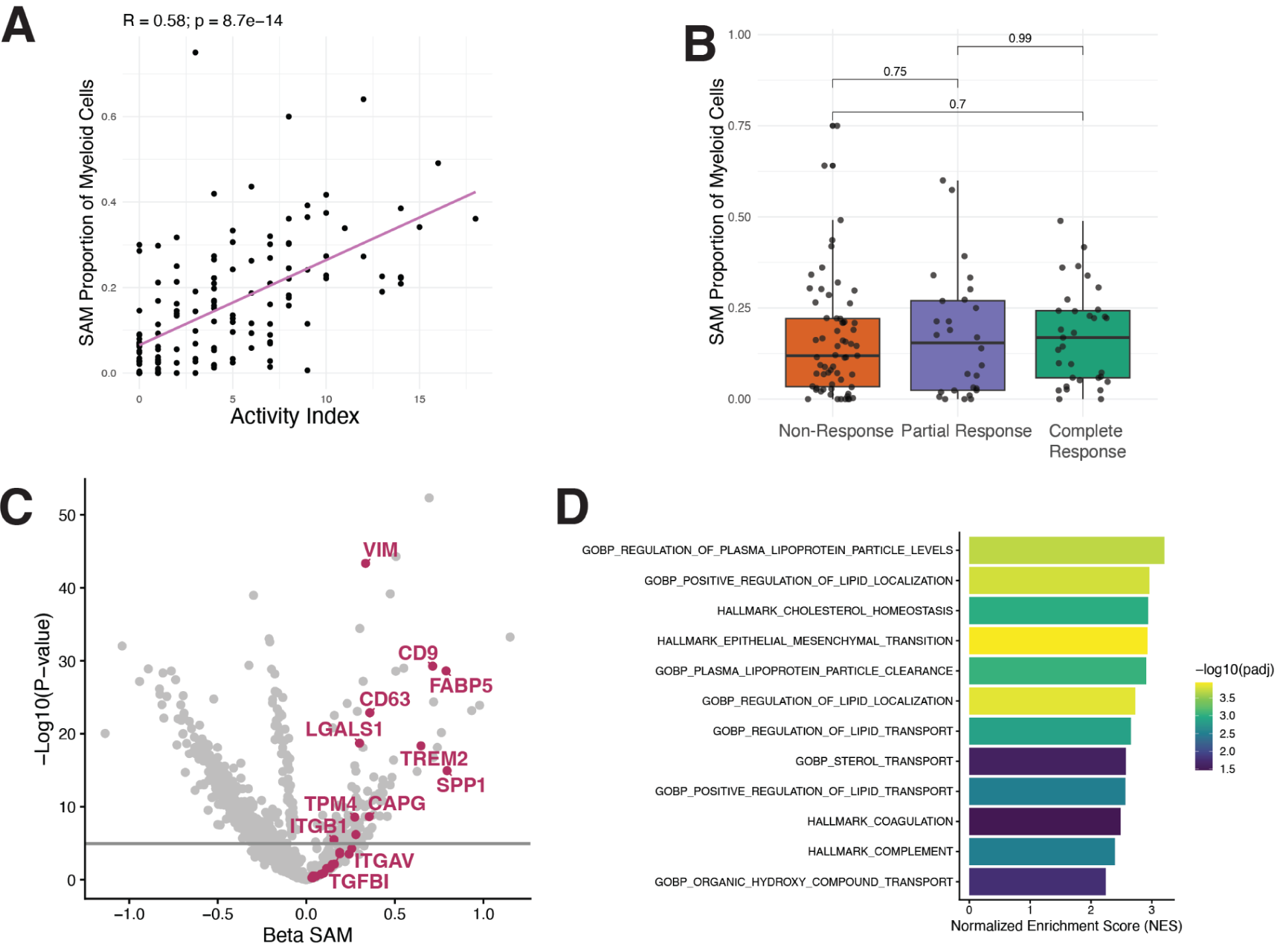
Differential gene expression of SAMs over other myeloid cells. **(A)** Correlation plot of activity and per-sample SAM proportion of myeloid cells. **(B)** Boxplot of per-sample SAM proportion stratified by treatment response. **(C)** Volcano plot of SAM vs nonSAM myeloid cell differential gene expression. Horizontal line drawn at -log10(p) = 0.05/4,455 genes tested. **(D)** GSEA bar plot indicating Normalized Enrichment Score (NES) and -log10(Bonferonni-corrected p value).

**Fig. S31.**
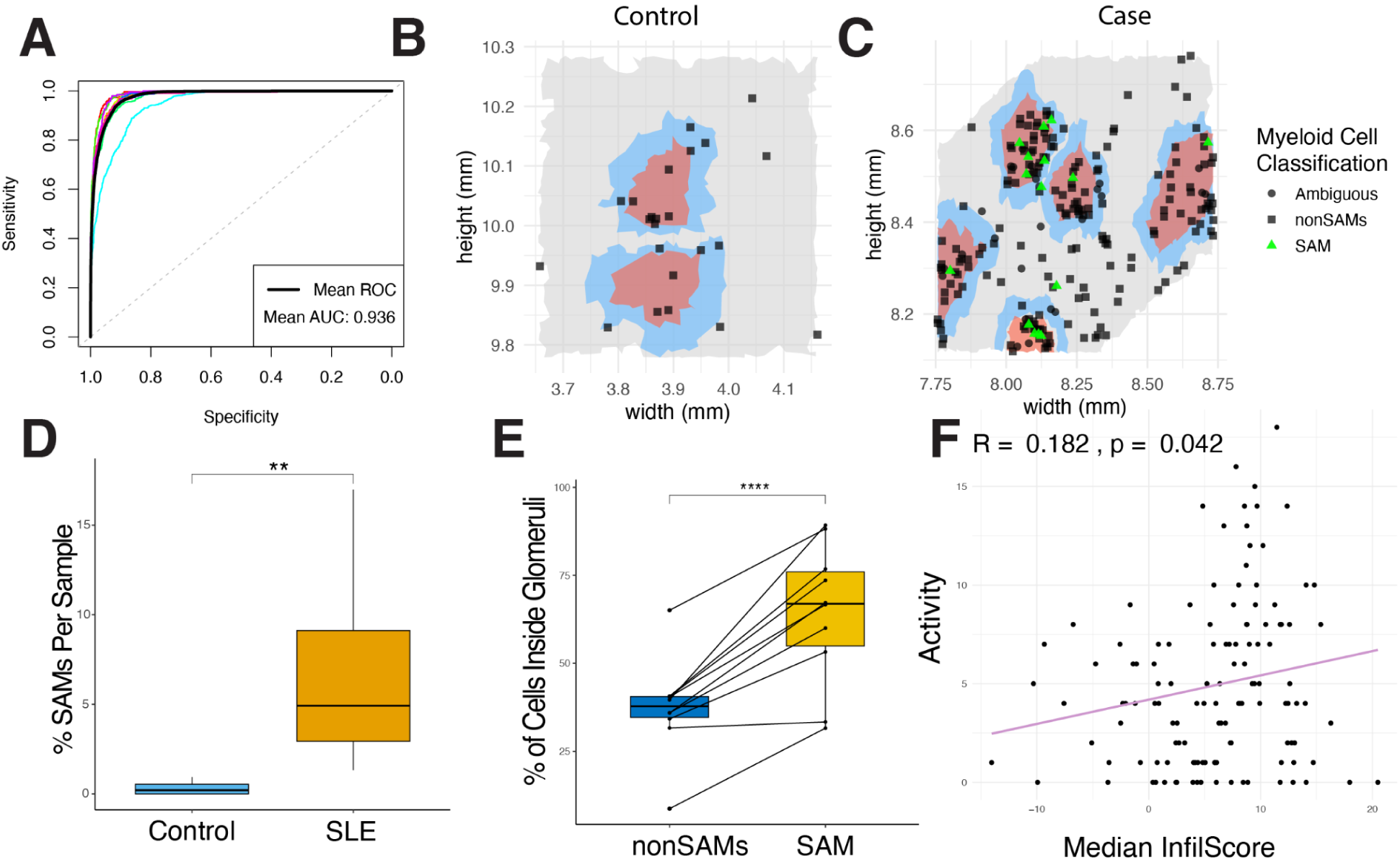
Integration of Myeloid Classifier with Spatial Data. **(A)** Average AUC of myeloid cell classifier. **(B, C)** Non-myeloid cells colored by spatial niche and myeloid cells distinguished by shape and colored by SAM marker FABP5 for a select control **(B)** and case **(C)** FOV. **(D)** Boxplot of percent SAMs per sample in control and SLE cases in spatial data. **(E)** Boxplot of percent of myeloid cells in case samples inside glomerular niches. **(F)** Per-sample scatterplot of median sample infiltration score versus clinical activity index.

**Fig. S32.**
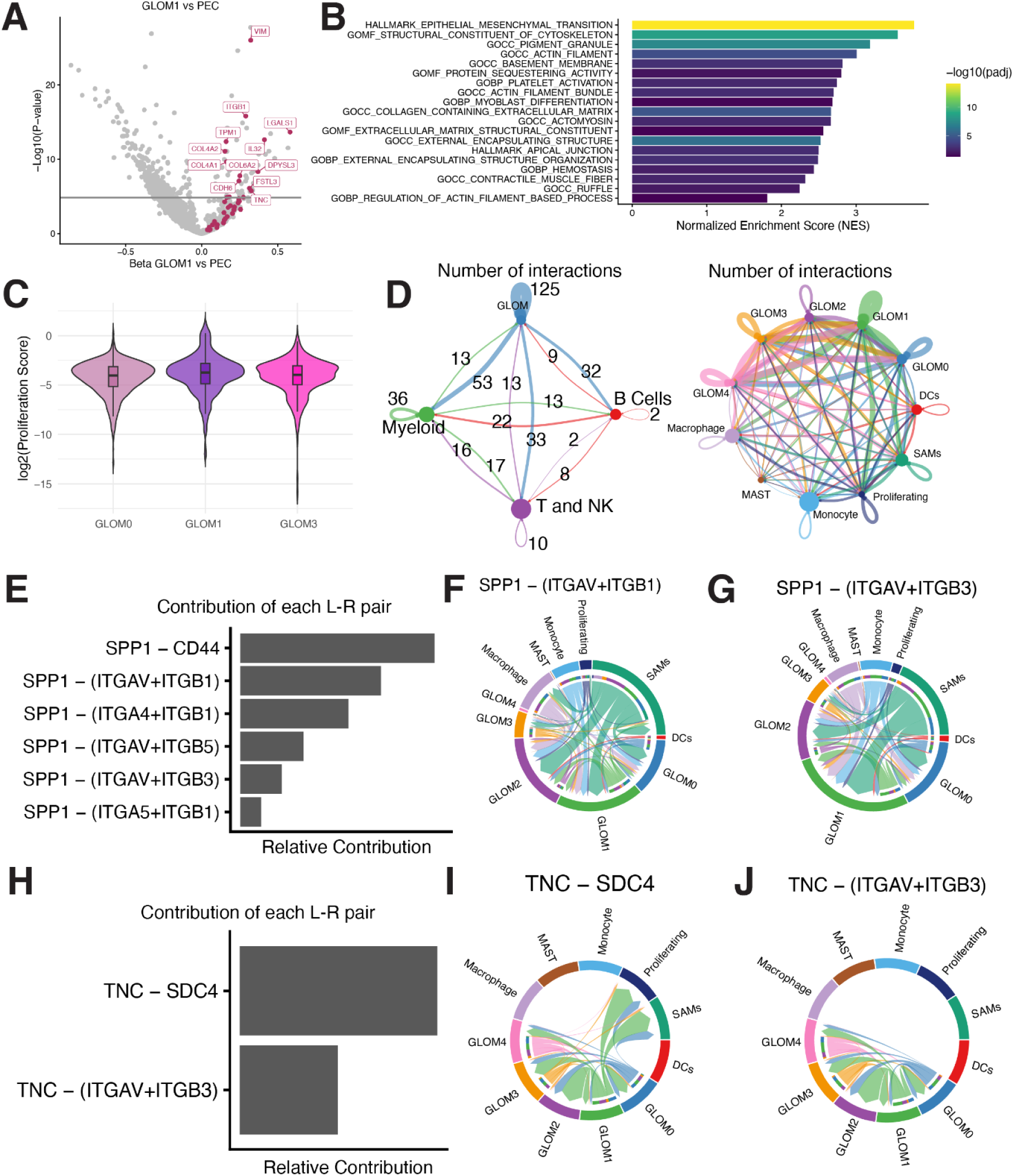
GLOM1 Gene Signatures. **(A)** Volcano plot of differentially expressed genes between GLOM1 and other PEC cells. Epithelial-mesenchymal transition genes are highlighted in red. A horizontal line is drawn at p < 0.05 after Bonferroni correction for genes tested. **(B)** Top significantly upregulated GSEA enrichment of differentially expressed genes in GLOM1 compared to other PEC populations. **(C)** Violin plot of log2 of infiltration score stratified by cell type. **(D)** Receptor-ligand analysis between glomeruli and broad immune tissue cell types (left) and subcategorized macrophage populations (left). **(E)** Relative contributions of ligand-receptor pairs of the SPP1 pathway. **(F-G)** Chord plot of select ligand-receptor pairings in the SPP1 pathway grouped by cell state for SPP1 and ITGAV+ITVB1 heterodimer (left) and ITGAV+ITGB3 heterodimer (right). **(H)** Relative contributions of ligand-receptor pairs of the Tenasin pathway. **(I-J)** Chord plot of select ligand-receptor pairings in the SPP1 pathway grouped by cell state for SPP1 and SDC4 (left) and ITGAV+ITGB3 heterodimer (right).

**Fig. S33.**
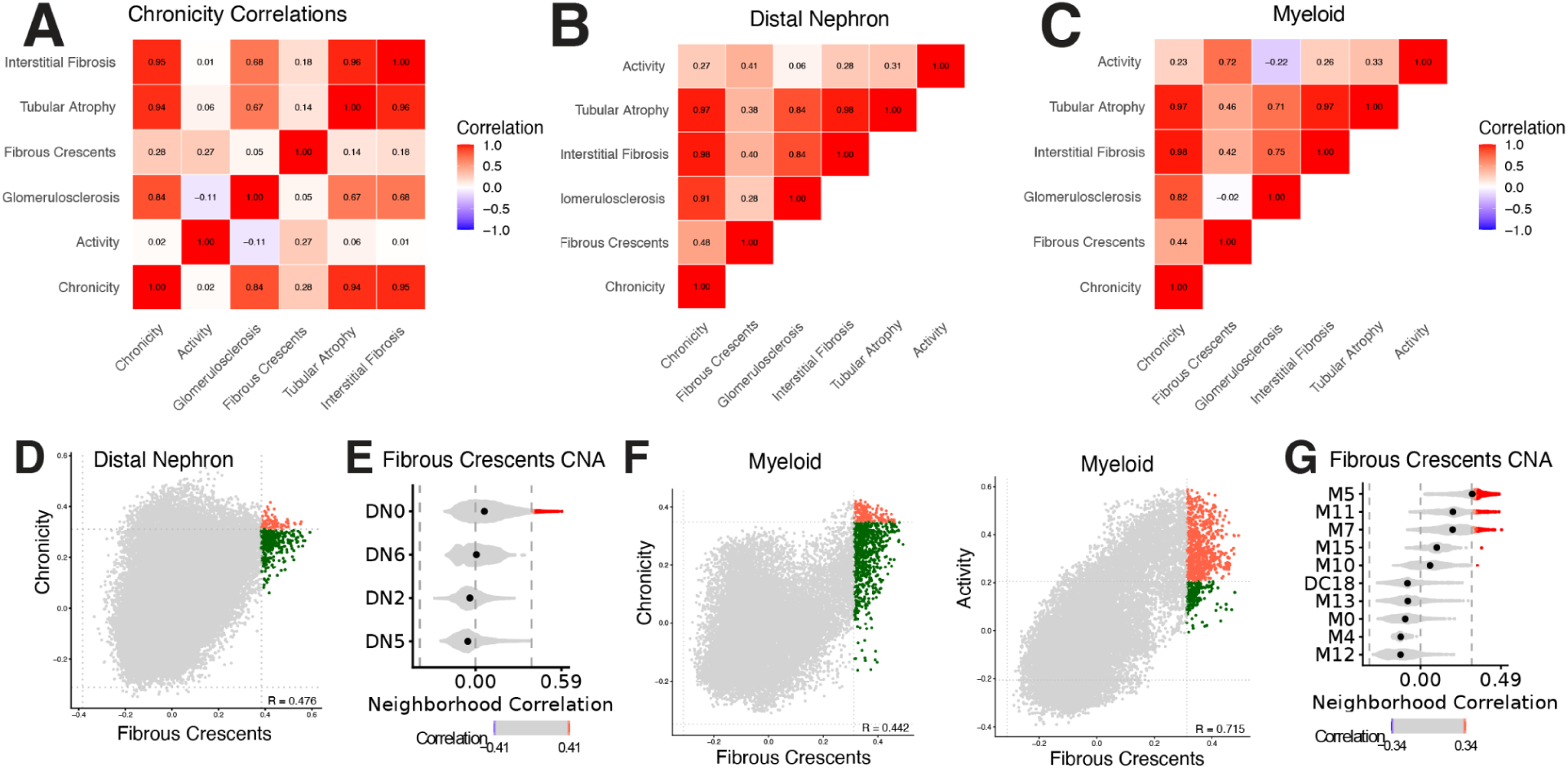
CNA with Chronicity Index Components. **(A)** Sample level correlations for chronicity components. **(B-C)** Neighborhood-level correlations for each chronicity component in distal nephron **(B)** and myeloid **(C)** cells. **(D)** Neighborhood-level correlation scatterplot for chronicity and fibrous crescent CNA. Grey lines are added for each test’s respective FDR threshold. Neighborhoods concordantly expanded in both CNA and fibrous crescent CNA are colored red, and novel neighborhoods significantly expanded in fibrous crescents but not in chronicity are colored in green. **(E)** Violin plot of neighborhood correlations of fibrous crescent CNA in distal nephron cells. Dashed gray line added at FDR < 10%. **(F)** Same as **(D)** for myeloid cells. **(G)** Same as **(E)** for myeloid cells. Neighborhoods that are significant in the subindex but not overall activity are highlighted in green.

**Fig. S34.**
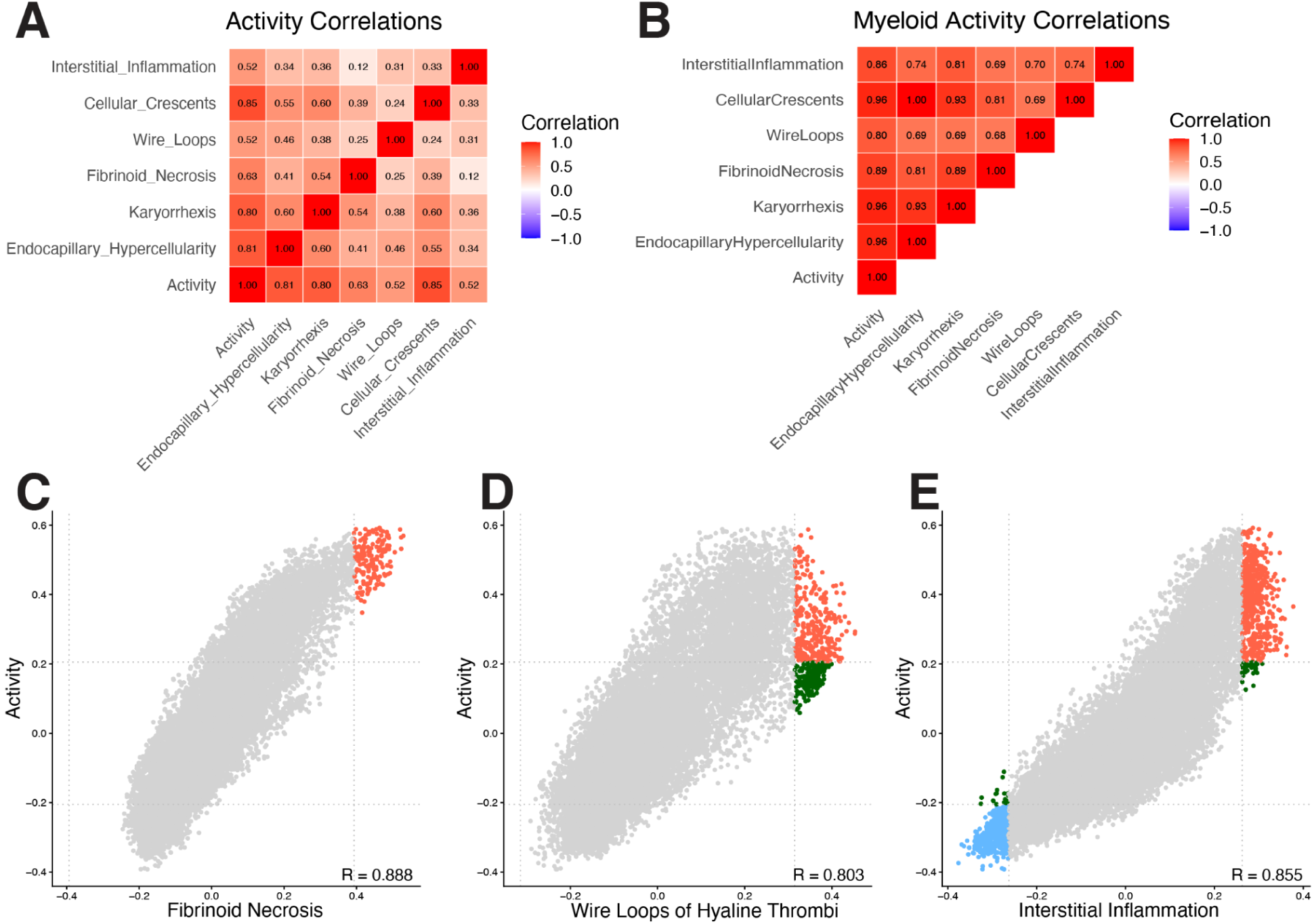
CNA with Activity Index Components. **(A)** Sample level correlations for activity components. **(B)** Neighborhood-level correlations for each activity component in myeloid cells. **(C-E)** Neighborhood-level correlation scatterplot lines between activity CNA and fibrinoid necrosis **(C),** wire **l**oops or hyaline thrombi **(D)**, and interstitial inflammation **(E)**. Dashed gray lines are added for each test’s respective FDR threshold. New neighborhoods that are significant in subindex CNA but not activity CNA are highlighted in green. Neighborhoods that are concordantly significantly expanded or depleted in both are colored red and blue, respectively.

**Fig. S35.**
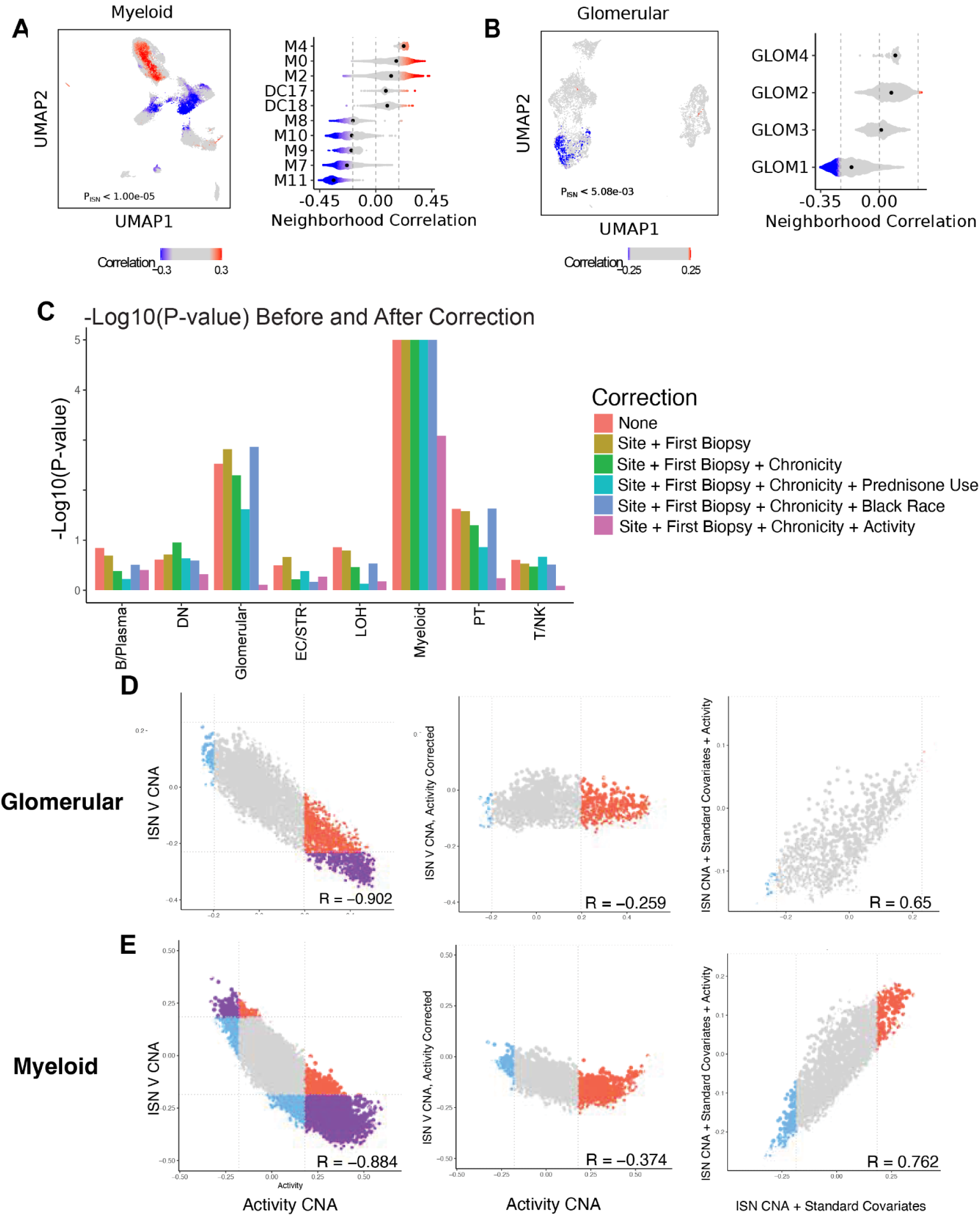
Activity CNA and ISN V CNA analysis. **(A-B)** CNA testing for associations with ISN class V, correcting for site, first biopsy, and chronicity index for myeloid **(A)** and glomerular **(B)** compartments. Left - UMAP displaying significant per-cell associations with ISN Class V, with FDR cutoff of 0.1. Non-significant associations are colored in grey. P-value is the global P-value for cell state associations with LN phenotype. Right - violin plots of clusters containing cells passing FDR significance for ISN class V association. Dashed vertical lines represent the correlation threshold with FDR < 0.1. **(C)** Global p values for CNA testing of ISN class V accounting for either no or additional relevant clinical covariates. **(D-E),** Scatterplots showing per-neighborhood correlations between activity index CNA and either ISN class V CNA either without (left column) or with (middle column) correcting for activity index. Additional scatterplots plot the correlation between ISN class V CNA without correcting for activity (x-axis) or with correction (y-axis). These plots were done for both the glomeruli **(D)** and myeloid **(E)** compartments. All CNA for **(D)** and **(E)** include major collection site, first biopsy status, and chronicity as covariates, along with any other specified covariates.

**Fig. S36.**
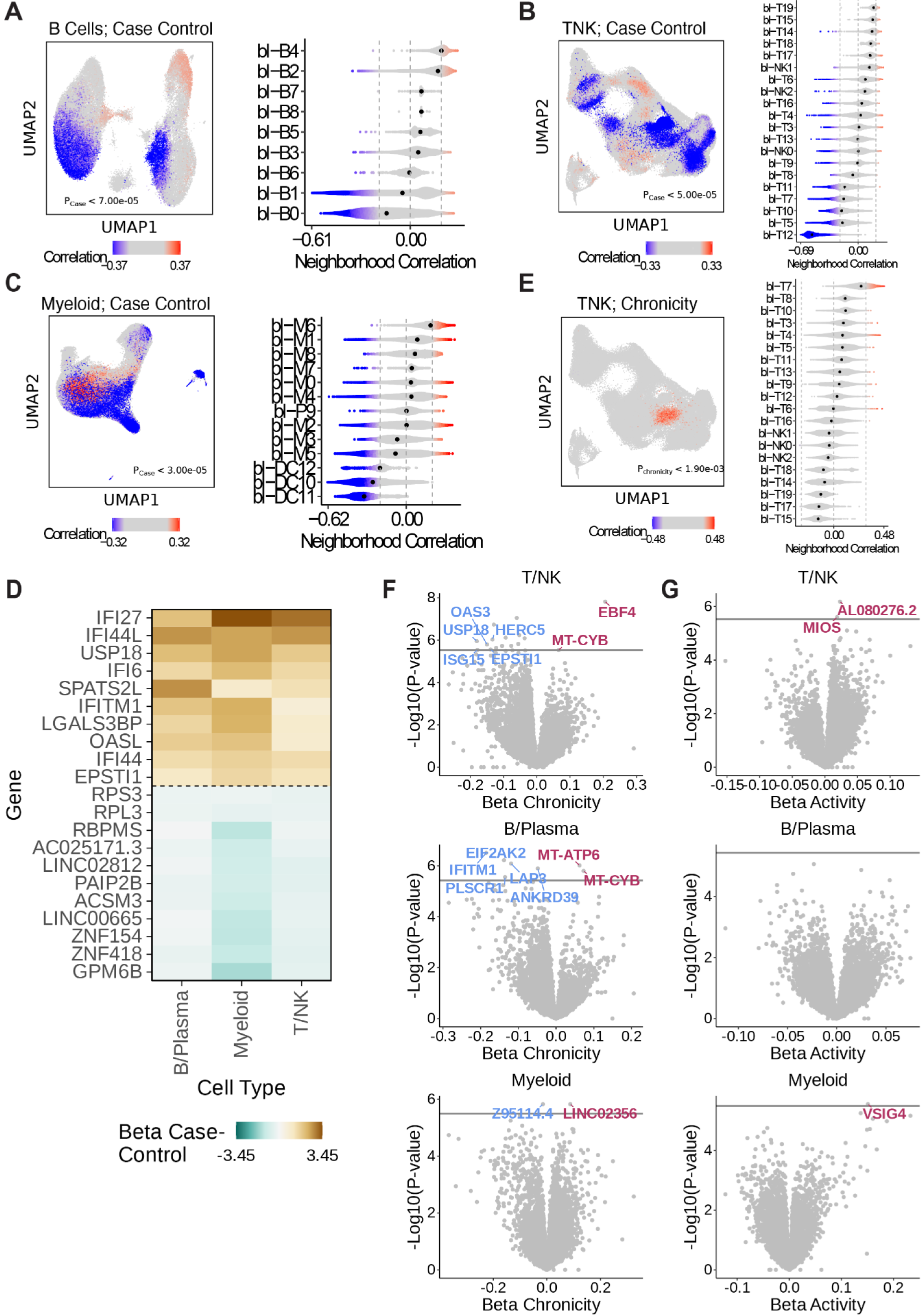
PBMC CNA analysis. **(A-C)** CNA testing for case-control status for B **(A)**, T and NK **(B)**, and Myeloid **(C)** cells, accounting for age and sex as potential covariates. Left - UMAP displaying significant per-cell associations. Non-significant associations are colored in grey. P-value is the global P-value for cell state associations. Right - violin plots of clusters containing cells passing FDR thresholding, indicated by the vertical lines. **(D)** For each cell type, differential expression for genes associated with case-control status in blood. Dotted lines separate genes enriched in most cell types and depleted in most cell types. **(E)** CNA testing for chronicity index in the T and NK compartment, accounting for site and first biopsy site as covariates. **(F-G),** Differentially expressed genes for each cell type with respect to chronicity index and activity index **(G)**. Genes upregulated with respect to each index are displayed in red, downregulated genes are displayed in blue. Horizontal lines indicate the Bonferonni corrected significance threshold for p < 0.05. The top 5 most significant genes in each direction are labeled for each cell type.

**Fig. S37.**
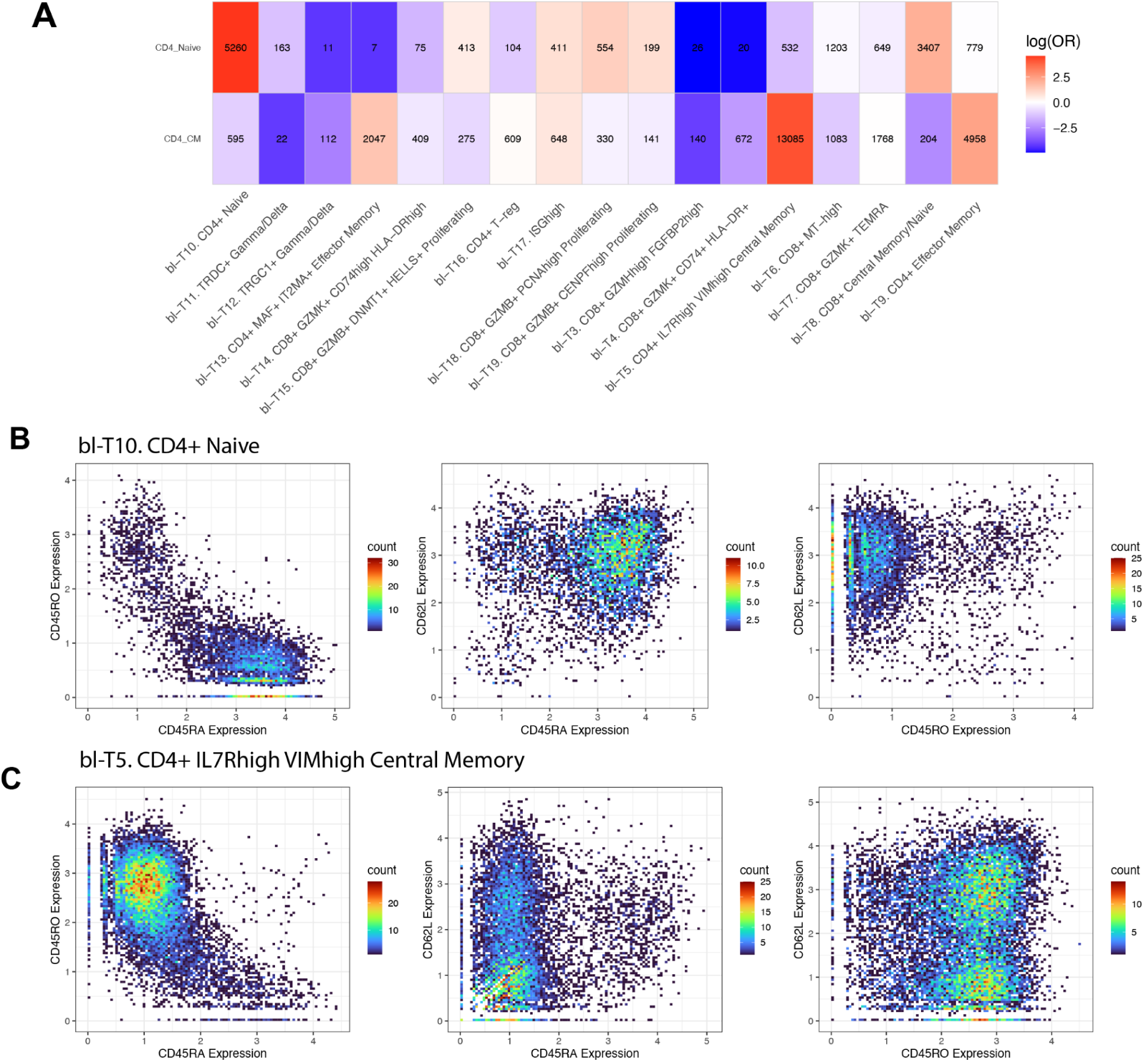
CITE-Seq of PBMC T cells. **(A)** log odds ratio heatmap of TCAT multinomial classification on PBMC T cells. Boxes are labeled with the number of cells assigned by TCAT to that label. **(B-E)** Normalized expression of CD45RA **(B, D)** and CD45RO **(C, E)** protein on all the T cells **(B, C)** and the bl-T10 cluster **(D, E)**. **(F)** Scatterplot of CD45RA protein and CD45RO protein expression within bl-T10.

**Fig. S38.**
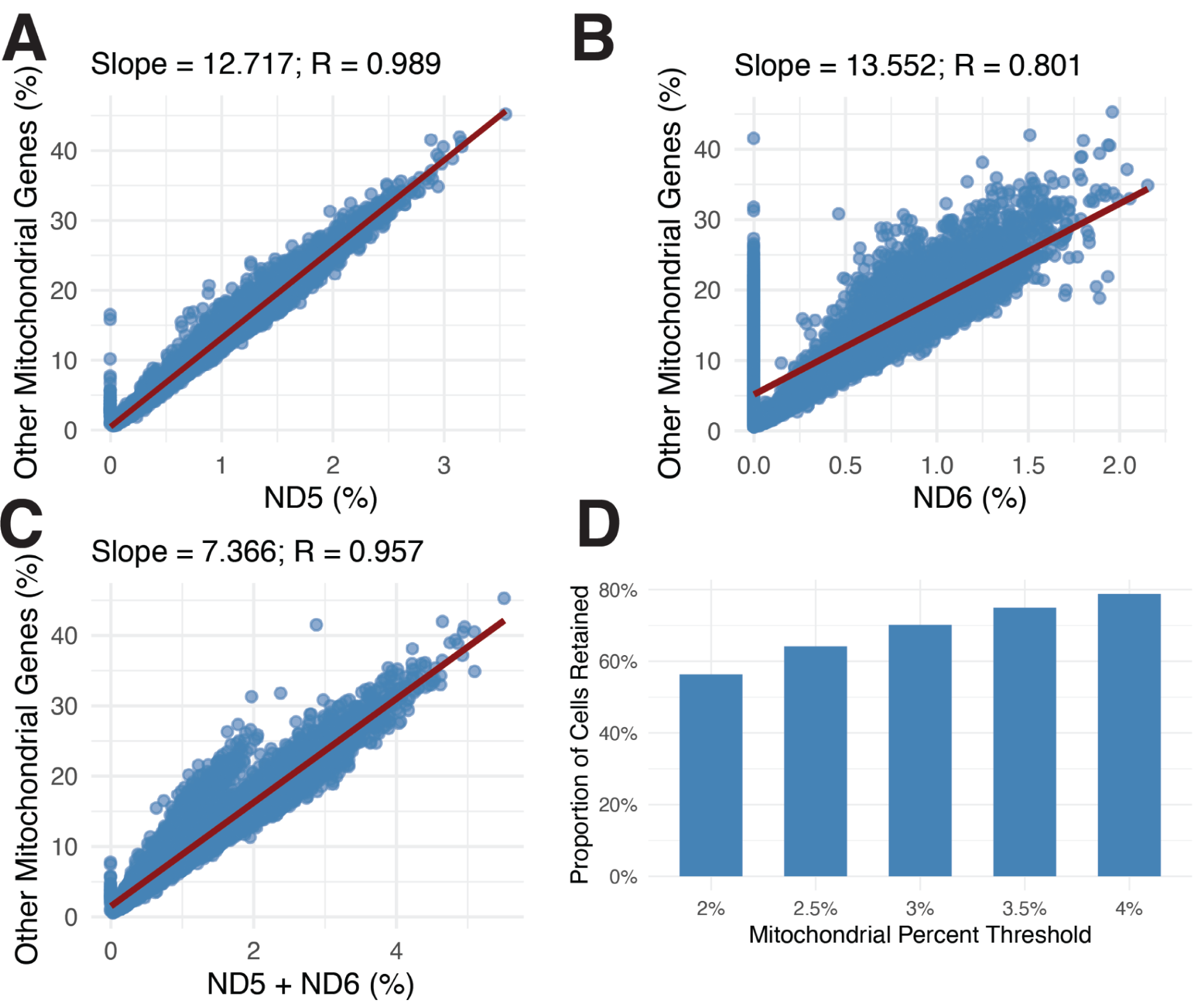
DASH Mitochondrial quality control. **(A-B)** Correlational plots of percentage of mitochondrial genes for MT-ND5 **(A)**, MT-ND6 **(B)**, and MT-ND5 + MT-ND6 against total mitochondrial content **(C)**. **(D)** Barplots of proportion of cells retained in AMP dataset at various MT-ND5 and MT-ND6 cutoffs.

## Supplementary Tables

**Table S1.** Cohort demographics

**Table S2.** Cell and sample counts for data modalities

**Table S3.** Cell counts before and after quality control

**Table S4.** Tissue cell state statistics

**Table S5.** Chronicity differential gene expression results

**Table S6.** CNA cell state correlations with chronicity index

**Table S7.** Sample-level correlations with chronicity index

**Table S8.** Chronicity CNA p-values in tissue

**Table S9.** Broad cell type Chronicity CNA FDR thresholding

**Table S10.** Activity differential gene expression results

**Table S11.** Sample-level correlations with activity index

**Table S12.** Chronicity subcomponent CNA p-values in tissue

**Table S13.** Activity subcomponent CNA p-values in tissue

**Table S14.** Donor counts for blood CNA analysis

**Table S15.** P values for blood CNA analysis

**Table S16.** DASH Sequences

